# Decoding the variant-to-function relationship for *LIPA*, a risk locus for coronary artery disease

**DOI:** 10.1101/2022.11.12.516293

**Authors:** Fang Li, Elise Flynn, Philip Ha, Mazal N. Zebak, Haoxiang Cheng, Chenyi Xue, Jianting Shi, Xun Wu, Ziyi Wang, Yujiao Meng, Jian Cui, Yizhou Zhu, Annie Rozenblyum, Jeana Chun, Antonio Hernandez-Ono, Ali Javaheri, Babak Razani, Marit Westerterp, Robert C Bauer, Yousin Suh, Ke Hao, Tuuli Lappalainen, Hanrui Zhang

**Affiliations:** Cardiometabolic Genomics Program, Division of Cardiology, Department of Medicine, Columbia University Irving Medical Center, New York, NY, USA; New York Genome Center, New York, NY, USA; Department of Systems Biology at Columbia University, New York, NY, USA; Department of Genetic and Genomic Sciences, Icahn School of Medicine at Mount Sinai, New York, NY, USA; Department of Obstetrics and Gynecology, Department of Genetics and Development, Columbia University Irving Medical Center, New York, NY, USA; Department of Medicine, Vagelos College of Physicians and Surgeons, Columbia University, New York, NY, USA; Cardiovascular Division, Washington University School of Medicine, St Louis, MO, USA; John Cochran Veterans Affairs Hospital, St Louis, MO, USA; Division of Cardiology and Vascular Medicine Institute, the Department of Medicine, and the Department of Veterans Affairs, University of Pittsburgh, Pittsburgh, PA, USA; Department of Pediatrics, University Medical Center Groningen, University of Groningen, Groningen, The Netherlands; Department of Gene Technology, Science for Life Laboratory, Royal Institute of Technology, Stockholm, Sweden

**Author notes:** **Corresponding authors:** Fang Li PhD, Cardiometabolic Genomics Program, Division of Cardiology, Department of Medicine, Columbia University Irving Medical Center, 630 West 168th Street, P&S10-401, New York, NY 10032, Hanrui Zhang MB PhD, Cardiometabolic Genomics Program, Division of Cardiology, Department of Medicine, Columbia University Irving Medical Center, 630 West 168th Street, P&S10-401, New York, NY 10032.

## Abstract

Translating human genomic discoveries into mechanistic insights requires linking genetic variations to candidate genes and their causal functional phenotypes. Genome-wide association studies have consistently identified *LIPA* as a risk locus for coronary artery disease (CAD), with previous expression quantitative trait loci (eQTL) analyses prioritizing *LIPA* as a candidate causal gene. However, functional studies elucidating the causal variants, regulatory mechanisms, target cell types, and their causal impact on atherosclerosis have been lacking. To address this gap, we applied functional genomics and experimental mouse models to establish the variant-to-function relationship at the *LIPA* locus. Our findings show that CAD risk alleles in the *LIPA* locus increase LIPA expression and enzyme activity specifically in monocytes/macrophages by enhancing PU.1 binding to an intronic enhancer region that interacts with the *LIPA* promoter. In myeloid *Lipa*-overexpressing mice, we observed larger atherosclerotic lesions accompanied by altered macrophage function, including increased macrophage accumulation due to enhanced monocyte recruitment, reduced neutral lipid accumulation, and upregulation of integrin and extracellular matrix pathways. Our work establishes a direct causal link between *LIPA* risk alleles and increased monocyte/macrophage LIPA that exacerbates atherosclerosis, bridging human functional genomic evidence to the mechanistic understanding of CAD.

**Graphical Abstract:** 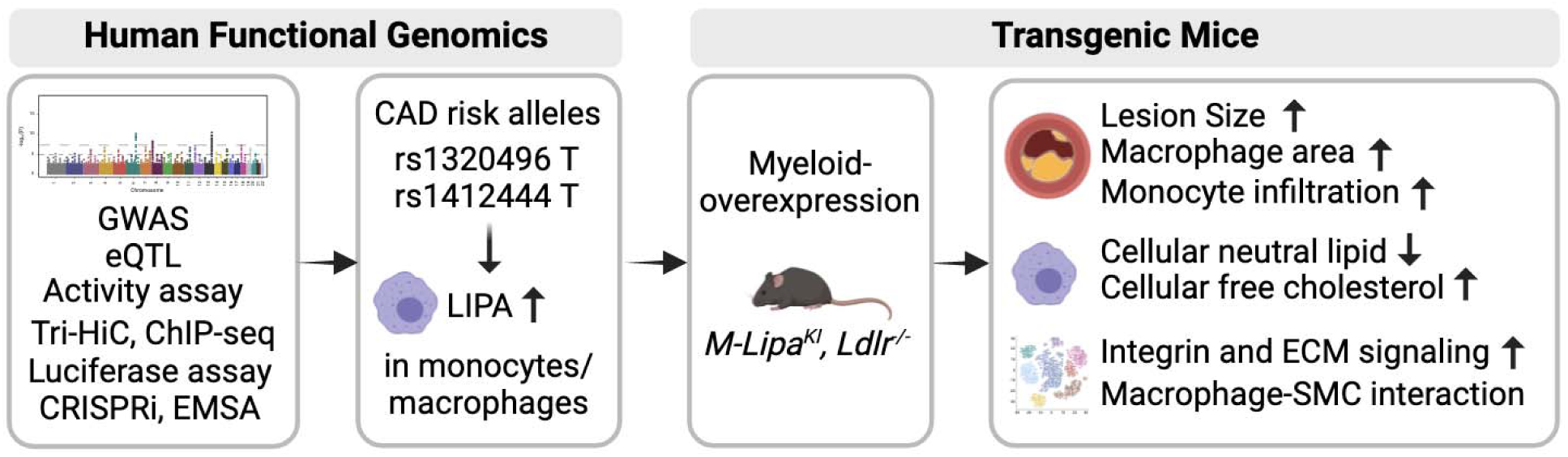

## Introduction

Coronary artery disease (CAD) remains the leading cause of death worldwide. The inheritability of CAD has been estimated to be 40% to 60%, supporting the contribution of inherited genetic variation to its pathogenesis (1). Genome-wide association studies (GWASs) determine the common genetic variants associated with diseases and traits, having collectively identified more than 300 CAD-associated risk loci (2–13). Identifying the causal genes that connect genetic variation to disease risk will inform novel biological mechanisms and therapeutic targets for CAD.

The *LIPA* locus has been reproducibly identified as a risk locus for CAD in multiple GWASs and meta-analyses across ethnic groups (**Table S1**) (2–13), but not plasma lipid traits (14), indicating a CAD-specific yet lipid metabolism-independent genetic mechanism. Colocalization analysis suggests that genetic variants associated with CAD are also associated with *LIPA* expression in the Stockholm-Tartu Atherosclerosis Reverse Networks Engineering Task study (STARNET) dataset (15), suggesting *LIPA* as the candidate causal gene at the locus. The *LIPA* gene encodes lysosomal acid lipase (also known as LAL), the only known acidic lipase hydrolyzing cholesteryl ester and triglyceride within lysosomes (16). Rare coding and splicing junction mutations in *LIPA* result in Mendelian disorders, including the infant-onset Wolman Disease due to a complete loss of LIPA and the later-onset Cholesteryl Ester Storage Disease (CESD) with up to 10% residual enzyme activity (16). CESD is characterized by hyperlipidemia, hepatosplenomegaly, and premature atherosclerosis, presumably due to severe dyslipidemia (17). Mice with whole body knockout of *Lipa* phenotypically resemble the late-onset CESD in humans despite the complete loss of *Lipa*, and die within days when on an *Ldlr^−/−^* background and challenged with dietary cholesterol (18). However, almost all common CAD variants at the *LIPA* locus, including the lead single nucleotide polymorphism (SNP) rs1412444, are intronic, suggesting that they likely exert *cis*-regulatory effects rather than impacting protein function directly. The sole exonic SNP in this linkage disequilibrium (LD) block, i.e. rs1051338, does not alter the expression, activity, lysosomal trafficking, or secretion of LIPA, further implicating that these noncoding variants likely contribute to CAD risk by modulating gene regulation via *cis*-regulatory effects (19).

The SNP rs1412444, and other SNPs in high LD, showed robust quantitative trait locus (QTL) signals in monocytes, with risk alleles of CAD linked to higher *LIPA* mRNA (2, 3), and more importantly, higher enzyme activity (19) in human peripheral blood monocytes. These findings support that CAD risk alleles are associated with higher LIPA activity in monocytes, which may be causally involved in CAD. Pathologically, monocytes play essential roles in the development and exacerbation of atherosclerosis. In atherosclerosis, plaque macrophages are largely derived from the infiltration of circulating monocytes (20). Monocytes and macrophages may therefore represent the most relevant cell types for the regulatory effects of CAD *LIPA* variants.

Despite strong statistical association and initial QTL studies supporting the directionality and the potential causal cell types, the variant-to-function relationship and how this important locus causally impacts atherosclerosis are unclear. Herein, we sought to delineate the causal variants, the effector gene, the cell types involved, and the regulatory mechanisms of this CAD risk locus; and to elucidate how elevated LIPA in these key causal cell types, as implicated by human functional genomic evidence, contributes to atherosclerosis pathogenesis. This work connects CAD-associated *LIPA* genetic variants to their biological functional, advancing our understanding of this locus’s causal role in elevating CAD risk.

## Results

### Integrative genomic analyses identify *LIPA* as the candidate causal gene within the *LIPA* locus

GWASs have identified a number of common genetic variants at the *LIPA* locus that are strongly associated with risks of CAD as summarized in **Table S1**. Leveraging the CARDIoGRAMplusC4D GWAS dataset (7, 13), we identified twenty SNPs (P < 1 × 10^−8^) significantly associated with CAD risk, with rs1412444 as the lead SNP (P = 5.15 × 10^−12^) **(Figure 1A** and **Table S2**).

**Figure 1.**
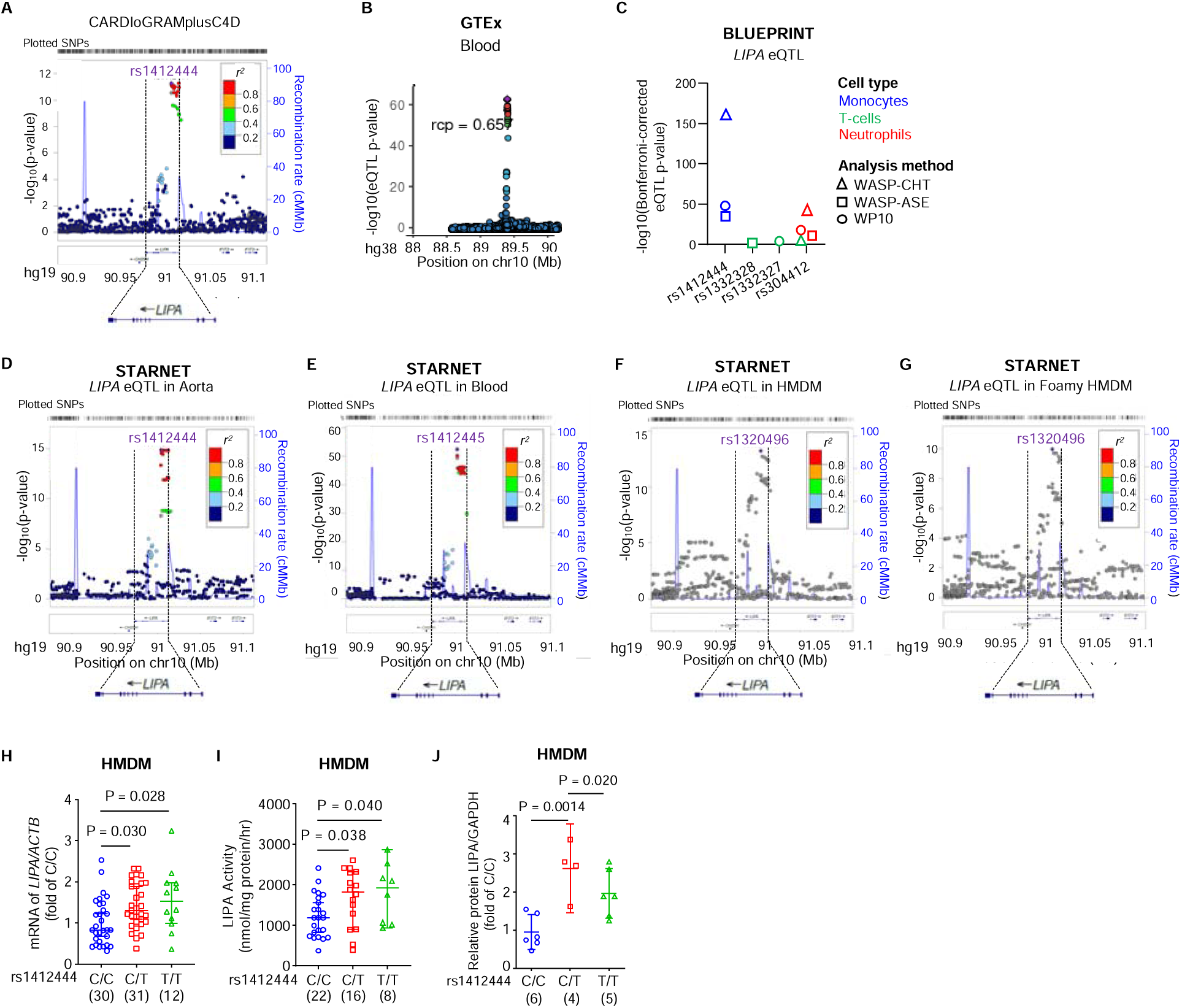
Genetic variants at the *LIPA* locus associated with higher risks of coronary artery disease (CAD) are also linked to higher *LIPA* mRNA, enzyme activity, and protein in human peripheral blood monocytes-derived macrophages (HMDM). **A,** LocusZoom plot to visualize the CARDIoGRAMplusC4D CAD GWAS signals at the *LIPA* locus. The GWAS lead SNP rs1412444 and the SNPs in linkage disequilibrium (LD) are color-coded based on pairwise *r^2^* with rs1412444. **B,** Colocalization analysis by integrating CARDIoGRAMplusC4D GWAS dataset with the Genotype-Tissue Expression project (GTEx) eQTL in the blood shows that *LIPA* is the only nearby gene at the locus (± 1 Mb flanking the *LIPA* gene) showing strong regional colocalization probability (rcp = 0.657). **C,** eQTL analysis of *LIPA* in the BLUEPRINT dataset shows strong eQTL signals in monocytes (blue), but not T-cells (green) or neutrophils (red). Three eQTL methods, including WASP-corrected combined haplotype test (WASP-CHT, triangle), WASP-corrected allele-specific expression (WASP-ASE, square), and BLUEPRINT Work Package 10 (WP10, circle), were employed in the analysis. **D-G,** LocusZoom plots to visualize *LIPA* eQTLs in atherosclerotic aortic wall (Aorta, **D**), blood (**E**), HMDM (**F**), and foamy HMDM (HMDM loaded with acetylated low-density lipoprotein, **G**) in the Stockholm-Tartu Atherosclerosis Reverse Network Engineering Task (STARNET) dataset, which comprises nine cardiometabolic tissues/cell types from subjects with CAD. The most significant SNP is marked by purple diamond. Other SNPs in the region are color-coded by their LD with the lead SNP according to pairwise *r^2^*. **H-J**, The risk allele (T) of the CAD lead SNP rs1412444 is associated with increased *LIPA* mRNA (**H**), enzyme activity (**I**), and protein (**J**) in HMDM. n = the number of subjects. Data are presented as mean ± SEM in (**H**) and median ± 95% Cl in (**I**) and (**J**).

To identify the causal gene and causal tissue/cell types at the locus, we applied integrative genomic analysis pipeline employing three methods from two broad classes: 1) Colocalization analysis; 2) Summarized Mendelian Randomization (SMR) followed by Heterogeneity in Dependent Instruments Test (HEIDI) (21) or MetaXcan (22). Using CARDIoGRAMplusC4D GWAS and GTEx eQTL data, which shows the strongest *LIPA* eQTL in blood (**Figure S1A**), colocalization analysis by ENLOC (23) suggest that *LIPA* is the only nearby gene showing strong regional colocalization probability (rcp) in this region (± 1 Mb flanking gene *LIPA,* rcp = 0.657, whereas all other genes have rcp = 0) (**Figure 1B** and **Figure S1B)**. By integrating CARDIoGRAMplusC4D and UKBB GWASs with STARNET eQTL data of 9 tissue/cell types in this region (± 500 kb flanking gene *LIPA*), we confirmed that *LIPA* showed significant colocalization signal (Coloc H_4_ > 0.8) in CAD-relevant tissue/cell types (**Table S3**). SMR-HEIDI test examines if the same variant is associated with gene expression and complex traits (24). SMR-HEIDI analysis also identified *LIPA* as the only candidate causal gene with P_SMR_ < 0.001 and P_HEIDI_ >= 0.05 (**Table S4)**. MetaXcan further confirmed *LIPA* as the candidate causal gene demonstrating the strongest MetaXcan evidence (P < 5 × 10^−8^) (**Table S5**). Expanding on prior work that hinted at *LIPA* as a causal gene (2, 3), our integrative genomic analyses provided strong statistical evidence establishing *LIPA* as the causal gene at this locus.

### Functional genomic data support monocytes and macrophages as the causal cell types

Functional regulatory variants and genomic elements often exert tissue- and cell type-specific effects to regulate the expression of their target genes. Settling causal tissues/cell types will enhance our understanding of disease-relevant regulatory mechanisms. We applied three strategies leveraging large-scale transcriptomic and epigenomic datasets to elucidate the involved cell types for the regulatory role of the *LIPA* CAD variants.

First, GTEx eQTL analysis revealed the strongest *LIPA* eQTL association in whole blood, followed by spleen and visceral adipose tissue (**Figure S1A**). Given the high immune cell content in these tissues, we examined the BLUEPRINT dataset to identify specific immune cell association (25). As shown in **Figure 1C**, monocytes showed the strongest eQTL associations for CAD SNPs across all three eQTL methods, including WASP-CHT (WASP-corrected combined haplotype test),(26) WASP-ASE (WASP-corrected allele-specific expression) (26), and WP10 eQTL, while T-cells showing no eQTL signals. Neutrophil exhibited a weak eQTL signal for SNP rs304412, which lacks significant CAD association (**Table S2**).

Second, eQTL studies in diseased tissue allow the discovery of associations not identifiable in healthy tissues, likely due to shifts in cell composition and regulatory landscape changes. Using STARNET, with eQTL studies in cardiometabolic tissues from patients with CAD, and CARDIoGRAMplusC4D datasets (± 500 kb flanking gene *LIPA*), our analysis found more evidence for the role of monocytes and macrophages in CAD-risk at the *LIPA* locus: 1) atherosclerotic aortic wall (AOR), blood, and human peripheral blood monocyte-derived macrophage (HMDM) (**Figure 1D-1F**, respectively) represent the strongest evidence as the causal tissues/cell types (P_eQTL_ < 5 × 10^−8^, **Table S6**), passing colocalization, SMR-HEIDI, and metaXan tests (Coloc H_4_ > 0.8, P_SMR_ < 0.001 and P_HEIDI_ >= 0.05, and P_metaXan_ < 5 × 10^−8^, **Table S3-S5**); 2) foamy macrophages (acetylated-LDL loaded HMDMs) showed strong eQTL (**Figure 1G**) meeting SMR-HEIDI (**Table S4**); 3) *LIPA* eQTL associations in liver, visceral abdominal adipose (VAF), and skeletal muscle (SKLM) had P_eQTL_ < 5 × 10^−8^ (**Figure S2, Table S6**), though SMR-HEIDI suggested that different variants (in high LD) independently regulated CAD risk and *LIPA* expression in these tissues (P_HEIDI_ < 0.05, **Table S4**); 4) no *LIPA* eQTL association was found in pre/early-atherosclerotic mammary arteries (MAM, **Figure S2**, **Table S6**), the healthy arteries mainly consist of endothelial cells and vascular smooth muscle cells unlikely to have immune cell accumulation. These results together support that monocytes/macrophages are likely the causal cell types for the *LIPA* locus.

Finally, we harnessed single-cell datasets to provide insights into cell type-specific regulatory mechanisms. Single-cell ATAC-seq of human atherosclerotic plaque identifies cell type-specific regulatory elements, with recent scATAC-seq of human atherosclerotic plaques (n = 41 subjects and 28,316 cells) revealing a macrophage-specific regulatory element containing rs1320496 and the lead SNP (rs1412444) for CAD (27). The regulatory element was absent in vascular smooth muscle cell-derived cells and other cell types in the plaque, underscoring a macrophage-specific regulatory mechanism. In addition, in the STARNET dataset, rs1320496 was the lead eQTL SNP (P = 1.11 × 10^−11^) for *LIPA* in foamy macrophages (**Figure 1G, Table S6**). Collectively, bulk and single-cell data highlight monocytes and macrophages as the causal cell type for *LIPA* variant regulation.

### *LIPA* CAD risk alleles are associated with higher *LIPA* mRNA, protein, and enzyme activity in monocytes and macrophages

Upon establishing the causal cell types, it is important to confirm how the CAD risk alleles affect *LIPA* expression in these cells. Previous eQTL studies show that the risk alleles (T) of rs1412444 and rs2246833 (in high LD, *r^2^* = 0.98) are associated with higher *LIPA* mRNA (2, 3), and more importantly, higher LIPA enzyme activity (19) in human monocytes. In light of the strong *LIPA* eQTL signals in HMDM and foamy macrophages (**Figure 1F** and **1G**) and the vital role of macrophages in atherosclerosis (28), the cause of CAD, we examined *LIPA* expression and enzyme activity in human macrophages.

Using HMDMs differentiated from peripheral blood mononuclear cells isolated from fully genotyped healthy individuals (**Table S7**), we observed a 1.42-fold and 1.57-fold increase in *LIPA* mRNA for C/T and T/T genotypes of rs1412444, respectively, and significant increase in LIPA enzyme activity (by 1.37-fold for C/T and 1.47-fold for T/T) compared to C/C, the non-risk genotype (**Figure 1H** and **1I**). LIPA protein levels also increase in risk allele carriers, confirmed by Western blot (29) (**Figure 1J**). Thus, both previous functional genomic datasets and our experimental cohort validate that CAD risk alleles link to higher LIPA expression and enzyme activity in monocytes (19) and macrophages.

### Experimental validation establishes the functional regulatory element and the regulatory mechanisms of the candidate causal variants

Identifying causal variants and linking them to the target gene remains challenging, yet it represents a critical step to definitively establish the causality of the locus. Most disease-associated noncoding variants reside in *cis*-regulatory elements (cREs) that regulate gene expression in a cell type-specific manner (30). We experimentally validated the cREs and functional variants regulating *LIPA* expression using the following strategies: 1) prioritize cREs harboring GWAS variants using ChIP-seq data in monocytes and macrophages (31) and luciferase assay; 2) use high-resolution Tri-HiC to identify chromatin looping between *LIPA* promoter and putative cREs, and CRISPR interference (CRISPRi) to validate the role of the cRE in regulating *LIPA* expression; 3) perform luciferase assay with site-directed mutagenesis to identify functional variant(s); and 4) analyze allele-specific binding (ASB) and motifs to understand transcription factor (TF) interactions.

CAD GWASs reproducibly identified rs1412444 and rs2246833, both intronic in *LIPA*, as the lead SNP or tag SNP (**Table S1**). Using epigenomic data in monocytes and macrophages (31), we visualized the locus, including histone marks (H3K4me1 and H3K27ac) and TF binding sites (**Figure 2A** and **Figure S3**). The rs1412444 region aligns with stronger enhancer marks and PU.1 binding (**Figure 2A**), showing enhancer activity by luciferase assay in THP-1 monocytes but not HEK293 cells, indicating a monocyte/macrophage-specific regulatory mechanism (**Figure 2B**). The rs2246833 region showed no enhancer activity in either cell line (**Figure 2B**), consistent with weaker enhancer features (**Figure 2A**).

**Figure 2.**
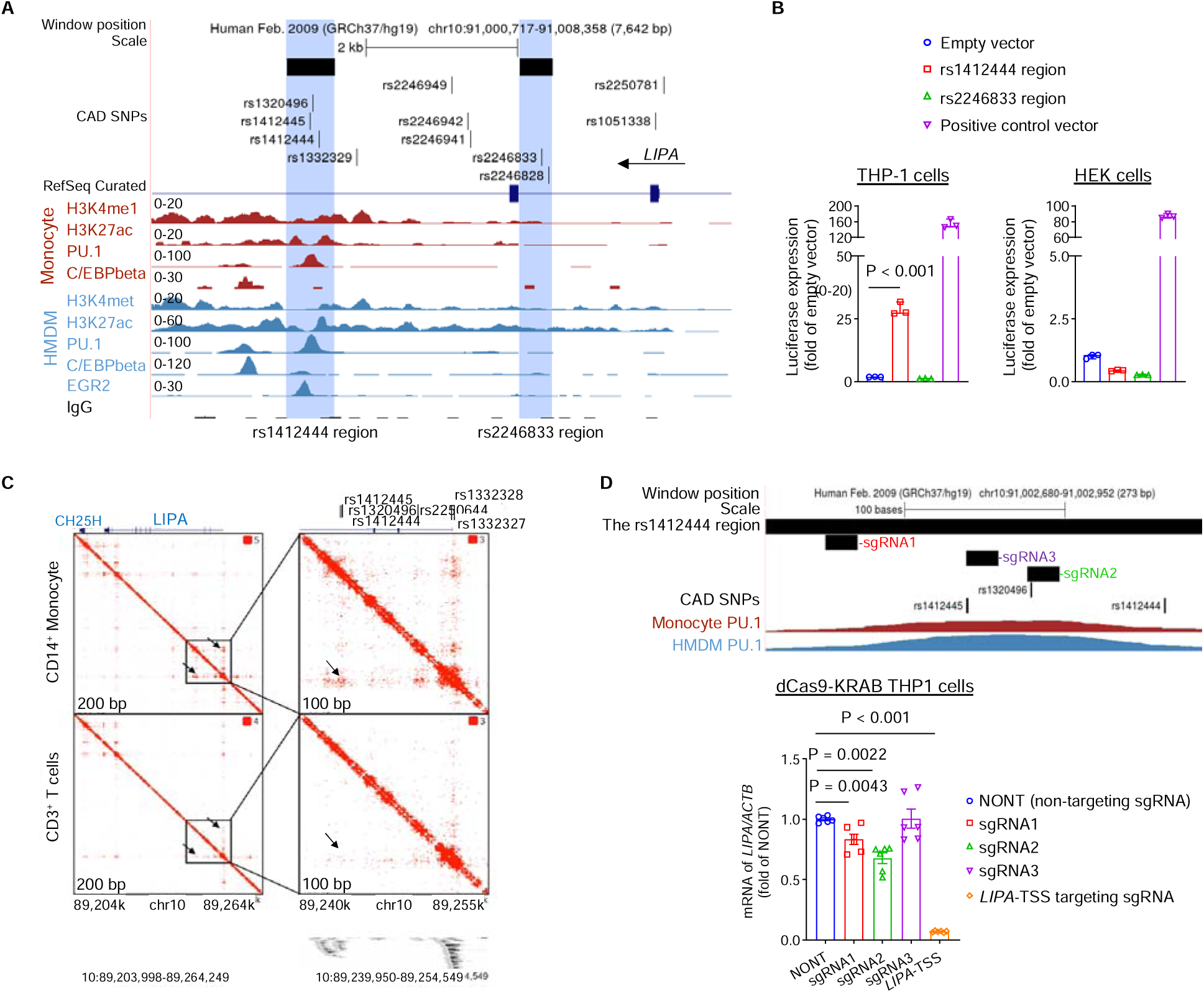
The non-coding genomic region containing rs1412444 is a myeloid-specific enhancer, interacting with the *LIPA* promoter and regulating *LIPA* expression. **A,** Genome Browser view of annotation tracks for the *LIPA* locus showing significant CAD GWAS SNPs, including the lead SNP (rs1412444) and SNPs in linkage disequilibrium (LD) as listed in **Table S2**, *LIPA* transcripts, and the regulatory landscape in human monocyte and HMDM. The potential enhancer regions showing H3K4me1 and H3K27ac modifications and hosting CAD GWAS SNPs, including the rs1412444 region and the rs2246883 region, are highlighted and prioritized for subsequent functional validation. **B,** The rs1412444 region, but not the rs2246833 region, shows enhancer activity by luciferase assay in THP-1 monocytes (**left**). Neither region shows enhancer activities in HEK 293 cells (**right**). (n = 3 independent experiments with 3 technical replicates, data are presented as median ± 95% CI.) **C,** High-resolution Tri-HiC depicts chromatin interaction between the rs1412444 region and the *LIPA* promoter in human CD14^+^ monocytes, but not CD3^+^ T-cells. **D,** CRISPRi targeting the rs1412444 enhancer region reduces *LIPA* expression. sgRNAs targeting the rs1412444 region were designed and transduced to dCas9-KRAB expressing THP-1 cells for CRISPRi-mediated silencing and mRNA expression of *LIPA* was determined by quantitative RT-PCR. The sgRNA1 is 68 bp upstream of rs1412445 and is the highest scored sgRNA within the region by CRISPick. The sgRNA2 span rs1320496. The sgRNA3 span rs1412445. The rs1412444 SNP region lacks a PAM sequence for sgRNA design. The sgRNA targeting the transcription start site of *LIPA* (*LIPA-* TSS) serves as the positive control. (n = 6 replicates, data are presented as mean ± SEM.)

Enhancers regulate gene expression by recruiting TFs and looping with the target gene promoter, a key mechanism for how noncoding variants influence transcription (30). Using high-resolution Tri-HiC data (32), we identified a strong interaction between the rs1412444 region and the *LIPA* promoter in human monocytes, but not T-cells (**Figure 2C**), aligning with eQTL data showing lack of eQTL association in T-cells (**Figure 1C**). CRISPRi targeting the rs1412444 region suppressed *LIPA* expression in THP-1 cells, especially by gRNA spanning rs1320496 (**Figure 2D**).

The rs1412444 region, containing SNPs rs1412445, rs1320496 and rs1412444, was prioritized as the causal cRE to further identify the functional variants in the region using both experimental and statistical methods. The SNP rs1412444 is the lead SNP in CAD GWAS (**Figure 1A**) and *LIPA* eQTL in atherosclerotic aortas (**Figure 1D**). The SNP rs1412445 is in extremely high LD with rs1412444 (*r^2^* = 0.99), thereby serving as a proxy. The SNP rs1320496 shows the strongest signal for *LIPA* eQTL in HMDM and foamy macrophages (**Figure 1F-1G**), and is partly linked with SNP rs1412444 (*r^2^* = 0.45, *D’* = 1). Among three major haplotypes (CCC, CTC, and TTT, defined by rs1412445, rs1320496, and rs1412444 genotypes, **Table S8**), compared to the non-risk haplotype CCC, the CTC and CTT haplotypes showed increased enhancer activity in monocytes by luciferase assay (**Figure 3A**), suggesting that both rs1412445/rs1412444 and rs10320496 may affect *LIPA* expression. Conditional analysis partially disentangles the contribution of rs1412445/rs1412444 and rs1320496 on *LIPA* expression and CAD risk, with both SNPs significantly linked to LIPA expression and CAD when analyzed independently, supporting the presence of multiple causal variants within the rs1412444 enhancer region (**Figure 3B**).

**Figure 3.**
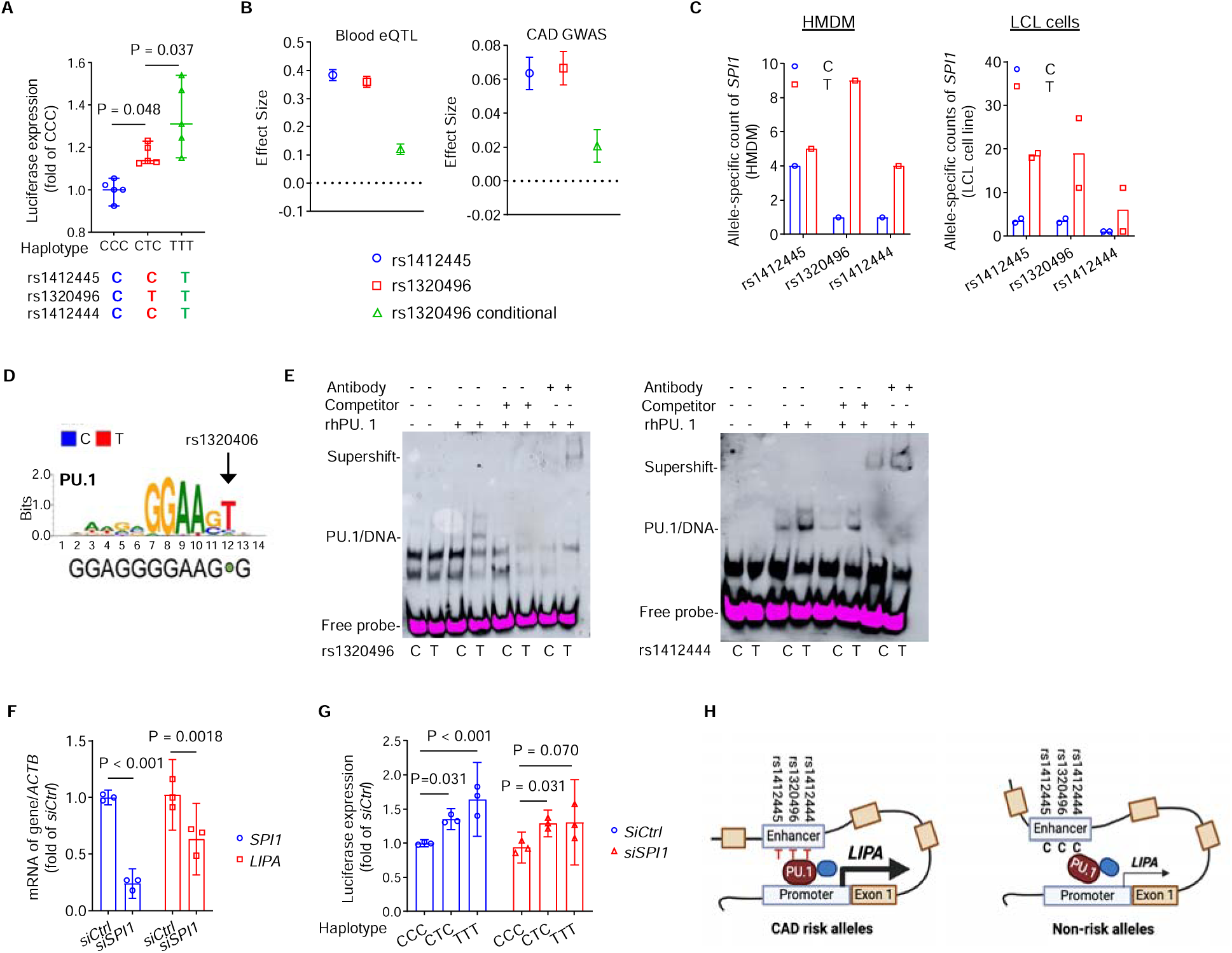
rs1320496 and rs1412444 represent the functional SNPs at the *LIPA* locus, with risk alleles enhancing PU.1 binding and regulating *LIPA* expression. **A,** Site-directed mutagenesis and luciferase assay confirmed that the risk haplotypes TTT (rs1412445-T, rs1320496-T, and rs1412444-T) and CTC (rs1412445-C, rs1320496-T, and rs1412444-C) lead to increased enhancer activities compared to the non-risk haplotype CCC (rs1412445-C, rs1320496-C, and rs1412444-C). (n = 5 independent experiments with 3 technical replicates, data are presented as median ± 95% CI.) **B,** Conditional analysis was performed to assess the independent contribution of SNPs (rs1412445, rs1320496, and rs141244) within the rs1412444-containing enhancer to eQTL and GWAS signals. The perfect LD between rs1412444 and rs1412445 (*r^2^* = 0.99) allows for the utilization of rs1412445 as a representative to capture the effects of both variants. The high LD (*r^2^* = 0.45) between rs1320496 and the other two variants prohibited full disentanglement. Both rs1412445 (blue) and rs1320496 (red) show independent association with *LIPA* expression (left, in blood, P < 10^−59^) and CAD risk (right, P < 10^−11^). After conditioning on rs1412445, rs1320496 remains showing nominally significant effects on both *LIPA* expression (left, in blood, P = 3 × 10^−10^) and CAD risk (right, P = 0.031) (green, rs1320496 conditional), suggesting that both SNPs independently contributing to the traits. In the context of eQTL analysis, the effect size is quantified as the log_2_ allelic fold change in *LIPA* gene expression, while in GWAS, it represents the log odds change in CAD risks. **C,** Allele-specific binding analysis of PU.1 ChIP-seq data in HMDM (GSM785501, n = 1 experiment) and LCL cell line (EBV-transformed lymphoblastoid B-cell lines) (ENCODE dataset, n = 2 independent experiments). Results suggest that PU.1 binds more favorably to the risk alleles (T). **D,** Motif analysis shows that the risk allele (T, red) of rs1320496 creates a PU.1 binding site. **E**, Electrophoretic Mobility Gel Shift Assay (EMSA) was conducted to determine the effects of different alleles on PU.1 binding for sequences containing SNP rs1320496 (left) and rs1412444 (right), using recombinant human PU.1 protein and anti-PU.1 antibody. Data shown are representative images from n = 2 independent experiments. **F**, siRNA-mediated knockdown of *SPI1* (encoding PU.1) in THP-1 cells reduces mRNA expression of *LIPA*. (n = 3 independent experiments with 2 technical replicates, data are presented as median ± 95% CI.) **G**, Knockdown of *SPI1* partly abolished the effects of risk alleles on increasing the enhancer activity of the rs1412444 region. Data were analyzed by two-way ANOVA followed by Tukey’s multiple comparisons test. (n = 3 independent experiments with 3 technical replicates, data are presented as median ± 95% CI.) **H**, Schematic figure summarizing results of functional genomic studies that the risk alleles of the functional variants at *LIPA* risk locus increases the expression of *LIPA* by enhancing PU.1 binding in monocytes/macrophages and via enhancer-promoter interaction.

We next sought to understand the mechanisms of action of the functional variants in the rs1412444 enhancer region. Many eQTL SNPs regulate target gene expression by altering TF binding. Using eQTL data from 49 tissues in GTEx, we have previously established that analyzing eQTL effect size as a function of TF expression level offers a generalizable approach to discover TF regulators of genetic variant effects (33). Using this analysis pipeline, we observed significant correlations between eQTL effects of rs1320496 and the expression levels of 9 TFs, with PU.1, encoded by the *SPI1* gene, showing the strongest effect (Spearman rho = 0.48, P = 6.0 × 10^−4^, **Figure S4A**). Indeed, the rs1412444 enhancer region overlaps with the PU.1 binding site, an important TF required for myeloid differentiation (**Figure 2A**) (34). We, therefore, hypothesize that the functional variants in the rs1412444 region may alter PU.1 binding affinity. To test this, we used: 1) allele-specific PU.1 ChIP-seq; 2) motif analysis; 3) electrophoresis mobility shift assay (EMSA); and 4) knockdown of *SPI1* to confirm causality. PU.1 ChIP-seq (35) showed higher PU.1 binding for the risk alleles (T) of rs1320496 and rs1412444 compared to non-risk alleles (C) (**Figure 3C**). Among the SNPs, only rs1320496 (T) was predicted to directly change a key nucleotide to enhance motif-matching (**Figure 3D**). EMSA further validated that the T allele of rs1320496 created a binding site for PU.1 (**Figure 3E**). The T allele of rs1412444 showed increased PU.1 binding compared to the C allele (**Figure 3E**), while the T allele of rs1412445 showed no evidence of binding to PU.1 (**Figure S4B**). We further validated that knockdown of *SPI1* reduces *LIPA* mRNA expression (**Figure 3F**), and partially abolished the effects of the risk alleles on increasing the enhancer activity of the rs1412444 region (**Figure 3G**). These findings confirm that risk alleles of rs1320496 and rs1412444 increase *LIPA* expression and contribute to CAD risk by enhancing PU.1 binding. (**Figure 3H**)

We also examined TFs beyond PU.1. A modest correlation between rs1412444 eQTL effects and *STAT1* expression was noted in GTEx data (Spearman rho = 0.26, P = 0.07, **Figure S5A**) and ENCODE data (**Figure S5D**), with STAT1 ChIP-seq in CD14^+^ monocytes showing an allelic imbalance (**Figure S5B**). However, *STAT1* knockdown did not affect *LIPA* expression (**Figure S5C**), suggesting STAT1 unlikely regulate *LIPA* CAD variants effects.

In summary, our functional genomic studies confirm that the risk alleles of genetic variants associated with CAD lead to increased *LIPA* mRNA, protein, and enzyme activity in monocytes and macrophages. These variants within the enhancer region interact with the *LIPA* promotor and regulate *LIPA* expression by altering PU.1 binding. Having established the causative role of these variants in elevating LIPA, we next aim to assess whether this increase exacerbates atherosclerosis using myeloid-specific *Lipa* overexpression in preclinical mouse models.

### Preclinical mouse studies demonstrate that myeloid-specific overexpression of *Lipa* increases atherosclerosis

To elucidate how increased myeloid LIPA impacts atherosclerosis *in vivo*, we developed a myeloid-specific *Lipa* overexpression mouse model (*LysMCre^+/−^, Lipa^KI/WT^, Ldlr^−/−^*, referred to as *M-Lipa^KI^*, **Figure 4A** and **S6A** for illustration of the construct and breeding strategy). Their corresponding littermates *LysMCre^−/−^, Lipa^KI/WT^, Ldlr^−/−^* were used as control (*Ctrl*). The percentage of GFP^+^ cells, a reporter of *Lipa* overexpression, was quantified across multiple tissues to demonstrate specificity and efficiency of overexpression (**Figure S6C-S6E**). In *M-Lipa^KI^* mice, GFP^+^ cells comprised 36.2%, 22.0%, 34.8%, and 68.5% of CD45^+^CD115^+^ monocytes/macrophages in blood, bone marrow (BM), spleen, and peritoneum, respectively (**Figure S6C**). GFP^+^ cells represented 46.3% and 70.2% of CD115^−^Ly6G^hi^ neutrophils in blood and BM (**Figure S6D**), with minimal GFP^+^ CD3^+^ lymphocytes (< 2.7%, **Figure S6E**), confirming myeloid-specific overexpression. The efficiency of *LysMCre* recombination is comparable to previous reports on *LysMCre*-mediated overexpression in mouse models (36).

**Figure 4.**
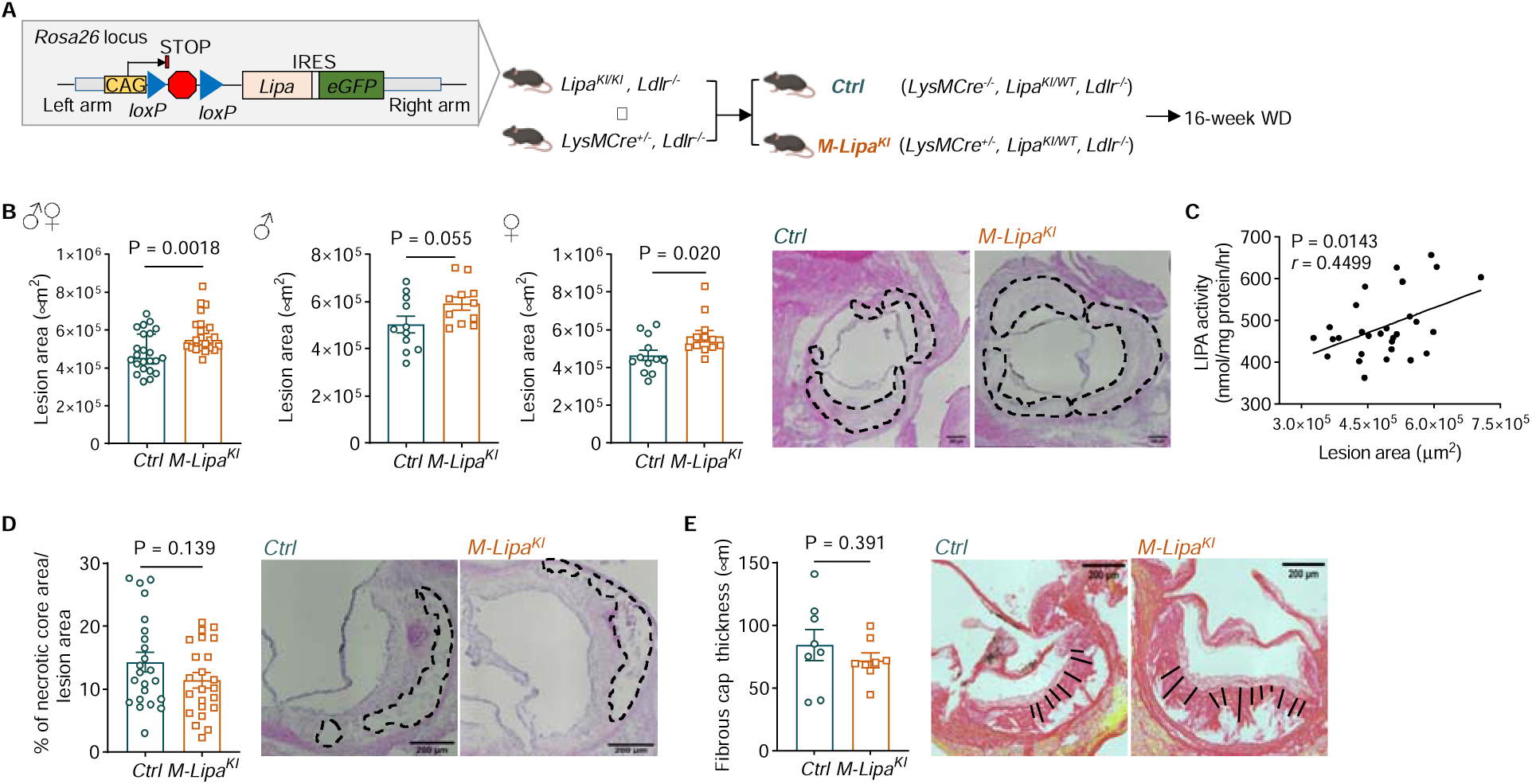
Myeloid overexpression of *Lipa* exacerbates atherosclerosis in *Ldlr^−/−^* mice. **A**, The plasmid construction and breeding strategy of the mouse model. Mice for conditional overexpression of *Lipa* (NM_001111100) were generated by *Rosa26* knock-in (KI) of CAG-loxP-STOP-loxP-Lipa-IRES-eGFP. To achieve myeloid-specific overexpression, *Lipa^KI/KI^* mice were bred with *LysMCre^+/−^* mice (heterozygous for the Cre allele) to delete the *loxP-STOP-loxP* cassette, thereby enabling overexpression of *Lipa* in myeloid cells in *LysMCre^+/−^*, *Lipa^KI/WT^* mice. Littermates with the genotype *LysMCre^−/−^, Lipa^KI/WT^* without overexpression serve as controls. To assess the impact of myeloid-specific overexpression of *Lipa* on atherosclerosis, these mice were bred onto an *Ldlr^−/−^* background. To induce atherosclerosis, *Ctrl* (*LysMCre^−/−^, Lipa^KI/WT^, Ldlr^−/−^*) and *M*-*Lipa^KI^* (*LysMCre^+/−^, Lipa^KI/WT^, Ldlr^−/−^*) mice were fed a Western diet (WD) for 16 weeks. The atherosclerotic lesion size and features of plaque stability in the aortic sinus were quantified. **B,** Myeloid overexpression of *Lipa* modestly but significantly increased atherosclerotic lesion size in mice of both sexes (left: combined, n = 23 mice), as well as in male mice (middle: n = 11) and female mice (right: n= 12) when analyzed separately. Data are presented as median ± 95% CI. Scale bar = 200 μm. **C**, Atherosclerotic lesion size is positively correlated with LIPA enzyme activity in peritoneal macrophages (PMs) (two-tailed Pearson’s correlation analysis, P = 0.014, *r* = 0.4499). **D**, No differences in the percentage of necrotic core area between the two genotypes (n = 11 male mice and 12 female mice). **E**, Sirius red staining shows comparable fibrous cap thickness between the two genotypes (n = 8 male mice). Data are presented as mean ± SEM. Scale bar = 200 μm for (**D**) and (**E**).

The overexpression of *Lipa* at mRNA, protein, and enzyme activity levels was confirmed in bone marrow-derived macrophages (BMDMs) and peritoneal macrophages (PMs) from *M-Lipa^KI^* mice of both sexes (**Figure S6F-S6O**). In male mice, *Lipa* mRNA and enzyme activity increased by 1.56-fold and 1.26-fold in *M-Lipa^KI^* BMDMs (**Figure S6F and S6G**) and by 3.49-fold and 1.44-fold in *M-Lipa^KI^* PMs (**Figure S6K and S6L**) compared to *Ctrl*, respectively. In female mice, the *Lipa* mRNA and enzyme activity increased by 1.41-fold and 1.21-fold in *M-Lipa^KI^* BMDMs (**Figure S6H and S6I**) and 2.52-fold and 1.24-fold in *M-Lipa^KI^* PMs (**Figure S6M and S6N**) compared to *Ctrl*, respectively. Consistently, LIPA protein significantly increased in both BMDMs (**Figure S6J**) and PMs (**Figure S6O**) from *M-Lipa^KI^* mice compared to *Ctrl*. Despite the modest overexpression, the enzyme activity increase was comparable to HMDMs of risk allele carriers (**Figure 1I**), supporting *M-Lipa^KI^* mice as a model for studying the causal effects of gain-of-function of LIPA observed in human functional genomic data.

*Ctrl* and *M-Lipa^KI^* mice were fed a Western diet (WD, TD88137, Envigo Teklad) for 16 weeks to induce atherosclerosis, with body weight, organ weight, and lipid profiles assessed (**Figure S7A-S7N**). Body weight, spleen weight/body weight, and liver weight/body weight were comparable between *Ctrl* and *M-Lipa^KI^* mice in both males (**Figure S7A-S7C**) and females (**Figure S7H-S7J**). No differences in total cholesterol (**Figure S7D** and **S7K**) or cholesterol in low-density lipoprotein (LDL) and high-density lipoprotein (HDL) fractions were observed (**Figure S7E** and **S7L,** after 4 h fasting and in both sexes). Female *M-Lipa^KI^* mice showed a trend toward higher plasma triglyceride (P = 0.050, **Figure S7M**) and higher triglyceride in both LDL and HDL fractions (**Figure S7N**), an effect not seen in males (**Figure S7F** and **S7G**). Overall, myeloid *Lipa* overexpression did not alter body weight or plasma cholesterol, consistent with human data linking *LIPA* variants to CAD but not metabolic traits, while the modest effects of myeloid *Lipa* overexpression on triglyceride metabolism specifically in female mice on a WD require further examination (37).

Atherosclerotic lesion size was significantly larger, with a 17.78% increase in male and 21.85% in female *M-Lipa^KI^* mice compared to *Ctrl* mice (**Figure 4B**). Lesion size positively correlated with LIPA activity in PMs (**Figure 4C**, P = 0.014, *r* = 0.4499), supporting the association between increased macrophage LIPA activity and atherogenesis. No differences in necrotic core area or fibrous cap thickness, features of plaque instability (38), were observed (**Figure 4D**, **4E**).

In summary, our findings highlight a pro-atherogenic role for increased myeloid Lipa expression without affecting body weight or lipid profile, in line with human genomic data.

### Myeloid *Lipa* overexpression promotes atherosclerosis with an increased macrophage content in the lesion, mainly attributable to increase monocyte recruitment

Myeloid *Lipa* overexpression significantly increased the CD68^+^ macrophage area in lesions for both sexes (**Figure 5A**), with no differences in macrophage proliferation or apoptosis, as determined by Ki67 and TUNEL (terminal deoxynucleotidyl transferase dUTP nick-end labeling) staining (**Figure 5B-5C**), suggesting other mechanisms for the increase.

**Figure 5.**
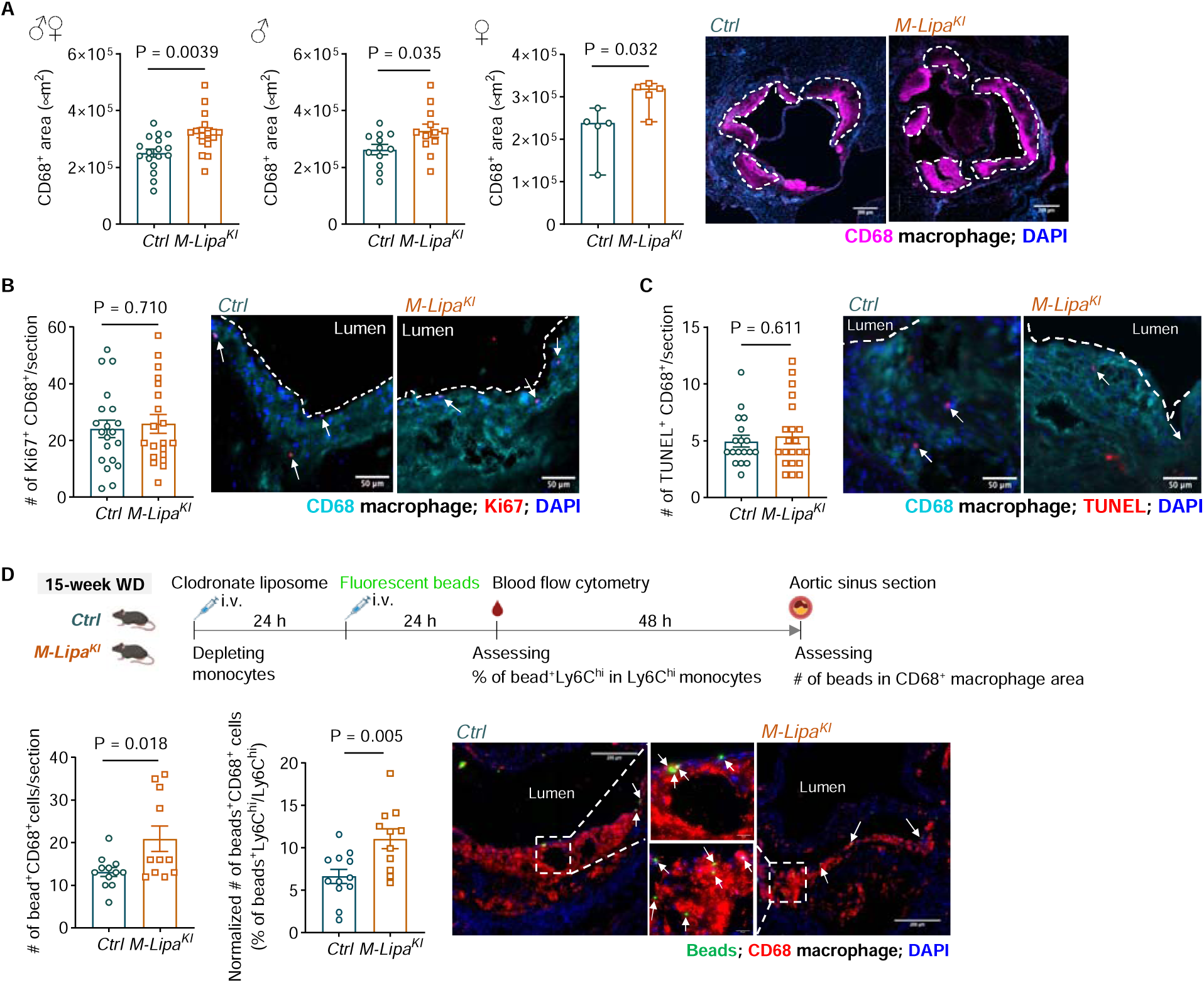
Myeloid overexpression of *Lipa* increases macrophage content in the atherosclerotic plaque by promoting monocyte recruitment. **A**, Immunofluorescence (IF) staining data suggest that myeloid overexpression of *Lipa* significantly increased the CD68^+^ [Magenta] macrophage area in the lesion in mice of both sexes (left: combined, n = 17 mice), as well as in male mice (middle: n = 12) and female mice (right: n = 5). Data are presented as median ± 95% CI. The white dashed contour indicates lesion area. Scale bar = 200 μm. **B-C**, IF staining and quantification for the proliferative macrophages (**B**, Ki67 [Red], CD68^+^ [Cyan], and DAPI [Blue]) and apoptotic macrophages (**C**, TUNEL [Red], CD68^+^ [Cyan], and DAPI [Blue]). (**B**, n = 13 male mice and 7 female mice; **C**, n = 13 male mice and 8 female mice). Data are presented as mean ± SEM. Scale bars = 50 μm. **D**, Monocyte infiltration was assessed by beads assay. Briefly, Ly6C^hi^ monocytes in the blood were pulse-labeled with fluorescent beads by transiently depleting circulating monocytes with clodronate liposome injection. Beads labeling efficiency was assessed by flow cytometry as the % of bead^+^Ly6C^hi^ monocytes in Ly6C^hi^ monocytes 24 h after beads injection. The number of beads in the CD68^+^ area of atherosclerotic lesion was quantified in mice three days post-injection. Results are presented as the number of beads in the CD68^+^ area per section (left), the number of beads in the CD68^+^ area per section normalized by the percentage of bead^+^ monocytes in Ly6C^hi^ monocytes (middle), and representative IF staining showing beads^+^ [Green] CD68^+^ macrophages [Red] in the lesion (right). For *Ctrl*, n = 3 male mice and 8 female mice; for *M*-*Lipa^KI^*, n = 3 male mice and 8 female mice; the average of two sections per mouse were reported. Data are presented as mean ± SEM. Scale bars = 200 μm.

During atherogenesis, blood monocytes continuously influx into the subendothelial space and contribute to lesion macrophage accumulation (28). Since plaque macrophage proliferation or apoptosis remained unaffected, we hypothesize that increased macrophage area may result from increased monocyte recruitment. To test this, Ly6C^hi^ circulating monocytes were pulse-labeled with fluorescent beads (**Figure 5D**) as previously described (39). Three days after labeling, bead numbers within the lesion were quantified. *M-Lipa^KI^* mice showed increased beads in CD68^+^ macrophage area within plaques (**Figure 5D**), indicating higher monocyte recruitment. We further confirmed that *M-Lipa^KI^* with WD feeding did not alter myelopoiesis in blood or BM (**Figure S8A and S8B**) or affect the percentage of hematopoietic progenitor cells in BM (**Figure S8C**). Chemokines, *e.g.,* CCL2 (40, 41), CCL8 (42), and CCL7 (43), have been implicated in regulating monocyte infiltration in atherosclerosis progression. The levels of plasma CCL2, CCL7, and CCL8, and 13 other chemokines and cytokines, showed no difference between genotypes (**Figure S9**). These results suggest that myeloid *Lipa* overexpression increases monocyte recruitment to the plaques, likely due to changes in the local plaque microenvironment rather than monocyte numbers or systemic inflammation.

### Myeloid *Lipa* overexpression leads to reduced neutral lipid accumulation and increased free cholesterol accumulation in macrophages

Beyond the increased macrophages in *M-Lipa^KI^* plaques, we further validated the functional effects of *Lipa* overexpression in macrophages. Using flow cytometry, we characterized foamy (SSC^hi^LipidTOX^hi^) aortic macrophages (CD45^+^CD11b^+^CD64^+^) (**Figure 6A** for study design and **Figure S10A** for gating strategy) (44). We confirmed that in atherosclerotic plaques, the percentage of SSC^hi^LipidTOX^hi^ foamy aortic macrophages was similar between *M-Lipa^KI^* and *Ctrl* mice (**Figure 6B**), but the mean fluorescent intensity (MFI) of LipidTOX in foamy aortic macrophages isolated from *M-Lipa^KI^* mice was significantly lower than that in *Ctrl* mice (**Figure 6C**), indicating reduced neutral lipids. Within *M-Lipa^KI^* mice, GFP^+^ aortic macrophages had lower neutral lipids than GFP^−^ aortic macrophages (**Figure 6D**), confirming intrinsic effects driven by *Lipa* overexpression.

**Figure 6.**
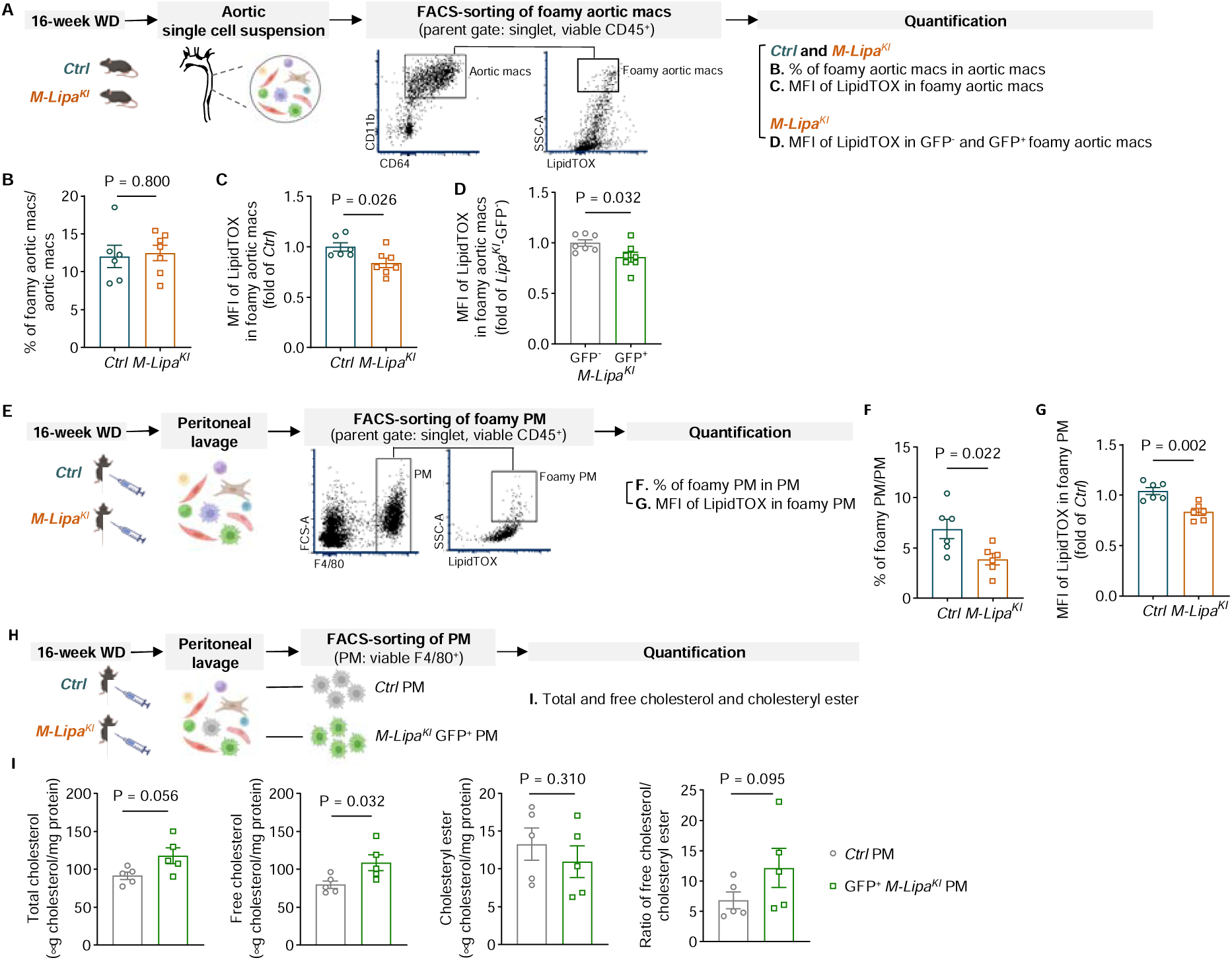
Myeloid overexpression of *Lipa* leads to reduced neutral lipid but increased free cholesterol accumulation in macrophages. **A,** Schematics of a flow cytometry-based method to characterize aortic macrophages (CD45^+^CD11b^+^CD64^+^) that are neutral lipid-enriched and foamy (SSC^hi^LipidTOX^hi^) in aortas dissected from *Ctrl* and *M*-*Lipa^KI^* mice fed a WD for 16 weeks. **B,** Percentage of foamy aortic macrophages in total aortic macrophages in *Ctrl* and *M*-*Lipa^KI^* mice. **C**, Mean fluorescent intensity (MFI) of LipidTOX in foamy aortic macrophages in *Ctrl* and *M*-*Lipa^KI^* mice. **D,** MFI of LipidTOX in GFP^−^ and GFP^+^ foamy aortic macrophages in *M*-*Lipa^KI^* mice. **B-D**, n = 6 male mice for *Ctrl*; n = 7 male mice for *M*-*Lipa^KI^*. Data are presented as mean ± SEM. **E,** Schematics of study design to characterize foamy (SSC^hi^LipidTOX^hi^) PMs (CD45^+^F4/80^+^) in *Ctrl* and *M*-*Lipa^KI^* mice fed a WD for 16 weeks. **F,** Percentage of foamy PMs in PMs of *Ctrl* and *M*-*Lipa^KI^* mice. **G,** MFI of LipidTOX in foamy PMs in *Ctrl* and *M*-*Lipa^KI^* mice. **F-G**, n = 6 male mice. Data are presented as mean ± SEM. **H,** F4/80^+^ PMs from *Ctrl* mice and F4/80^+^GFP^+^ PMs from *M*-*Lipa^KI^* mice that fed a WD for 16 weeks were sorted for quantification of cellular cholesterol levels in (**I**). **I,** Quantification of total cholesterol (left), free cholesterol (second left), cholesteryl ester (second right), and ratio of free cholesterol/cholesteryl ester (right). (n = 5 female mice, with analysis based on the averages of two technical replicates. Data are presented as median ± 95% CI.)

We also assessed SSC^hi^LipidTOX^hi^ foamy PMs (CD45^+^F4/80^+^) from WD-fed mice (**Figure 6E** for study design and **Figure S10B** for gating strategy). The percentage and LipidTOX MFI of foamy PMs were significantly lower in *M-Lipa^KI^* mice compared to *Ctrl* mice, indicating reduced neutral lipid accumulation (**Figure 6F**, **6G**). Fluorometric cholesterol assays in PMs (**Figure 6H** for study design) further showed higher total and free cholesterol but lower cholesteryl ester levels in *M-Lipa^KI^* PMs (**Figure 6I**), aligning with LipidTOX findings. These data consistently show reduced neutral lipid accumulation in *Lipa^KI^* aortic macrophages and PMs.

We further explored if reduced neutral lipid accumulation in *M-Lipa^KI^* PMs from WD-fed mice could be replicated *in vitro* in cultured macrophages loaded with modified lipoproteins. PMs from mice fed a normal laboratory diet (ND) were treated with oxidized-LDL (oxLDL, **Figure S11A** for study design). We indeed observe a lower MFI of LipidTOX in oxLDL-loaded *M-Lipa^KI^* PMs compared to *Ctrl* PMs (**Figure S11B**), which was not due to a significant change in oxLDL-binding or uptake (**Figure S11C** and **S11D**). As expected, *Lipa* mRNA was elevated in *M-Lipa^KI^* PMs (**Figure S11E**), while genes for lipoprotein uptake were similar (**Figure S11E)**. *M-Lipa^KI^* PMs had lower expression of cholesterol biosynthesis genes (**Figure S11F**), suggesting increased free cholesterol accumulation in the endoplasmic reticulum thus a suppression of cholesterol biosynthesis genes. No differences were seen in cholesterol efflux gene expression (45), including *Abca1* and *Abcg1* (**Figure S11G**). It is worth noting that oxLDL loading only led to a slight increase in total and free cholesterol, but levels were similar between *M-Lipa^KI^* and *Ctrl* PMs (**Figure S11H**). Thus, oxLDL-loaded PMs replicate some, but not all, phenotypic features of PMs from WD-fed mice, underscoring the importance of carefully assessing the appropriate cellular model for mechanistic studies.

### Myeloid *Lipa* overexpression distinctly alters the transcriptomic signature of aortic macrophages and PMs

Building on the observed phenotypic changes, we sought to determine how myeloid *Lipa* overexpression alters macrophage transcriptomes. To compare the transcriptomic signature of aortic macrophages between *Ctrl* and *M-Lipa^KI^* mice fed a WD for 16 weeks, we performed bulk RNA-seq using the low-input RNA-seq method owing to the limited materials of aortic macrophages (**Figure 7A** for study design). We identified 300 upregulated and 123 downregulated DE genes in *M-Lipa^KI^* vs. *Ctrl* aortic macrophages (**Figure 7B** and **Table S9**, absolute fold change > 1.5, FDR-adjusted P < 0.05). Gene Set Enrichment Analysis (GSEA) showed enrichment of integrin signaling and cell-matrix adhesion pathways in the upregulated DE genes (**Figure 7C** and **Table S10**). The increased integrins (*Itga6*, *Itga9*, *Itgb3*, *Itgax*, etc.) and adhesion molecules and receptors (*Cspg4*, *Dag1, Fn1, Kdr, Tnc, Npnt,* etc.) in *M-Lipa^KI^* aortic macrophages may contribute to the increased monocyte adhesion and infiltration and macrophage adhesion in the atherosclerotic plaque microenvironment, thus promoting atherosclerosis (**Figure 7D)**.

**Figure 7.**
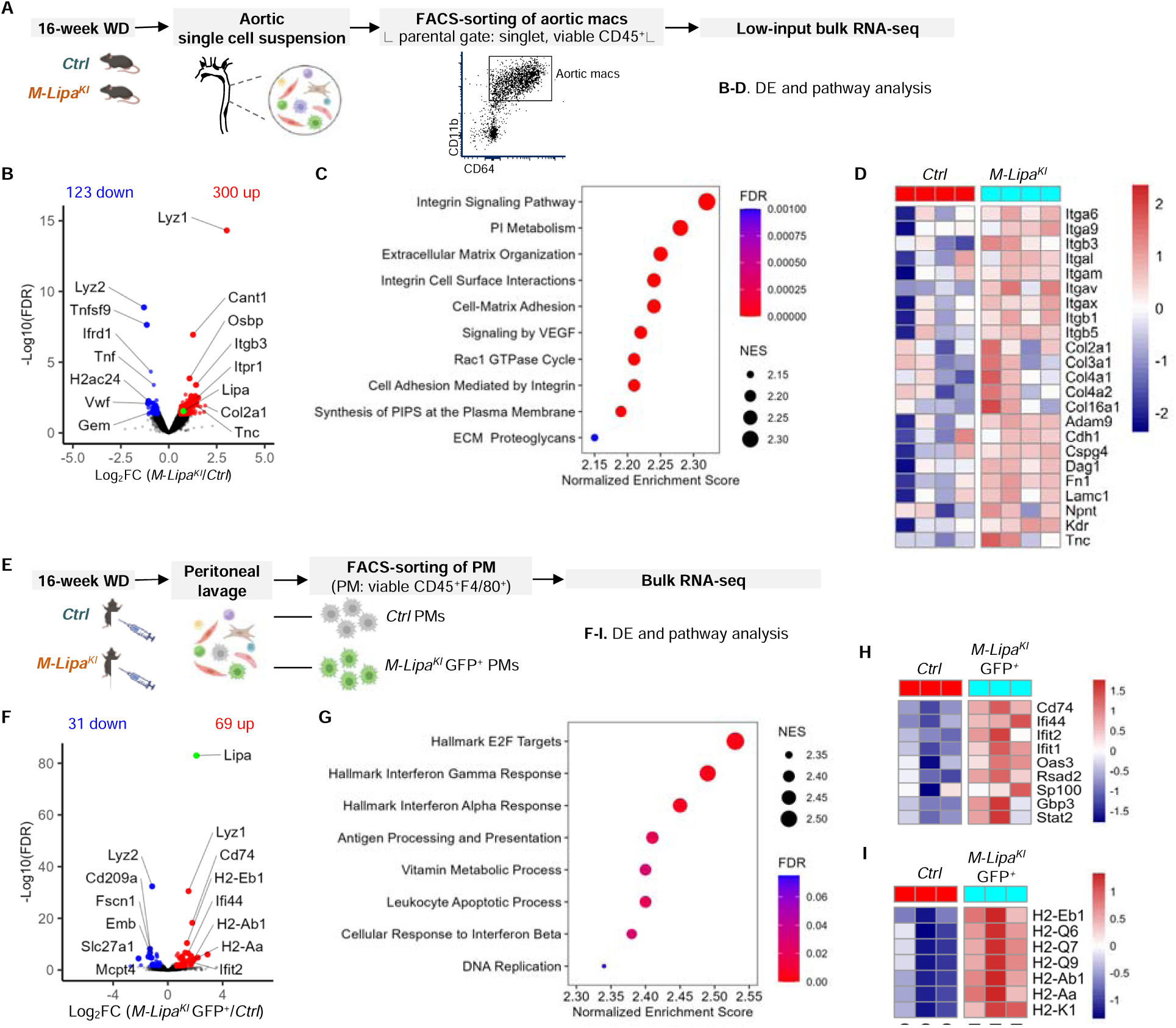
Bulk RNA-seq analyses reveal distinct effects of myeloid overexpression of *Lipa* on aortic macrophages and PMs. **A,** Viable macrophages (CD45^+^CD11b^+^CD64^+^) were sorted from aortas of *Ctrl* and *M*-*Lipa^KI^* mice fed a WD for 16 weeks and subjected to low-input RNA-seq. n = 4 biological replicates per genotype, with each replicate comprising pooled samples from 2-3 male mice, resulting a total of 10 male mice per genotype. **B**, Volcano plot to visualize the top DE genes (aortic macrophages isolated from *M*-*Lipa^KI^* mice *versus* those from *Ctrl* mice, all fed a WD for 16 weeks). **C**, The top enriched pathways in the upregulated genes in aortic macrophages of *M*-*Lipa^KI^* mice as determined by Gene Set Enrichment Analysis (GSEA). **D,** Heatmap visualization of top DE genes and the leading-edge subsets of the associated integrin signaling and cell-matrix mediated adhesion pathways. **E**, F4/80^+^ PMs from *Ctrl* mice and F4/80^+^GFP^+^ PMs from *M*-*Lipa^KI^* mice fed a WD for 16 weeks were sorted for bulk RNA-seq. n = 3 female mice. **F,** Volcano plot to visualize the top DE genes (F4/80^+^GFP^+^ PMs from *M-Lipa^KI^* mice *vs.* F4/80^+^ PMs from *Ctrl* mice, all fed a WD for 16 weeks). **G,** The top enriched pathways in the upregulated genes in *Lipa*-overexpressing PMs as determined by GSEA. **H-I,** Heatmap visualization of top DE genes and the leading-edge subsets of the associated interferon response pathways (**H**) and antigen processing and presentation pathways (**I**). **F-I**, n = 3 female mice.

To further examine if *Lipa* overexpression uniquely affect aortic macrophages, we performed RNA-seq on PMs from *Ctrl* and *M-Lipa^KI^* mice fed a WD for 16 weeks (**Figure 7E** for study design). We identified a total of 69 upregulated and 31 downregulated DE genes (**Figure 7F** and **Table S11**, absolute fold change > 1.5, FDR-adjusted P < 0.05). GSEA indicated enrichment in interferon signaling and antigen presentation pathways in the upregulated DE genes in *M-Lipa^KI^* vs. *Ctrl* PMs (**Figure 7G** and **Table S12**). This is consistent with the recent finding that lipid loading suppressed the expression of interferon-stimulated genes, e.g., *Ifit2*, *Ifit1*, *Oas3*, and *Stat2*, in mouse and human macrophages (46), as visualized in **Figure 7H**. The upregulated expression of major histocompatibility complex genes, e.g., *H2-Eb1*, *H2-Q6*, *H2-Ab1*, *H2-K1*, etc. in PMs of *M-Lipa^KI^* mice implicate immune activation (**Figure 7I**). To explore whether our observation of increased free cholesterol in PMs of *M-Lipa^KI^* mice may link to a proinflammatory transcriptomic signature, we further validated the mRNA of *Il18* and *Il1b* in PMs using qRT-PCR (**Figure S12**) and the plasma level of IL-18 and IL1β (**Figure S9**) in mice. However, no significant differences were observed between *Ctrl* and *M-Lipa^KI^*, suggesting NLRP3 inflammasome may not be the primary proinflammatory driving force in this context, despite the known role of excessive free cholesterol accumulation in the endoplasmic reticulum in activating the NLRP3 inflammasome (47, 48).

These findings emphasize the distinct effects of *Lipa* overexpression on aortic macrophages compared to other macrophage types, highlighting the importance of examining context-specific cellular responses in disease mechanisms.

### Single-cell RNA-seq analysis reveals that myeloid *Lip* a overexpression alters cell heterogeneity and transcriptomic profile of atherosclerotic aortic cells

The transcriptomic signature of increased integrin and extracellular matrix (ECM) genes in *Lipa*-overexpressing aortic macrophage prompts us to speculate whether the changes reflect altered macrophage subpopulations and whether macrophage transcriptomic changes can alter other plaque cell types via cell-cell interaction. To gain insights into the effects of myeloid *Lipa* overexpression on the phenotypic and functional heterogeneity of plaque cells, we carried out single-cell RNA-sequencing (scRNA-seq) analysis. Aortic single-cell suspensions were obtained from *Ctrl* and *M-Lipa^KI^* mice fed a WD for 16 weeks. Live single cells were sequenced using the 10X Chromium platform (**Figure 8A** for study design and **Figure S13A** for gating strategy). Data analysis was performed using Seurat v4.3.0.1 (49). Annotation of cell types was based on top marker genes of each cluster and by comparing with marker genes defined in previous meta-analyses of atherosclerotic plaque scRNA-seq data (**Table S13** for the top 15 marker genes and **Figure 8B** for the visualization of cell clusters and **Figure 8C** the top 3 marker genes) (50). Data confirmed an increased macrophage proportion in *M-Lipa^KI^* mice (**Figure 8D** and **Table S14**), consistent with histological data (**Figure 5A**). Results also suggest decreased SMC proportion and increased fibroblast proportion (**Figure 8D** and **Table S14**). Understanding the noisy nature of scRNA-seq, we performed DE analysis for each cell type to gain exploratory insights into effects of myeloid *Lipa* overexpression on the transcriptomic signature of different plaque cell types. While we observed modest changes, notably, among the 6 cell types, we identified the most DE genes in SMCs between *M-Lipa^KI^* and *Ctrl* mice (**Table S15** for DE genes and **Table S16** for canonical pathway analysis using Ingenuity Pathway Analysis, IPA). Focused analysis of SMCs reveals 6 SMC subclusters consistent with these identified in the literature (50) (**Figure 8E** for UMAP visualization and **Figure S13B** for marker gene visualization). In *M-Lipa^KI^* mice, a reduction in SMC1 and an increase in Cxcl12 SMC and fibroblast-like SMCs were observed (**Figure 8F**). IPA suggests the activation of canonical pathways in SMCs of *M-Lipa^KI^* mice, including the Integrin Cell Surface Interactions and Atherosclerosis Signaling canonical pathways (**Figure 8G** and **Table S16**), which corroborates with the observed increase in atherosclerotic lesion and the increased integrin and ECM genes in aortic macrophages by bulk RNA-seq. Upregulated DE genes in SMCs of *M-Lipa^KI^* mice include *Col1a1*, *Col1a2*, *Lum*, *Thbs1*, *Vcam1*, and *Mmp2* (**Table S15**), all of which have been implicated in contributing to increased ECM remodeling,(51) with *Vcam1* also playing a role in the recruitment and retention of immune cells within plaques (52). Among the 6 SMC subclusters, SMC1 and Cxcl12 SMCs showed the most DE genes between *Ctrl* and *M-Lipa^KI^* mice. IPA reveals activation of the Integrin Cell Surface Interactions canonical pathway in the SMC1 subcluster (**Figure 8H** and **Table S16**), while inhibition of the Smooth Muscle Contraction pathway in the Cxcl12 SMC subcluster (**Figure 8I** and **Table S16**). Collectively, these data suggest that in *M-Lipa^KI^* mice, plaque SMCs may undergo increased dedifferentiation, and macrophage-SMC interactions could be part of the mechanisms contributing to the enhanced atheroprogression. Even though the transcriptomic signature of macrophages did not differ between *M-Lipa^KI^* and *Ctrl* mice in this dataset (**Table S15** and **S16**), clustering analysis identified subclusters of aortic macrophages, including Trem2^+^, Ccl2^+^, Lyve1^+^ (resident-like), Il1b^+^ (inflammatory), and proliferative macrophages, aligning with those consistently reported in the literature (53, 54), showing comparable subcluster proportions (**Figure S13C-S13E**).

**Figure 8.**
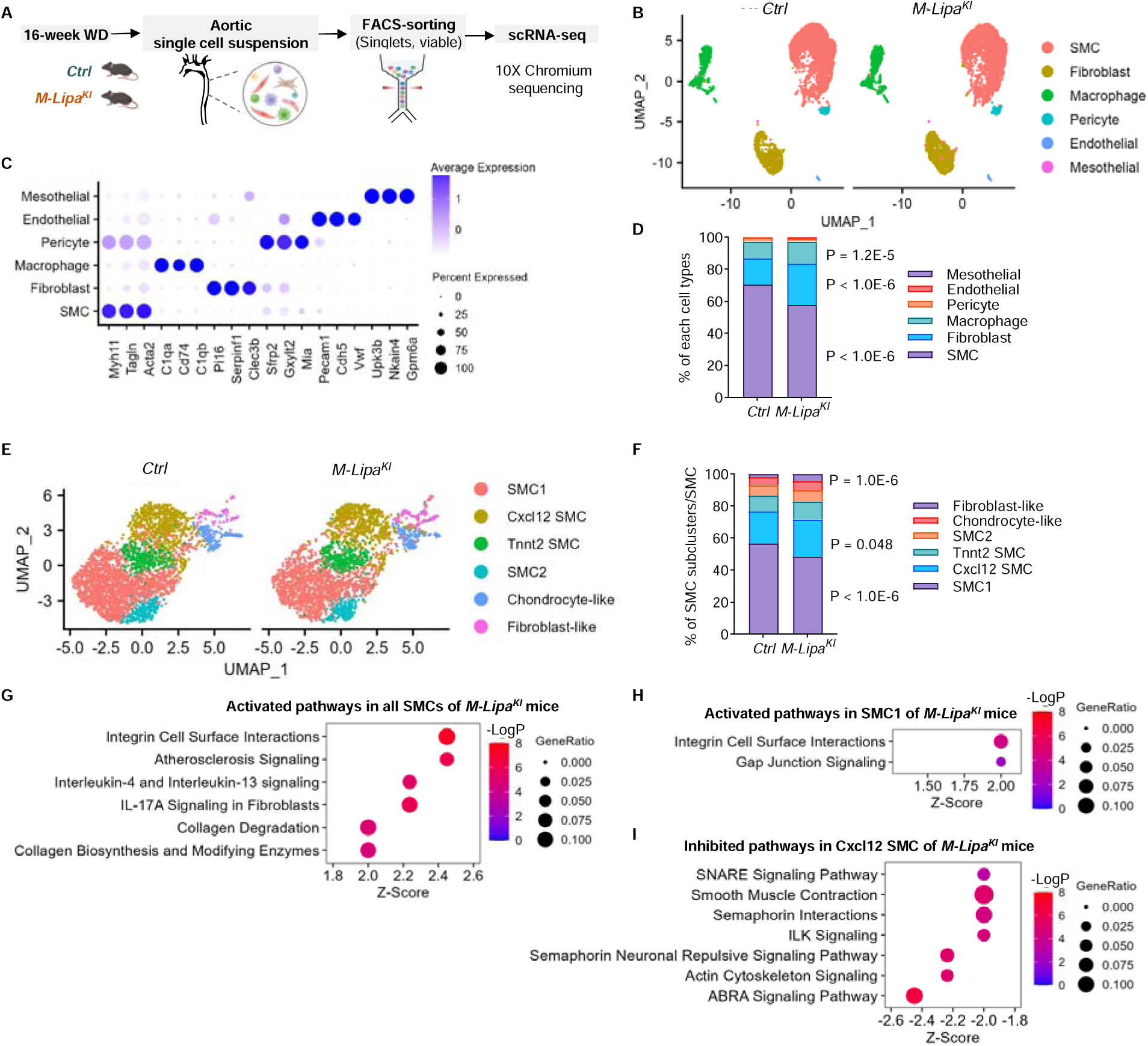
Single-cell RNA-seq (scRNA-seq) analyses suggest that myeloid overexpression of *Lip* a alters both the proportions and transcriptomic profiles of different aortic cell types in atherosclerosis, including smooth muscle cells. **A,** Viable cells isolated from aortas of *Ctrl* and *M*-*Lipa^KI^* mice fed a WD for 16 weeks were subjected to scRNA-seq. Cells were obtained from a pooled sample of n = 5 male mice. **B**, Uniform Manifold Approximation and Projection (UMAP) visualization of 6 cell types identified from the analyses. **C**, Dot plot visualization of top marker genes of each cell type. **D**, Stacked bar plot shows the proportion of each cell type in *Ctrl* (left) and *M*-*Lipa^KI^*(right) aortic cells. *M*-*Lipa^KI^* mice show increased macrophages and fibroblasts, while decreased smooth muscle cells (SMCs). Data were analyzed by Chi-square test with Bonferroni correction. **E**, Sub-clustering analysis of SMCs and UMAP visualization of 6 SMC subclusters. **F**, Stacked bar plot shows the proportion of each SMC subcluster in *Ctrl* (left) and *M*-*Lipa^KI^* (right) SMCs. Data were analyzed by Chi-square test with Bonferroni correction. **G**, Top activated canonical pathways by Ingenuity Pathway Analysis (IPA) in SMCs. **H**, Top activated canonical pathways by IPA in SMC1. **I**, Top inhibited canonical pathways by IPA in Cxcl12 SMC.

Given the transcriptomic differences in aortic macrophages observed in bulk RNA-seq data, we speculate that while scRNA-seq provides valuable information on cellular heterogeneity, it may be less sensitive in detecting transcriptomic differences, especially when sequencing all aortic cells with limited reads per cell type. To gain deeper molecular insights into the effects of *M-Lipa^KI^* on the phenotypic and functional heterogeneity of aortic macrophages, we conducted scRNA-seq analysis specifically focused on these cells. Since *LysMCre* does not result in 100% recombination, we leverage this by separating CD45^+^GFP^−^ (no *Lipa* overexpression) and CD45^+^GFP^+^ (*Lipa*-overexpressing) cells from enzymatically dissociated aortas of *M-Lipa^KI^* mice (**Figure S14A** for study design and **Figure S14B** for gating strategies). This approach allowed us to compare aortic macrophages with or without *Lipa* overexpression within the same plaque microenvironment (**Table S17** for the top 20 marker genes and **Figure S14C** for subcluster identification and assignment by comparing with marker genes defined in previous meta-analyses of atherosclerotic plaque scRNA-seq data) (53, 54).

A total of 12 distinct cell clusters were identified, including 5 macrophage clusters and 4 monocyte/dendritic clusters, as well as small numbers of B cells, fibroblasts, and SMCs that are present likely because of sorting-related contamination (**Figure S14C).** The GFP^+^ sample has more than 95% of the cells annotated as macrophages/monocytes, confirming the myeloid specificity of *Lipa* overexpression in our model. We have also validated the increased *Lipa* mRNA in GFP^+^ aortic macrophages compared to GFP^−^ aortic macrophages sorted from the aortas of *M-Lipa^KI^* mice, as well as in macrophages sorted from the aortas of *Ctrl* mice (**Figure S14D**). A total of 5 macrophage subclusters were identified in both GFP^−^ and GFP^+^ populations (**Figure S14E**), including Trem2 macrophages (cluster 0), Trem2 foamy macrophages (cluster 1), Lyve1 macrophages (cluster 2), Il1b macrophages (cluster 4), and proliferative macrophages (cluster 5). Both cluster 0 and cluster 1 had higher *Trem2* expression (**Figure S14E**), while cluster 1 had high expression of lipid processing genes *Fabp5* and *Gpnmb*, and lysosomal genes *Ctsl* and *Atp6v0d2* (**Figure S14E** and **Table S17**), a signature resembling increased lipid metabolism, that we refer to as Trem2 foamy macrophages.(54) Notably, the percentage of Trem2 foamy macrophages (cluster 1) is lower in the GFP^+^ population (17.32%) compared to the GFP^−^ population (36.00%, **Figure S14F** and **Table S18**). This lower proportion of lipid metabolism-associated aortic macrophage based on scRNA-seq analysis is consistent with lower neutral lipids in GFP^+^ aortic macrophages observed by flow cytometry (**Figure 6D**). DE and IPA analyses suggest cluster 0 Trem2 macrophages had the most DE genes (**Table S19**), representing a transcriptomic signature showing predicted activation of pathways involving oxidative stress and inflammation, such as iNOS Signaling, TREM1 Signaling, and TNFR1 Signaling (**Figure S14G** and **Table S20**); these pathways were also identified by IPA in the merged datasets between GFP- and GFP+ macrophages (**Figure S14H, Table S20**).

Taken together, histological and phenotypic analyses confirm an increased macrophage area and decreased neutral lipid accumulation in aortic macrophages of *M-Lipa^KI^* mice; scRNA-seq analysis further indicates that myeloid *Lipa* overexpression increases the overall proportion of plaque macrophages while reducing the proportion of the macrophage subcluster associated with lipid metabolism. Furthermore, the data suggest that myeloid *Lipa* overexpression influences SMCs, leading to a transcriptomic signature showing the activation of Integrin Cell Surface Interactions and Atherosclerosis Signaling canonical pathways, which may contribute to plaque progression.

## Discussion

Despite the first CAD GWAS being more than a decade ago, successful translation of genetic associations into causal genes and pathways has been confined to a small subset of GWAS loci (30). A number of genetic risk loci for CAD act through established CAD risk factors, such as lipid traits (13), which themselves have a significant genetic determination. Yet, many confirmed loci appear to act through novel mechanisms that are independent of traditional risk factors. Elucidating these mechanisms is critical for a better understanding of CAD pathogenesis and the identification of therapeutic targets. *LIPA* is one of the CAD loci that is specifically associated with CAD (2–13), but not with lipid traits (14). It is therefore of great interest to tackle the variant-to-function relationship for the *LIPA* locus and to determine how altered *LIPA* expression causally impacts atherosclerosis through mechanisms beyond its effects on plasma lipids. Here, using functional genomic approaches and experimental mouse models, we present evidence that *LIPA* is causal in CAD. Our results support that: 1) *LIPA* expression and enzyme activity are higher in HMDM of CAD risk allele carriers; 2) the CAD GWAS lead SNP rs1412444 resides in an enhancer region that shows strong interaction with the *LIPA* promoter in a monocyte-specific manner, with CRISPRi targeting the region suppressing *LIPA* expression; 3) within the enhancer region, risk alleles of rs1412444 and rs1320496 show higher binding to PU.1 and lead to increased enhancer activity; 4) Myeloid overexpression of *Lipa* leads to increased atherosclerosis lesion area and macrophage area in mice on *Ldlr^−/−^* background fed a WD, due to increased monocyte recruitment; 5) Myeloid overexpression of *Lipa* leads to an increase in free cholesterol and decrease in cholesteryl ester accumulation in PMs of mice fed a WD, accompanied by upregulation of genes in interferon signaling and antigen presentation pathways; 6) Myeloid overexpression of *Lipa* alters plaque cell heterogeneity and transcriptomic profile; the upregulation of genes in integrin signaling and cell-matrix adhesion pathways may contribute to macrophage-SMC interaction and facilitate monocyte recruitment, thus exacerbating atherosclerosis. Taken together, these data demonstrate that increased *LIPA* in myeloid cells drives atheroprogression, a directionality consistent with human GWAS and eQTLs finding for the *LIPA* locus (2–4).

Delineating causal variants is often challenging, partly because most GWAS variants are noncoding and in high LD with multiple variants within the same locus. A coding variant rs1051338, in high LD with the lead SNP rs1412444 (*r^2^* = 0.86), causes a nonsynonymous threonine to proline change within the signal peptide of LIPA protein. Because of the essential role of LIPA in lipid metabolism, the initial speculation was that rs1051338 serves as a causal variant contributing to lower LIPA activity (55). Further work by Evans *et al*. revealed that the nonsynonymous change within the signal peptide of LIPA protein did not affect protein trafficking, enzyme activity, or secretion (19). Evans *et al*. also confirmed that the risk allele (C) of rs1051338 is associated with increased *LIPA* expression and enzyme activity in human monocytes (19). Our work established the same directionality in human macrophages. We have also demonstrated the cell type-specific regulatory role of the *LIPA* CAD variants, *i.e.,* the enhancer activity of the rs1412444 region is monocyte-specific, and the chromatin interaction between the rs1412444 enhancer region and *LIPA* promoter was observed in monocytes, but not T-cells. We established that rs1320496 and rs1412444 are the functional SNPs within the rs1412444 enhancer region that regulate allele-specific interaction with PU.1, collectively exhibiting additive effects and suggesting a model of multiple causal variants within a single locus. Protein QTL (pQTL) studies represent a critical level of information for understanding the genetic regulation of protein expression. Although pQTL studies were performed in STARNET plasma, LIPA was not included in the original panel (NPX Olink platform). Future functional genomic data at the proteomic and metabolomic levels, in a cell and tissue-specific manner, and experimental validation by high-throughput functional genomic screening, will further enhance our ability to connect genetic variation to the functional mechanisms at scale.

This work has provided valuable mechanistic insights, while highlighting the importance of selecting appropriate cellular models for mechanistic studies. PMs of *M-Lipa^KI^* mice fed a WD showed increased free cholesterol accumulation and a transcriptomic signature showing upregulated interferon signaling. We attempted to model this phenotype *in vitro* using oxLDL-loaded PMs from mice fed a ND. However, our results did not show an effect of *M-Lipa^KI^* on oxLDL uptake or free cholesterol accumulation. OxLDL loading induced a modest increase in total cholesterol, in contrast to the remarkable total cholesterol and cholesteryl ester accumulation in PMs isolated from WD-fed mice. These results highlight that *in vitro* cultured macrophages loaded with chemically modified oxLDL do not fully resemble the foamy macrophages formed in an *in vivo* environment with WD-induced hyperlipidemia. Previous work suggests that lysosomal CE hydrolysis leads to free cholesterol accumulation in the lysosome when macrophages are loaded with oxLDL *in vitro*, disrupting lysosomal homeostasis, including the maintenance of an acidic pH (56). It has also been observed that macrophages form extracellular lysosomal synapses where the LIPA enzyme remains active and mediates catabolism of aggregated-LDL, generating free cholesterol for internalization (57). These *in vitro* studies align with our observation that *Lipa^KI^* increases free cholesterol accumulation. It is also worth noting that plaque macrophages demonstrate a transcriptomic signature of upregulated integrin and cell-matrix adhesion pathways, affirming increased monocyte infiltration and macrophage content in the atherosclerotic plaques of *M-Lipa^KI^* mice, while PMs of these mice showed enhanced interferon signaling. The differences in the effects of *Lipa* overexpression on the transcriptomic signature of PMs and plaque macrophages highlight the importance of local microenvironment in shaping cellular phenotypes.

While our work establishes *LIPA*’s genetic contribution to CAD, its physiological role as the only known acidic lipase in the lysosome, in intracellular lipid metabolism is equally critical. Loss-of-function of *LIPA* is the cause of Mendelian disorders characterized by lysosomal neutral lipid accumulation in multiple tissues (17). Enzyme replacement therapy with recombinant LIPA protein has demonstrated efficacy and safety in patients with *LIPA* deficiency (58). These findings have demonstrated the essential role of *LIPA* in lipid homeostasis and the benefits of supplementing LIPA when there is a severe LIPA deficiency. However, the effects of further increasing LIPA above the normal physiological level and for a prolonged period remain elusive. Loss of function of *LIPA* in humans led to severe dyslipidemia and hepatomegaly, prompting a hypothesis that overexpressing *LIPA* in the liver would prevent hepatic lipid accumulation. Yet, a recent study demonstrated that hepatic *LIPA* overexpression by AAV8 did not attenuate steatosis but unexpectedly exacerbated inflammation and immune cell infiltration in the liver of mice fed a WD (59). Our work using myeloid-specific *Lipa* overexpression mice established that increased *Lipa* in monocytes/macrophages increases atherosclerosis. Our study on the role of myeloid-specific *Lipa* overexpression in CAD prompts the needs to assess the benefits and risks in potentially therapeutically targeting LIPA in CAD. For example, the assumption that further enhancing LIPA will have a favorable role in accelerating lysosomal lipid catabolism should be carefully evaluated in a cell type and disease-specific manner.

In addition to the intracellular and cell-intrinsic effects observed with *Lipa* overexpression, another unexplored aspect is the potential transferability of the LIPA enzyme. This raises the question of how increased *Lipa* expression in myeloid cells might impact other cell types within the plaque, as well as in organs with resident or infiltrated macrophage populations. For instance, we observed altered SMC heterogeneity and transcriptomic profile, though it remains unclear whether this could be partly regulated by enzyme transfer that increases SMC LIPA levels. Additionally, we noted a slight elevation in triglyceride levels among female *M-Lipa^KI^* mice, while male mice did not show this change. Exploring the potential impact of myeloid-*Lipa* overexpression on liver and adipose biology stands as a future area of research. Moreover, our mouse model for conditional overexpression of *Lipa* presents an opportunity to explore the functions of *Lipa* and lysosomal lipid metabolism across various tissue and cell types, both under normal homeostasis and disease contexts.

In conclusion, our results established that *LIPA* risk alleles drive increased myeloid LIPA expression, which in turn aggravates atherosclerosis. These findings are consistent with and support human functional genomic discoveries, highlighting the critical role of LIPA in the progression of CAD. The workflow and functional genomic resources applied in this study provide valuable tools that will accelerate the systematic interrogation of genetic contributions to CAD risk, particularly through the roles of monocytes and macrophages.

## Methods

Detailed description of methods and materials are available in the online-only Supplemental Material.

### Sex as a biological variable

Our study examined male and female animals, and similar findings are reported for both sexes.

### Statistics

All statistical analyses were performed using GraphPad Prism 9 as described below, unless otherwise specified in the figure legends. When sample size (n) ≥ 6, data were tested for normality using Shapiro-Wilk test (when n < 8) or D’Agostino-Pearson test (when n ≥ 8). F-test of equality of variances was performed to compare the two sample variances. Data that passed normality tests are presented as mean ± standard error of mean (SEM) and analyzed using two-tailed Student’s *t*-test for comparison of two groups and equal variances (or with Welch’s correction if F-test was not satisfied), or one-way ANOVA with Tukey’s post-hoc tests for one independent variable with more than two groups. Data that did not follow a normal distribution were analyzed using nonparametric tests, *i.e.,* Mann-Whitney U test or Kruskal-Wallis test, and are presented as median ± 95% confidence interval (CI). Nonparametric tests were also used when n < 6. P < 0.05 was considered statistically significant. The number of independent experiments and biological replicates is specified in the figure legends.

### Study approval

All human study protocols were approved by the Human Subjects Research Institutional Review Board at Columbia University (AAAQ6510). All animal experiments were approved by the Institutional Animal Care and Use Committee at Columbia University (AABN5560). Mice were cared for according to the NIH guidelines.

### Data availability

Integrative genomic analyses have incorporated two data sources: GWAS summary statistics from the UK BioBank (UKBB) (60) or Coronary Artery Disease Genome Wide Replication and Meta-analysis plus The Coronary Artery Disease Genetics (CARDIoGRAMplusC4D) (7) and tissue/cell-type specific eQTLs from the Genotype-Tissue Expression project (GTEx) v8,(61) STARNET (15), or the BLUEPRINT project (25). GTEx is a large multi-tissue eQTL dataset containing 48 tissue types from ∼900 healthy subjects (61). STARNET is an eQTL dataset in nine disease-relevant metabolic tissues/cells collected from ∼600 CAD patients (15).

The bulk RNA-seq and single-cell RNA-seq datasets generated during the current study are deposited in the Gene Expression Omnibus (GEO) with accession number GSE243139 (secure token for the reviewers: kfwtsccihbufpgd). All code for data analysis associated with the current submission will be available via the lab GitHub repository at https://github.com/hanruizhang/ upon acceptance of the manuscript.

## Author contributions

FL and HZ conceived and designed the research. FL designed and conducted majority of the experiments. FL analyzed RNA-seq and scRNA-seq data. EF performed functional genomic analysis using GTEx and ENCODE data. HC performed functional genomic analysis using STARNET data. CX conducted imputation analysis and provided critical feedback on scRNA-seq data analysis. YZ performed and analyzed Tri-HiC data. PH, MNZ, JS, XW, ZW, YM, JC, AR, JC, AHO performed experiments. AJ, BR, MW, RCB, and YS provided input on experimental design. KH and TL were instrumental in the interpretation of functional genomic data. FL and HZ analyzed and interpreted results and wrote the paper. HZ directed and supervised the project and funding.

## Acknowledgements

We would like to acknowledge the NIH funding sources to the Columbia Center for Translational Immunology (CCTi) Flow Cytometry Core by grant number S10OD020056 and S10RR027050 and P30DK063608; the NIH-supported microscopy resources in the Center for Biologic Imaging, specifically the confocal microscope supported by grant number S10OD019973-01; the NIH/NCI Cancer Center Support Grant P30CA013696 for the use of resources at the Columbia Genome Center; and the Columbia Stem Cell Initiative (CSCI) Flow Cytometry Core under the leadership of Michael Kissner. We thank Dr. Chyuan-Sheng (Victor) Lin, Manager of the Genetically Modified Mouse Models shared resource at Columbia University Herbert Irving Comprehensive Cancer Center (HICCC), for their expertise in generating the mouse model for *Lipa* overexpression used in this study. The authors thank Drs. Towfique Raj and Jack Humphrey (Icahn School of Medicine at Mount Sinai), and Drs. Johan Björkegren (Karolinska Institutet) and Sean A. Bankier (University of Bergen) for access to pQTL studies performed in STARNET plasma samples. The authors would like to acknowledge Dr. Anna Ataran (Washington University in St. Louis) for support in sample collection. Schematic figures were created with BioRender.com.

