## Supplemental Material for "Decoding the variant-to-function relationship for *LIPA*, a risk locus for coronary artery disease"

Li F et al.

### DESCRIPTION OF SUPPLEMENTAL TABLES IN EXCEL FORMAT

**Table S1.** A list of previous GWASs that identified *LIPA* as a risk locus for CAD

**Table S2.** A list of SNPs associated with CAD at the *LIPA* locus

**Table S3.** Colocalization analysis by integrating STARNET datasets with CARDIoGRAMplusC4D and UKBB GWASs

**Table S4.** SMR-HEIDI analysis by integrating STARNET datasets with CARDIoGRAMplusC4D and UKBB GWASs

**Table S5**. MetaXcan analysis by integrating STARNET datasets with CARDIoGRAMplusC4D and UKBB GWASs

**Table S6.** *LIPA* eQTL in STARNET tissues

**Table S7.** Genotype imputation in our human cohort using Haplotype Reference Consortium reference panel

**Table S8.** The three haplotypes identified in the rs1412444 region

**Table S9**. DESeq2 output of bulk RNA-seq in aortic plaque macrophages (*M-Lipa*$^{KI}$ *vs. Ctrl*)

**Table S10**. Gene Set Enrichment Analysis output of bulk RNA-seq in aortic plaque macrophages (*M-Lipa*$^{KI}$ *vs. Ctrl*)

**Table S11.** DESeq2 output of bulk RNA-seq in peritoneal macrophages (PMs) (*M-Lipa*$^{KI}$ *vs. Ctrl*)

**Table S12.** Gene Set Enrichment Analysis output of bulk RNA-seq in peritoneal macrophages (PMs) (*M-Lipa*$^{KI}$ *vs. Ctrl*)

**Table S13.** Top 15 marker genes for subclusters of macrophages and SMCs identified in scRNA-seq of aortic cells

**Table S14**. Proportion/cell number of different cell types and subclusters of macrophages and SMCs in scRNA-seq of aortic cells

**Table S15**. Differential expression analysis of different cell types and their subclusters identified in scRNA-seq of aortic cells

**Table S16**. Ingenuity Pathway Analysis of differentially expressed genes in fibroblasts, SMCs, and subclusters of SMCs identified in scRNA-seq of aortic cells

**Table S17.** Top 20 marker genes for each cluster identified in scRNA-seq of aortic $CD45^+$ cells

**Table S18**. Proportion/cell number of macrophage subclusters identified in scRNA-seq of aortic $CD45^+$ cells

**Table S19.** Differential expression analysis for macrophages and each macrophage subcluster ($GFP^+$ vs. $GFP^-$) identified in scRNA-seq of aortic $CD45^+$ cells

**Table S20.** Ingenuity Pathway Analysis of differentially expressed genes in macrophages and each macrophage subcluster ($GFP^+$ vs. $GFP^-$) identified in scRNA-seq of aortic $CD45^+$ cells

### SUPPLEMENTAL METHODS

#### Integrative genomics analyses (IGA)

The IGA pipeline incorporated two data sources: genome-wide association study (GWAS) summary statistics, including the UK BioBank (UKBB)(1) or Coronary Artery Disease Genome Wide Replication and Meta-analysis plus the Coronary Artery Disease Genetics (CARDIoGRAMplusC4D),(2) and tissue/cell-type specific expression Quantitative Trait Loci (eQTLs) from the Genotype-Tissue Expression project (GTEx) v8,(3) Stockholm-Tartu Atherosclerosis Reverse Networks Engineering Task study (STARNET),(4) or the BLUEPRINT project.(5) This pipeline utilized methods from two broad classes: (1) Colocalization analysis; and (2) Summarized Mendelian Randomization (SMR) followed by Heterogeneity in Dependent Instruments Test (HEIDI) or MetaXcan. We intersected the results of Colocalization analysis and SMR-HEIDI analysis to prioritize the causal gene and the causal tissues/cell types at the *LIPA* risk locus.

**Colocalization analysis:** Colocalization analysis assesses the posterior probability that GWAS CAD statistics and eQTL datasets share the same causal variants.(6) Colocalization between CARDIoGRAMplusC4D GWAS loci and GTEx *LIPA* eQTLs in blood was previously performed by the GTEx Consortium using the Enrichment Estimation Aided Colocalization Analysis (ENLOC) software.(7-9) GTEx tissue gene expression levels and genotypes were processed with the deterministic approximation of posteriors (DAP) algorithm,(10) and CAD GWAS variants were split into approximately linkage disequilibrium (LD)-independent regions (± 1 Mb flanking gene body). ENLOC was run per region to compute SNP-level posterior probabilities for being causal and a regional colocalization probability (rcp) by summing up the SNP-level probabilities that the GWAS and eQTL share a genetic effect.

The colocalization analysis between GWAS statistics (CARDIoGRAMplusC4D and UKBB) and STARTNET dataset were performed using COLOC v2.3.6 in R.(6) In the analysis, only eQTL signals with a threshold below $5 \times 10^{-8}$ were considered, and the nearby genes ± 500 kb flanking the *LIPA* gene, including *LIPA, IFIT3, IFIT2, IFIT1, CH25H, FAS, ANKRD22, STMBPL1, KIF20B, and SLC16A12*, were examined. Five hypotheses were evaluated: H0, no association with either GWAS or STARNET; H1, association with GWAS but not STARNET; H2, association with STARNET but not GWAS; H3, multiple SNPs associate with either GWAS or STARNET independently; H4, shared SNP(s) associate with both GWAS and STARNET.(11) We consider posterior probability (PP.H4) ≥ 0.8 as evidence that CAD risk and gene expression of a given tissue/cell type were mediated by the same genetic variant.

**SMR and HEIDI:** The SMR-HEIDI approach was built to test if the same genetic variant affects gene expression and complex traits. SMR unitizes summary-level data from multiple cohorts of GWAS and eQTL to identify genes whose expression are associated with a phenotype result from either a pleiotropic model (where the same variant influence both gene expression and trait) or a linkage model (where two or more genetic variants in LD independently regulate gene expression and trait).(12, 13) HEIDI test distinguishes pleiotropy from linkage by employing multiple SNPs in a *cis*-eQTL region, under the null hypothesis that the gene and trait are associated because of a single causal variant. HEIDI (P >= 0.05) indicates that we could not reject the null hypothesis that there is a single causal variant affecting both gene expression and a trait (disease risk). The analyses were conducted in the region flanking 500 kb on either side of the neary genes, including *LIPA, IFIT3, IFIT2, IFIT1, CH25H, FAS, ANKRD22, STMBPL1, KIF20B, and SLC16A12*, using the methods published by Zhu *et al* https://yanglab.westlake.edu.cn/software/smr/#Overview.(13) Associations passed both SMR ($P_{SMR}$ < 0.001) and HEIDI ($P_{HEIDI}$ >= 0.05) tests were considered significant.

**MetaXcan**: MetaXcan combines information from GWASs and eQTL data to estimate how much the expression of specific genes might influence the risk of developing certain diseases or traits. MetaXcan was applied to integrate each pair of the GWAS summary statistics and the eQTL dataset of a given STARNET tissue/cell type to identify putative causal genes for CAD. The prediction models were built using methods published by the PrediXcan/MetaXcan authors which are available at https://github.com/hakyimlab/PredictDB-Tutorial.(14) The analyses were conducted in the region flanking 1 Mb on the either side of the *LIPA* gene. P value < 5E-8 was considered as significant.

**Genotyping of human samples and genotype imputation**

All study protocols were approved by the Human Subjects Research Institutional Review Board at Columbia University (AAAQ6510). Blood from healthy subjects were obtained for the isolation of buffy coat. DNA extracted from buffy coat was genotyped using Illumina Infinium Multi-Ethnic Global Array. The following filters were applied to variants and individuals. First, we excluded variants with > 10% missing rate and individuals with > 5% genotyping missing rate. We then estimated identity by descent using variants with > 5% minor allele frequency (MAF) and identified pairs of related individuals (PI_HAT > 0.05). Only one sample with the highest genotyping rate from each group of related individuals were included for downstream analysis. Variants with Hardy-Weinberg test $P < 1\times10^{-6}$ in Caucasian population were excluded from analysis. We then calculated variant genotyping rate again and excluded variants with > 5% missing rate. Duplicate variants were also excluded from analysis. The above filtering was performed using PLINK v1.07 and custom scripts.(15) A total of 1,905,967 variants in 319 individuals were included after filtering.(16)

The filtered genotype data were split into a subset of 175 Caucasians and a subset of 79 African Americans.(16) Only variants within an arbitrary region (hg19_dna range = chr10:29801728 - 105338344) that covered *LIPA* locus were used for imputation. Pre-imputation checks were performed following recommendations from Michigan Imputation Server.(17) Briefly, imputation check script v4.3.0 and Haplotype Reference Consortium (HRC) reference panel v1.1 were downloaded from McCarthy Group Tools (https://www.well.ox.ac.uk/~wrayner/tools/). The imputation check script was run on the Caucasian genotype data and the African American genotype data to check strand, alleles, reference/alternate allele assignment, and frequency differences from the reference panel. A total of 28,494 variants from 175 Caucasians and 24,829 variants from 79 African Americans passed quality control and were submitted for genotype imputation on Michigan Imputation Server. HRC reference panel was used for both datasets. “EUR” population was selected for Caucasian data and “Other/Mixed” population was selected for African American data. Other options were kept as default.

**Isolation of human peripheral blood mononuclear cells (PBMCs)**

**Isolation of PBMCs from whole blood and cryopreservation:** All study protocols were approved by the Human Subjects Research Institutional Review Board at Columbia University (AAAQ6510). Blood from healthy subjects were obtained for the isolation of peripheral blood mononuclear cells (PBMCs). The BD VACUTAINER® Mononuclear Cell Preparation Tube (CPT™) with Sodium Citrate (BD, Franklin Lakes, NJ) for the separation of mononuclear cells from whole blood was used for blood collection using the standard technique for BD Vacutainer™. After collection, the tube was stored upright at room temperature (RT) and was processed within 2 h. Tubes were centrifuged at 1,800 × g for 30 minutes at RT. After centrifugation, lymphocyte and monocyte band (PBMC layer) was collected using a serological pipette, and was washed with ample amount of Dulbecco's Phosphate Buffered Saline (PBS, modified, without calcium chloride and magnesium chloride) followed by centrifuging at 330 × g for 10 min at RT. PBMCs were either freshly cultured and differentiate to HMDMs, or cryopreserved in freezing media (90% FBS + 10% DMSO) at 2-3 × $10^6$ cells/ml freezing media was stored in liquid nitrogen vapor phase for subsequent recovery and differentiation.

**Isolation of PBMCs from buffy coat:** Buffy coat products are purchased from New York Blood Center and diluted with three volumes of PBS supplemented with 2 mM EDTA. The diluted buffy coat was carefully laid on top of Ficoll (Sigma, GE17-5442-02) at a 4:3 ratio and centrifuged at 400 × g for 40 min at 20°C without brake. The PBMC layer was collected and washed twice with PBS containing 2% FBS, 5 mM EDTA, 20 mM HEPES, and 1 mM sodium pyruvate.

**Differentiation of human peripheral blood monocyte-derived macrophage (HMDM)**

Differentiation of human monocyte-derived macrophages (HMDMs) from PBMCs was performed as described.(18) Freshly isolated PBMC were plated (6-well plates, 2 × $10^6$ cells/well; 24-well plates, 1 × $10^6$ cells/well) differentiated into macrophages in 2 mL of RPMI medium supplemented with 20% FBS and 100 ng/µL human recombinant M-CSF (Goldbio, 120-09-100) for 7 days. Differentiated HMDMs were then used for qRT-qPCR, Western blot and enzymatic activity assays as described below.

**High-resolution Tri-HiC**

**Tri-HiC library construction:** Limited resolution of existing HiC methods resulted in vague interpretation of interplays in topologically associating domains and restriction site scanty regions. Tri-HiC combined the application of three restriction enzymes resulting in up to 100 bp fragments, thus reducing gaps of undigested intervals and improving detection of promoter-enhancer interactions. Human PBMCs were isolated from buffy coats using Ficoll reagents based on manufacturer's instruction. $CD14^+$ Monocytes and $CD3^+$ T-cells were sorted with BD FACSAria cell sorter (BD Biosciences). The Tri-HiC was conducted as previously described.(19) Briefly, 1 - 3 × $10^5$ of monocytes or 5 × $10^5$ - 1 × $10^6$ of T cells were fixed with 1% (v/v) formaldehyde (Thermo Scientific, 28908) in PBS for 10 min at room temperature (RT). Cells were washed twice with cold PBS, pelleted (300 × g, 4 min, 4°C), then treated with 250 µL cold lysis buffer (10 mM Tris-HCl pH 8.0, 10 mM NaCl, 0.2% IGEPAL CA630 (Sigma, I8896), and 50 µL protease inhibitors (Sigma, P8340). The resulting crude nuclei were permeabilized in 200 µL lysis buffer containing 0.5% (v/v) Triton X-100 for 15 min at RT. Subsequently, a triple digestion was carried out by incubating the nuclei in a 250 µL mixture containing 0.1% SDS, 1X Cutsmart buffer (NEB, B6004), 10 µL MboI (NEB,

R0147S), 5 µL CviQI (NEB. R0639S), and 5 µL CviAII (NEB, R0640S) at RT for 2 h, followed by 2 h at 37°C. Restriction digestion was then heat inactivated by incubating at 65°C for 20 min. For end blunting, a 50 µL master mix comprising 1.5 µL of 10mM CTP, GTP, and TTP, 37.5 µL of 0.4 mM biotin-dATP (Thermo Scientific, 19524016), and 4 µL of DNA polymerase I Klenow (NEB, M0210S) was added and incubated at 37°C for 1 h with rotation. Blunt end ligation was performed by mixing the suspension with a 200 µL master mix containing 50 µL of 10X T4 ligase buffer (NEB, B0202S), 2.5 µL of 20 mg/mL BSA, and 5 µL of 400 U/µL T4 ligase (NEB, M0202), and incubating under alternating temperature conditions (2 h RT, overnight at 4°C, then 2 h RT). The processed nuclei were de-crosslinked in 800 µL 1X T4 ligase buffer containing 500 mM NaCl and 1 mg proteinase K (Invitrogen, 25530049) at 68°C for 2 h, purified by phenol-chloroform extraction, then resuspended in 50 µL 10 mM Tris-HCl for library construction.

Purified DNA (300 ng) was tagmented by incubating with 20 µL TDE1 (Illumina, 20034197) in 100 µL 1X TD buffer at 55°C for 10 min, then purified with DNA Clean & Concentrator-5 (Zymo Research, D4014). The elute was mixed with 2X NEBNext high fidelity PCR master mix (NEB, M0541S), 1.25 µM ATAC-seq primers(20) in a final volume of 100 µL, then amplified by a 2-cycle PCR with cycling condition of 72°C for 5 min, 2 cycles of 98°C for 10 s, 63°C for 10 s, 72°C for 1 min, and final extension at 72°C for 5 min. The PCR product was then incubated with 1.5 mg/mL of prewashed Streptavidin C1 Dynabeads (Thermo Scientific, 65001) in 100 µL of 2X Binding & Washing Buffer (10 mM Tris-HCl pH 7.5, 1 mM EDTA, 2 M NaCl) for 15 min at RT with rotation. Beads were separated and washed twice in a magnet with 500 µL of Binding & Washing Buffer, and once with 500 µL 10 mM Tris-HCl. Beads were then resuspended in 100 µL of PCR solution containing 2X NEBNext high fidelity PCR master mix and 0.5 µM ATAC-seq primers. A following PCR reaction was performed with the cycling condition of 98°C for 30 s, 8 cycles of 98°C for 10 s, 63°C for 10 s, 72°C for 1 min, and final extension at 72°C for 5 min. The amplified library in the supernatant was purified and size-selected using 0.55-1 X SPRI beads. Samples were sequenced for 800 million reads on average on the Illumina Nova-Seq S4 platform (2 $\times$ 100 bp pair-end).

**Tri-HiC data analysis:** The raw Fastq files were aligned to the hg38 genome (GCA_000001405.15) and processed using the Pairtools with default parameters.(21) The obtained interaction pair tags were processed by Juicer to generate the interaction contact matrix in .hic format.(22) Alignments with MAPQ > 30 were subjected to downstream analysis. Juicebox was used to visualize the contact matrices.(22) Chromatin interaction loops were called by using the HiCCUPS algorithm with the following parameters: -r 500,1000,2000,5000,10000 -f 0.1 -p 4,2,2,2,2 -i 20,10,10,6,6 -t 0.1,1.25,1.75,2 -d 2000,2000,4000,10000,20000.(23)

**Independent single-nucleotide polymorphism (SNP) effects analysis**

To disentangle the effects of variants in high LD, we used various linear regression models to estimate the SNPs' independent effects on both *LIPA* gene expression and coronary artery disease (CAD). Since rs1412444 and rs1412445 were in almost perfect LD ($r^2 = 0.99$) in both GTEx and CARDIoGRAMplusC4D cohorts, we included only rs1412445 in our models ($G_P$) as well as rs1320496 ($G_G$).
For *LIPA* eQTL analysis, we fit four linear models:

rs1412445 only: $$E = \beta_R G_R + \beta_C cov + \varepsilon \quad (1)$$

rs1320496 only: $$E = \beta_G G_G + \beta_C cov + \varepsilon \quad (2)$$

Interaction model: $$E = \beta_R G_R + \beta_G G_G + \beta_{RG} G_R * G_G + \beta_C cov + \varepsilon \quad (3)$$

Conditional model: $$E - \beta_R^1 * G_R = \beta_G G_G + \beta_C cov + \varepsilon \quad (4)$$
where E is $\log_2$-transformed *LIPA* gene count data normalized with DESeq2; $G_P$ and $G_G$ are alternative allele dosages for rs1412444/rs1412445 and rs1320496, respectively; and *cov* are the covariates used by GTEx for eQTL discovery including genotype, sequencing information, donor sex, and Probabilistic Estimation of Expression Residuals (PEER) method factors.(8, 24) For the conditional model, we used $\beta_P$ from the rs1412445 only model to remove the effect of rs1412445 on expression, and a linear model using rs1320496 was fit to the residual expression.

For CAD GWAS analysis, we used individual SNP beta estimates and P values as reported in the summary statistics. We calculated the conditional effect of rs1320496 on CAD (conditional on rs1412445) using the cojo tool from the GCTA software package.(25, 26) GCTA was run with CAD GWAS summary statistics, GTEx genotypes for LD structure, and rs1412445 as a conditional SNP.

**Allelic-specific binding (ASB) and motif analysis**

The motif for PU.1, which is encoded by the *SPI1* gene, was derived from the Encyclopedia of DNA Elements (ENCODE) datasets.(27) PU.1 ASB at rs1412445 was first discovered in AlleleDB.(28) However, rs1320496 was not evaluated within AlleleDB due to its absence in the 1000 Genomes Project. We found that the GM12891 LCL cell line ENCODE PU.1 ChIP-seq sample was heterozygous for all three SNPs of interest, including rs1412445, rs1320496, and rs1412444.(29) Therefore, we used the Integrative Genomics Viewer to count the reads aligned to each allele of each SNP.(30) Motif-based binding prediction was performed by comparing both risk allele (T) and non-risk allele (C) of three SNPs to the PU.1 TF motif from ENCODE.(27)

**Dual-luciferase reporter assays and site-directed mutagenesis**

**Maintenance of cell lines:** The human embryonic kidney HEK293 (ATCC, CRL-1573) cells were cultured in DMEM medium supplemented with 10% FBS. Human THP-1 (ATCC, TIB-202) monocytic cells were maintained in RPMI medium supplemented with 10% FBS, 1 mM sodium pyruvate, 10 mM HEPES buffer, 50 µM 2-Mercaptoethanol and 2 mM L-glutamine. All cell lines were confirmed to be mycoplasma-free by MycoAlert Mycoplasma Detection Kit (Lonza, LT07-418).

**Plasmids construction:** Two putative enhancer regions surrounding SNPs rs1412444 (the rs1412444 region, hg19_dna range = chr10:91002499 - 91003138) and rs2246833 (the rs2246833 region, hg19_dna range = chr10:91005571 - 91006002) were amplified from human genomic DNA using PCR primers: rs1412444 region sense 5’-ATATATGGTACCTTTCTGTTAGTATACGAGGAGCC, rs1412444 region anti-sense 5’-ATATATCTCGAGCAGTGGGGAGTCTTCAAGGA, rs2246833 region sense 5’-ATATATGGTACCCAGTCTCCCACATTAACAAGCA, and rs2246833 region anti-sense 5’-ATATATCTCGAGCCAGCAGGGGATCTCTCAAA. The amplified fragment was introduced into KpnI and XhoI restriction sites of pGL4.23 vectors (Promega, E8411) to construct plasmids containing the rs1412444 region (pGL4.23-rs1412444) or the rs2246833 region (pGL4.23-rs2246833).

**Site-directed mutagenesis:** Non-risk allele (C) of SNPs rs1412444 and rs1412445 were replaced to risk alleles (T) by using QuickChange II XL Site-Directed Mutagenesis Kit (Agilent, 200521) in the rs1412444 region. Risk allele T of SNP rs1320496 was introduced by overlap extension PCR. The PCR primers including:
rs1412445-T, sense 5'-GGTCATTAGGAGGATGTTGGTGCTATTAATAATAGAGGAGG,
rs1412445-T, anti-sense 5'-CCTCCTCTATTATTAATAGCACCAACATCCTCCTAATGACC,
rs1320496-T, sense1 5'-ATATAT GGTACCTTTCTGTTAGTATACGAGGAGCC,
rs1320496-T, anti-sense1 CATGCATCCCACTTCCCCTCC,
rs1320496-T, sense2 5'-GGAGGGGAAGTGGGATGCATG,
rs1320496-T, anti-sense2, 5'-ATATATCTCGAG CAGTGGGGAGTCTTCAAGGA,
rs1412444-T, sense 5'-GCCTTTAAACACTGGAAATAACACCAGTGGC,
rs1412444-T, anti-sense 5'- GCCACTGGTGTTATTTCCAGTGTTTAAAGGC.
Plasmids that resembled three haplotypes in human subjects were constructed: pGL4.23-CCC represented plasmids with non-risk alleles (C) of all three SNPs; pGL4.23-CTC referred to plasmids containing risk allele (T) of SNP rs1320496; and pGL4.23-TTT represented plasmids containing risk alleles (T) of all three SNPs in rs1412444 region. All constructs were verified by Sanger sequencing.

**Dual-luciferase reporter assay:** THP-1 monocytes and HEK293 cells were transfected using the Lonza Nucleofector system with program FF-100 (Lonza, V4XC-3024). For each transfection, $1\times10^6$ cells were suspended in 100 μL of Nucleofector buffer, and 500 ng of pGL4.23-based test vectors along with 20 ng of pGL4.73 *Renilla* control vector (Promega, E6911) were added. Luciferase activity was measured 24-h post-electroporation using the Dual-Luciferase Assay System (Promega, E1960), and the results were normalized to *Renilla* luciferase activity.

**Electrophoretic mobility shift assay (EMSA)**

To assess ASB of PU.1 to different SNP variants, EMSA was performed according to the manufacturer's instruction (Thermo Scientific, 20148). The assay aimed to determine the differential binding affinity of PU.1 to biotinylated DNA probes (~ 40 bp) containing either the non-risk allele (C) or the risk allele (T) at the SNPs rs1412445, rs1320496, and rs1412444. For each assay, 20 pmol of biotin-labeled probes were incubated with 100 ng of recombinant human PU.1 protein (Sino Biological, 17186-H07E) at RT for 30 min in a buffer composed of 10 mM Tris, 50 mM KCl, 1 mM DTT, 5% glycerol, 100 mM $MgCl_2$, 1% NP-40, and 2 μg/μL polydA:dT. Competitive binding assays were performed under the same conditions, with an excess (100-fold) of unlabeled cold probes added prior to the biotin-labeled probe. For supershift assay, 2 μg PU.1 antibody (Santa Cruz, sc-390405X) was incubated with the recombinant PU.1 protein for 1 h on ice prior to the addition of the biotin-labeled probe. Free and bound probes were then resolved on a 6% DNA retardation gel (Thermo Scientific, EC63652BOX) run in 0.5X Tris-borate-EDTA buffer (Thermo Scientific, J63487.K3), followed by transferred to a Nylon membrane (Thermo Scientific, 77016) for imaging using the GE AI600 Imager. The sequence of DNA probe used are:
rs1412445-C: GGTCATTAGGAGGATGCTGGTGCTATTAATAATAGAG,
rs1412445-T: GGTCATTAGGAGGATGTTGGTGCTATTAATAATAGAG,
rs1320496-C: CAGGAGGGGAAGCGGGATGCATGAGAAGAGGACGCGTTTC,
rs1320496-T: CAGGAGGGGAAGTGGGATGCATGAGAAGAGGACGCGTTTC,
rs1412444-C: CTATTTGCCTTTAAACACCGGAAATAACACCAGTGGC,
rs1412444-T: CTATTTGCCTTTAAACACTGGAAATAACACCAGTGGC.

The non-risk alleles (C) and risk alleles (T) were highlighted in red.

**CRISPR interference targeting the rs1412444 enhancer region**

**Lentivirus packaging:** HEK239T (ATCC, CRL-3216) cells were maintained in DMEM medium supplemented with 10% FBS. Lentivirus particles were generated by co-transfecting HEK293T cells with the standard packaging plasmids pMD2G (Addgene, 12259) and psPAX2 (Addgene, 12260), along with either the dCas9-KRAB expression plasmid (Addgene, 60954) or the CRISPRia-v2 plasmid for single guide RNAs (sgRNAs) (Addgene, 84832), using Lipofectamine 2000 (Thermo Scientific, 11668027). Viral supernatant was harvested 48 - 72 h following transfection and filtered through a 0.45 µm SFCA syringe filter.

**Establishing CRISPRi THP-1 cells**: To establish the CRISPRi cell line, THP-1 cells were transduced with lentiviral constructs expressing dCas9-KRAB in RPMI medium supplemented with 10% FBS and 10 µg/mL polybrene. Polyclonal THP-1 cells stably expressing dCas9-KRAB were enriched by cell sorting using a BD FACSAria cell sorter (BD Biosciences), selecting for cells with stable mCherry expression.

**CRISPRi editing**: The sgRNAs targeting the rs1412444 enhancer region (hg19_dna range = chr10:91002499 - 91003138) were chosen based on their rankings in CRISPRick, with priority given to sgRNAs that span the SNPs rs1412445 and rs1320496. The SNP rs1412444 was deemed untargetable in the current setup due to lack of PAM sequence. The design of sgRNAs were illustrated schematically in Figure 2D. Briefly, sgRNA1 is located 68 upstream of rs1412445 and is the highest scored sgRNA within the region. The sgRNA2 spans the SNP rs1320496, and sgRNA3 spans the SNP rs1412445. The sequences of the non-targeting sgRNA and sgRNA targeting the transcription start site (TSS) of *LIPA* were obtained from previous study.(31) The sgRNAs were cloned into the CRISPRia-v2 vector, which was digested with BstXI (NEB, R0113S) and BlpI (NEB, R0585S). Sorted CRISPRi THP-1 cells were then virally transduced with constructs expressing the sgRNAs. Cells stably expressing both mCherry and BFP were enriched by sorting on a BD FACSAria cell sorter (BD Biosciences) and subsequently used in downstream experiments. The sequence of sgRNAs used are listed as: sgRNA1, GAACTGAAGCAATTTAACGT; sgRNA2: GAAGCGGGATGCATGAGAAG; sgRNA3: GCTGGTGCTATTAATAATAG; non-target (NONT): GGAGTTAAGGCCTCGTCTAG; and sgRNA *Lipa-TSS*: GTCGCAGTGCCAGCTCTCAG.

**siRNA-mediated knockdown**

THP-1 cells were seeded at a density of $1 \times 10^6$ cells/mL in a non-TC treated 6-well plate. For each well, 6 µL of lipofectamine RNAiMAX (Thermo Scientific, 13778100) was diluted in 100 µL of OptiMEM (Gibco, 31985062). Separately, 80 pmol siRNA was diluted in 100 µL of Opti-MEM, and then combined with the diluted Lipofectamin RNAiMAX and incubated at RT for 10 min to allow complex formation. The siRNA-lipid complex was then added dropwise to the cells. Cells were incubated under standard condition, and 48-h post-transfection, cells were harvested for RNA extraction and dual-luciferase reporter assay. Pre-designed siRNAs targeting *SPI1* (s13352), *STAT1* (s277 and s278), and the negative control (4390843) were purchased from Thermo Scientific.

**Experimental animals**

All animal protocols were approved by the Institutional Animal Care and Use Committee at Columbia University (AABN5560). Mice were cared for according to the NIH guidelines.

**Generation of *Lipa*$^{KI/WT}$ transgenic mice:** Mice for conditional overexpression of *Lipa* (NM_001111100) were generated by *Rosa26* knock-in (KI) of CAG-loxP-STOP-loxP-Lipa-IRES-eGFP at Columbia University Irving Medical Center Transgenic Mice Core Facility. Briefly, the construction incorporated a short 1.09 kb *Rosa26* genomic sequence upstream of the cassette and a longer 4.34 kb homologous arm downstream of the cassette, which aided in integration.(32) The constructed vectors were then electroporated into FL19 (albino/agouti C57BL/6N) embryonic stem cells. Positive clones exhibiting eGFP fluorescence were confirmed, expanded, and injected into blastocysts from C57BL/6J mice. These blastocysts were subsequently implanted into pseudopregnant female mice. Resulting male chimeras had high percentage of albino/agouti coat color were crossbred with C57BL/6J females to obtain germline transmission. The appearance of agouti pups indicated the successful germline transmission. Among agouti pups, *Lipa*$^{KI/WT}$ mice were genotyped by PCR with primers: WT, sense 5'-AAGGGAGCTGCAGTGGAGTA, anti-sense 5'-CCATGTGGCTCAATAATGAAA; and KI, sense 5'-AGCCATACCACATTTGTAGAGG, anti-sense 5'-CTTCAGGATCCACAGCTGATAC. *Lipa*$^{KI/WT}$ mice were crossbred with C57BL/6J mice to reintroduce the black fur colorization.

**Myeloid-specific *Lipa* overexpression**: To achieve myeloid-specific overexpression, *Lipa*$^{KI/KI}$ mice were bred with *LysMCre*$^{+/-}$ mice to delete the *loxP-STOP-loxP* cassette, thereby enabling overexpression of *Lipa* in myeloid cells in *LysMCre*$^{+/-}$, *Lipa*$^{KI/WT}$ mice. Littermates with the genotype *LysMCre*$^{+/-}$, *Lipa*$^{KI/WT}$ served as controls without overexpression. To assess the impact of myeloid-specific overexpression of *Lipa* on atherosclerosis, these mice were bred on an *Ldlr*$^{-/-}$ background. *Ctrl* (*LysMCre*$^{-/-}$, *Lipa*$^{KI/WT}$, *Ldlr*$^{-/-}$) and *M-Lipa*$^{KI}$ (*LysMCre*$^{+/-}$, *Lipa*$^{KI/WT}$, *Ldlr*$^{-/-}$) mice were fed a Western diet (WD, TD88137, Harlan Teklad) or normal laboratory diet (ND, PicoLab Rodent Diet 20, 5053) as indicated in different experiments. Mice were socially housed in standard cages at 22°C under a 12 - 12 h light-dark cycle with ad libitum access to water and food in the barrier facility.

Age- and sex-matched littermates, including both males and females, were included in the study. Mice died unexpectedly during the study for reasons unrelated to the experimental procedures, and mice that required euthanasia prior to the endpoint were excluded from the study. Data from mice where there were technical failures, including equipment malfunction, were excluded from the analyses. Mice were grouped by genotype thus not randomized. For blood biochemical measurement and RNA-seq library preparation, the technicians were blinded from experimental design. Imaging and histological data were also analyzed blindly. For other experiments, investigators were not blinded to mouse genotypes during data collection and analysis to ensure that age/sex-matched pairs were used for each genotype and in each independent experiment.

**Bone marrow-derived macrophage (BMDM) differentiation and culture**

Bone marrow cells were flushed out from tibias and femurs with RPMI medium and filtered through 40 μm cell strainers. Bone marrow cells were washed twice with RPMI medium by centrifuging at 300 × g for 5 min at 4°C. The collected cells were cultured in DMEM supplemented with 10% FBS, 20% L929-cell conditional medium, and 2 mM L-glutamine. Fresh medium was replenished every 2 - 3 days during differentiation. Assays were performed in BMDMs from day 7 to day 10.

**Peritoneal macrophage (PM) isolation and culture**

Mice were euthanized according to approved IACUC protocol. Ice-cold PBS (10 mL) supplemented with 10% FBS and 2 mM EDTA was injected into the abdominal cavity. After gentle massage of the peritoneum for 5 min, peritoneal lavage was collected using a 10 mL syringe with 25G needle. Peritoneal cells were pelleted at 300 × g for 5 min and stained with F4/80-PE (BioLegend 123110, clone BM8, 10 ng/µL) in 400 µL of FACS buffer for 10 min on ice. The F4/80$^+$ PMs from *Ctrl* mice and F4/80$^+$GFP$^+$ PMs from *M-Lipa$^{KI}$* mice were collected using the BD Influx cell sorter (BD Biosciences). PMs sorted from mice fed an ND (8-12 weeks old) were cultured in DMEM supplemented with 10% FBS, 100 ng/µL M-CSF, and 2 mM L-glutamine for 48 h before the indicated assays. PMs sorted from mice fed a WD for 16 weeks were subjected to the indicated assays without culturing.

**Quantitative reverse-transcription PCR (qRT-PCR)**

Total RNA was extracted with Quick-RNA Miniprep kit (Zymo Research, R1055) and cDNA was synthesized using High-Capacity cDNA Reverse Transcription Kit (Applied Biosystems, 4368813). Real-time qPCR was performed using either SYBR Green Master Mix (Thermo Scientific, A25742) or TaqMan Fast Advanced Master Mix (Thermo Scientific, 4444557) in a 10 µL reaction volume on the QuantStudio 7 Flex Real-Time PCR System (Applied Biosystems, 4485701). Analysis of gene expression was determined using the $2^{-\Delta\Delta Ct}$ method with *Actb* as the reference gene unless otherwise specified. For human samples, the following primers were used: *LIPA*, sense 5'-CTAGAATCTGCCAGCAAGCC, anti-sense 5'-TGTGCCTTAACCGAATTCCT; *ACTB*, sense 5'-AGAGCTACGAGCTGCCTGAC, anti-sense 5'-AGCACTGTGTTGGCGTACAG. For mouse samples, the following probes were used: *Abca1* (Mm00442646_m1), *Abcg1* (Mm00437390_m1), *Actb* (Mm02619580_g1), *Ccl2* (Mm00441242_m1), *Ccl7* (Mm00443113_m1), *Ccl8* (Mm01297183_m1), *Ccl12* (Mm01617100_m1), *Cd36* (Mm00432403_m1), *Cyp5a1* (Mm00490968_m1), *Il1b* (Mm00434228_m1), *Il18* (Mm00434225_m1), *Lipa* (Mm00498820_m1), *Lss* (Mm00461312_m1), *Msr1* (Mm00446214_m1), *Sc5d* (Mm00555295_m1), *Hsd17b7* (Mm00501703_m1), and *Olr1* (Mm00454582_m1).

**Lysosomal acid lipase activity assay**

LIPA enzyme activity was measured using fluorogenic substrate 4-methylumbelliferyl-oleate (4-MUO, Sigma, 75164), as previously described.(33) Briefly, cells were seeded on a 6-well plate and lysed with 100 µL RIPA buffer (Sigma, R0278) containing protease inhibitor (Sigma, 11697498001) on ice for 30 min. The lysate was centrifuged at 12,000 × g for 20 min at 4°C, and the supernatant was collected. A stock solution of 4-MUO was prepared in DMSO at a concentration of 100 mg/mL and diluted 100-fold with 4% Triton X -100 immediately before the assays. In each reaction, protein lysate (~10 µg) was combined with 140 µL of 200 µM sodium acetate buffer (pH = 5.5) and 50 µL of the diluted 4-MUO substrate, then incubated at 37°C for 30 min. The reactions were stopped by adding 100 µL of 1M Tris-

HCl (pH = 8.0, Thermo Scientific, 15568025). A 200 µL aliquot of the final reaction mixture was transferred to a black 96-well polystyrene plate (Corning, 3915) and read using a BioTek Microplate Reader with excitation at 360 nm and emission at 460 nm. Protein concentration was measured using a BCA assay (Thermo Scientific, PI23228). Enzyme activities were normalized to protein concentration and reported as nmol/mg of cell lysate/hr.

**Western blot**

Cells were seeded on 6-well plates and lysed with 100 µL RIPA buffer supplemented with protease inhibitor. Protein concentrations were measured with BCA Assay kit. Equal amounts (15 µg) of protein lysate were diluted with 4X LDS sample buffer (Thermo Scientific, NP0007) containing 50 mM dithiothreitol (Thermo Scientific, NP0004). The mixtures were loaded onto 4 - 12% Bis-Tris gels (Thermo Scientific, NP0322BOX) and transferred to 0.45 µm nitrocellulose membranes. After blocking with 5% non-fat milk for 1 h at RT, membranes were incubated with an anti-LIPA primary antibody (for human sample, OriGene, TA309730, 1 ng/µL; for mouse sample, Proteintech, 12956-1-AP, 0.3 ng/µL) overnight at 4°C. Membranes were washed with TBS buffer (Thermo Scientific, 28358) containing 0.1% (v/v) Tween-20 for three times and then incubated with an HRP-conjugated secondary antibody (1:2000) at RT for 1 h. GAPDH-HRP (Cell Signaling, 3683S, 1 ng/µL) was used as the loading control. Protein bands were visualized using SuperSignal West Pico Chemiluminescent Substrate (Thermo Scientific, 34080) and imaged with a GE AI600 Imager. Protein expression levels were quantified using ImageJ 1.53c.

The specificity of the rabbit polyclonal LIPA antibody (OriGene, TA309730) was previously validated in macrophages differentiated with human iPSC with CRISPR-mediated knockout of *LIPA*.(33) Validation of the rabbit polyclonal LIPA antibody (Proteintech, 12956-1-AP) was provided by the vendor, using HEK 293 cells with RNAi-mediated *LIPA* knockdown, as shown on the product webpage. The selection of antibodies for human versus mouse samples was based on their differential sensitivity, with the rabbit polyclonal LIPA antibody (OriGene, TA309730) demonstrating better sensitivity in human samples, while the rabbit polyclonal LIPA antibody (Proteintech, 12956-1-AP) showed higher sensitivity in mouse samples.

**Plasma lipid profile**

Blood samples were collected from mice after a 4-h fasting via retro-orbital bleeding into EDTA-coated tubes. Plasma was obtained by centrifuging the blood at 1000 × g for 15 min at 4°C. Plasma total cholesterol and triglyceride levels were measured using enzymatic kits (cholesterol: Wako, 999-02601; triglyceride: Thermo Scientific, TR22421). To assess the distribution of cholesterol in different lipoprotein fractions, pooled plasma from sex- and age-matched mice were fractionated using Fast Protein Liquid Chromatography (FPLC). Briefly, 350 µL of pooled plasma was loaded onto a Superose 6 column (GE Healthcare) and eluted with 22 mL of elution buffer (0.15 M NaCl, 1 mM EDTA) in 500 µL fractions at a flow rate of 0.3 mL/min at 4°C. Cholesterol and triglyceride levels in each fraction were determined using the enzymatic kits as specified above.

**Mouse complete blood cell count and differential count**

Blood samples (~100 µL/mouse) were collected into EDTA-coated tubes (BD, 366643) via retro-orbital bleeding, then subjected to a complete blood count (CBC) with differential analysis using a Heska Element HT5 by the diagnostic lab at the Institute of Comparative Medicine at Columbia University Irvine Medical Center.

**Flow cytometry of blood, spleen and bone marrow (BM) cells**

Blood samples were collected into EDTA-coated tubes via retro-orbital bleeding from mice. Spleens were gently meshed using a syringe plunger and passed through 40 µm cell strainers. Red blood cells (RBCs) in the blood, spleen, and bone marrow (BM) samples were lysed with cold RBC lysis buffer (BioLegend, 420301) on ice for 5 min. Cell suspensions were incubated with anti-mouse CD16/32 (BioLegend 101302, clone 93, 10 ng/µL) in FACS buffer (PBS supplemented with 2% FBS, 20 mM HEPEs and 5 mM EDTA) on ice for 10 min to block nonspecific binding of antibodies to Fc receptors. To evaluate the percentage of $GFP^+$ cells within the $CD115^+$, $Ly6G^{hi}$, and $CD3^+$ population, the following panel was used: CD45.2-PE/Cy7 (BioLegend 109830, clone 104, 2.5 ng/µL), CD115-APC (BioLegend 135509, clone AFS98, 2.5 ng/µL), Ly6G-PE (BioLegend 127607, clone 1A8, 2.5 ng/µL), Ly6C-PerCP/Cy5.5 (BioLegend 128011, Clone HK1.4, 2.5 ng/µL), and CD3-BV650 (BioLegend 100229, clone 17A2, 10 ng/µL). To assess the percentage of leukocytes in BM, the following panel was used: CD45.2-PE/Cy7, CD115-APC, and Gr1-PE (BioLegend 108407, Clone RB6-8C5, 2.5 ng/µL). To assess the percentage of progenitor cells in BM, the cells were incubated with CD16/32-APC/Cy7 (BioLegend 101327, clone 93, 2.5 ng/µL) for 10 min on ice in dark, then stained with following antibodies: Lineage antibody cocktail-PerCP/Cy5.5 (BD Biosciences 561317, 20 ng/µL), cKit-PE/Cy7 (BioLegend 105813, clone 2B8, 2.5 ng/µL), Sca1-BV711 (BD Biosciences 563992, clone D7, 4 ng/µL), and CD34-AF647 (BD Biosciences 560230, Clone RAM34, 4 ng/µL) for 15 min on ice. Data were acquired on the BD LSRII and analyzed with FCS Express 7.

**Chemokine and cytokine quantification by flow cytometry-based multiplex immunoassays**

Blood was collected from mice fed a WD for 15 weeks. Chemokines and cytokines in the plasma were assessed using a customized LEGENDplex 13-plex kit (LEGENDplex, BioLegend) following the manufacturer's instruction. Plasma samples were mixed with beads coated with capture antibodies specific for CXCL1, TNFα, CCL24, CCL8, IL18, CCL22, CX3CL, CCL7, TGFβ, IL6, IFNβ, CCL2, and IL1β, and incubated on a 96-well filter plate for 2 h. Beads were washed and incubated with biotin-labeled detection antibodies for 1 h, followed by a final incubation with streptavidin-PE for 30 min. Beads were analyzed using a Bio-Rad ZE5 Cell Analyzer. Analysis was performed using the LEGENDplex analysis software (BioLegend), which distinguishes between the 13 different analytes on the basis of bead size and internal dye.

**Atherosclerotic lesion analysis**

**Preparation of serial sections of the aortic root**: After 16 weeks on WD, mice were euthanized, the aortic root and the base of the heart were freshly transferred to the bottom of cryomold filled with Tissue Frozen Medium (General Data Healthcare, TFM-C) with the aortic root perpendicular to the bottom surface and freeze on dry ice for cryosectioning. When three valves of aortic sinus appeared, consecutive sections at a thickness of 10 μm were collected until the aortic wall became incomplete (Leica, CM1850). Tissue sections were placed onto Superfrost Plus Microscope Slides (Fisher, 22-037-246) sequentially, with two sections on each slide. The section that initially presented broken leaflets in all three valves was designated as 'slide 0.'

**Lesion area and necrotic core area:** Slides numbered 15, 10, and 5 prior to slide 0, as well as slide 0 itself, along with the slide numbered 5 following slide 0, were utilized for the quantification of both the lesion area and the necrotic core area. Hematoxylin & Eosin staining were performed on these five slides, which were at the same spatial latitude (mouse to mouse) and spaced 100 μm apart. The average from five slides for each mouse was used to determine the lesion size and necrotic core area. Necrotic core was defined as an area devoid of intact cells based on the latest scientific guideline from American Heart Association.(34) Images were captured with Nikon Eclipse Ti Microscope and analyzed with ImageJ 1.53c software by an observer blinded to the group assignment.

**Fibrous cap thickness:** Aortic root sections (one section per mouse) obtained from the same lesional spatial latitude were stained with Picrosirius Red per the manufacture's instruction (Polysciences, 24901-500). Cap thickness was measured on each section at even intervals 5 μm apart. Then the average thickness was reported.

**Immunofluorescence staining**: Frozen sections from the same spatial regions (mouse to mouse) were fixed in 4% (v/v) PFA for 20 min at RT, washed three times with PBS, then permeabilized with 0.2% Triton-X 100 for 20 min at RT. After blocking for 1 h in 10% goat serum (Invitrogen, 50062Z) at RT, sections were incubated overnight at 4°C with rat-anti-mouse-CD68 (Abcam, ab53444, 5 ng/μL), and rabbit-anti-mouse-Ki67 (Thermo Scientific, MA514520, 1 ng/μL) in the 10% goat serum. Slides were then washed with PBS three times and incubated with secondary antibodies (Invitrogen, A21247 and A21429) for 1.5 h at RT. TUNEL staining was performed following the instructions of manufacture (Roche, 12156792910). Slides were mounted with Prolong Gold Antifade Mountant with DAPI (Invitrogen, P36931). Images were captured with ImageXpress Micro4 Microscope and analyzed with ImageJ 1.53c software by an observer blinded to the group assignment.

**Labeling blood monocytes and assessing monocyte recruitment to atherosclerotic plaques**

Pulse-labeling of monocytes using fluorescent beads to determine monocyte recruitment to the atherosclerotic plaque was performed in *Ctrl* and *M-Lipa*$^{KI}$ mice fed a WD for 15 weeks.(35) To label Ly6C$^{hi}$ monocytes, circulating monocytes were first depleted by intravenous injection of clodronate liposome solution (Fisher Scientific, CLD-8909, 100 μL per 25 g of body weight) one day before beads injection. By depleting all circulating monocytes, including both the classical Ly6C$^{hi}$ monocytes and non-classical Ly6C$^{low}$ monocytes, the injected beads are captured by Ly6C$^{hi}$ monocytes repopulating the blood.(36, 37) To pulse label Ly6C$^{hi}$ monocytes that reappear in the circulation, Fluoresbrite Polychromatic Red-labeled latex beads (Polysciences, 18660-5, 1 μm) diluted 1:4 in sterile PBS were injected intravenously (100 μL per 25 g of body). The labeling efficiency of Ly6C$^{hi}$ monocytes was assessed using flow cytometry at 24 h after beads injection. Specifically, blood cells were blocked with CD16/32 and stained with CD45.2-BV711, CD115-APC, Gr1-PE/Cy7 (BioLegend 108415, clone RB6-8C5, 2 ng/μL) to determine the percentage of bead-positive Ly6C$^{hi}$ monocyte (Beads$^{+}$CD45$^{+}$CD115$^{+}$Gr1$^{hi}$)

in all leukocytes (CD45$^+$). Data were collected using NovoCyte flow cytometer (Agilent) and analyzed by FCS Express 7.

On day 3 after beads injection, mice were euthanized to assess monocyte recruitment to the atherosclerotic lesion. OCT-embedded frozen sections of aortic root (10 μm) were fixed with 4% PFA and stained with rat anti-CD68 antibodies to label CD68$^+$ macrophage area in the lesion as described above. The number of beads in CD68$^+$ lesion area per aortic root section was counted and normalized by the labeling efficiency of beads in the Ly6C$^{hi}$ monocytes in the blood.

**Flow cytometry-based method to identify lipid-laden foamy macrophages in the atherosclerotic aorta**

The whole aorta, including aortic arch, ascending and descending aortas, were dissected from *Ctrl* and *M-Lipa*$^{KI}$ mice fed a WD for 16 weeks and digested into single cells with enzyme cocktails as described above. Non-specific binding to the Fc receptor was blocked by incubating cells with CD16/32 for 10 min in 200 μL FACS buffer on ice. Then, the cell suspension was incubated with CD45.2-BV711 (BioLegend 109847, clone 104, 2.5 ng/μL), CD64-APC (BioLegend 139306, clone X54-5/7.1, 2.5 ng/μL), CD11b-PE/Cy7(BioLegend 101216, Clone M1/70, 2.5 ng/μL), and LipidTOX (Thermo Fisher H34476, 1:200) in 400 μL FACS buffer for 30 min on ice in dark. Cells were washed twice with FACS buffer and stained with DAPI prior to loading into flow cytometry. Foamy macrophages were characterized as DAPI$^-$CD45$^+$CD11b$^+$CD64$^+$SSC$^{hi}$LipidTOX$^{hi}$ cells. The percentage of foamy macrophages in aortic macrophages (DAPI$^-$CD45$^+$CD11b$^+$CD64$^+$) was quantified. Data were acquired using a BD LSR II flow cytometer (BD Biosciences) and analyzed with FCS Express 7.

**FACS sorting of aortic macrophages**

The whole aortas, including aortic arch, ascending and descending aortas, from *Ctrl* and *M-Lipa*$^{KI}$ mice fed a 16-week WD were dissected and processed into single cells using enzyme cocktails as described above. Non-specific binding to the Fc receptor was blocked by incubating cells with CD16/32 for 10 min in FACS buffer on ice. Cells were then stained with CD45.2-PE (BioLegend 109807, clone 104, 2.5 ng/μL), CD11b-PE/Cy7, and CD64 in 200 μL FACS buffer on ice for 15 min. Cells were stained with DAPI prior to loading into the FACSAria cell sorter (BD Biosciences). For *Ctrl* mice, aortic macrophages (DAPI$^-$CD45$^+$CD11b$^+$CD64$^+$) were directly sorted into 600 μL RNA Lysis Buffer (Zymo Research R1060-1-50). For *M-Lipa*$^{KI}$ mice, GFP$^+$ and GFP$^-$ aortic macrophages were collected separately.

**Flow cytometry-based method to identify lipid-laden foamy PMs in the peritoneum**

The peritoneal cells were collected as described above from *Ctrl* and *M-Lipa*$^{KI}$ mice fed a WD for 16 weeks. The peritoneal cells ($1\times10^6$) were stained with CD45.2-PE/Cy7 (BioLegend 109830, clone 104, 2.5 ng/μL), F4/80-APC/Cy7 (BioLegend 123117, clone BM8, 10 ng/μL), and LipidTOX (Thermo Fisher H34475, 1:200) in 400 μL FACS buffer for 30 min on ice in dark. Cells were washed twice with FACS buffer and stained with DAPI right before analysis. Foamy PMs were identified as DAPI$^-$CD45$^+$F4/80$^+$SSC$^{hi}$LipidTOX$^{hi}$ populations and the percentage of foamy PMs in PMs (DAPI$^-$CD45$^+$F4/80$^+$) was quantified. Data were acquired on a LSRII and analyzed with FCS Express 7.

**Intracellular neutral lipids staining**

PMs sorted from mice fed an ND were seeded in Chambered Coverslip (Ibidi, 80826) at the density of 150, 000 cells/well for 48 h. Cells were then loaded with 50 μg/mL of oxLDL (Thermo Scientific, L34357) for 24 h in DMEM supplemented with 2% FBS, followed by two washes with PBS. Subsequently, the cells were fixed with 4% PFA and stained with LipidTOX (1:200) for 30 min at RT in dark. Cells were imaged using a Nikon Ti Eclipse Confocal Microscope equipped with 60x/1.49 Apo TIRF oil immersion lens. Quantification was performed using Cell Profiler 4.2.4.

**Binding and uptake of Dil-labeled oxLDL**

PMs sorted from mice fed an ND were seeded in Chambered Coverslips (Ibidi 80826) at a density of 150,000 cells/well for 48 h. Cells were then treated with 10 μg/mL Dil-oxLDL (Thermo Scientific, L34358) for 45 min at 4°C to assess binding. To evaluate uptake, cells were incubated with 10 μg/mL Dil-oxLDL for 45 min at 37°C. Cells were washed with PBS twice before fixed with 4% PFA. Cells were imaged using a Nikon Ti Eclipse Confocal Microscope equipped with 60x/1.49 Apo TIRF oil immersion lens. Quantification was performed using Cell Profiler 4.2.4.

**Quantification of intracellular cholesterol**

Intracellular cholesterol levels were quantified using fluorometric assays in PMs isolated from ND-fed mice loaded with modified-LDL or PMs isolated from WD-fed mice. PMs sorted from mice fed an ND were seeded in 12-well plate at a density of 0.5 x $10^6$ cells/well and cultured for ~18 h, then loaded with 50 μg/mL oxLDL for 24 h. PMs sorted from mice fed a WD were directly used for the assay without culturing. The cholesterol content was measured using Amplex Red Cholesterol Kit (Thermo Scientific, A12216) according to the manufacturer's instruction. Briefly, to extract lipids, 1 mL of hexane/isopropanol 3:2 (v/v) was added to the cells in the plates and incubate at RT for 30 min. The extracts were then centrifuged at 21 000 g for 10 min at RT, and the resulting supernatant was dried under Nitrogen flow in glass tubes. The dried lipids were reconstituted with 200 µL of 5X reaction buffer, and 10 µL of the reconstituted lipids was used to each reaction. Free cholesterol and total cholesterol were quantified by performing reactions in absence and presence of cholesterol esterase. Esterified cholesterol was calculated by subtracting free cholesterol from the total value. After lipid extraction, cellular protein was dissolved in 100 µL of 0.2 N sodium hydroxide. Protein concentration was measured using a BCA assay. Cholesterol quantities were normalized to input protein.

**Bulk RNA-sequencing (RNA-seq) and data analysis**

**Bulk RNA-seq sample preparation and sequencing:** Total RNA was extracted using the Quick-RNA Miniprep kit (Zymo Research, R1055). Library preparation and RNA sequencing were performed by the Columbia Genome Center. For macrophages isolated from plaque samples, the SMART-Seq v4 Ultra Low Input RNA Kit for Sequencing (Takara, 634894) was used to reverse transcribe and amplify RNA from an input of 10 ng per sample. Following cDNA synthesis, libraries were prepared using the Nextera XT (Illumina, FC-131-1096). Sequencing was performed on the NovaSeq 6000 System (Illumina). Samples were multiplexed in each lane, yielding approximately 40 million 100 bp paired-end reads per sample. For PMs, mRNA was isolated using a poly-A tail pulldown method, and cDNA libraries were constructed using the TruSeq Stranded mRNA Library Prep Kit (Illumina, RS-122-2001/RS-122/2002). Sequencing was performed on the AVITI System (Element Biosciences). Samples were multiplexed in each lane, yielding approximately 20 million 75 bp paired-end reads per sample.

**Bulk RNA-seq data processing, differential expression analysis, and Gene Set Enrichment Analysis:** Raw sequencing reads were quantified with Salmon v1.10.2 to obtain transcript abundance counts by mapping to mouse genome GRCm39 GENCODE v.M33. The resulting counts were summarized to the gene level using tximport v1.28.0.(38) Differential expression was analyzed using DESeq2 v1.40.0.(39) Differentially expressed (DE) genes were defined by an absolute fold change > 1.5 and false discovery rate (FDR)-adjusted P value < 0.05 and visualized using Volcano plots. The output of DESeq2 was scored and ranked by apeglm v1.14.0(40) using ranking metrics "−log10 P value multiplied by the sign of shrunken log2 fold-change".(41) The ranked gene list was then subjected to Gene Set Enrichment Analysis (GSEA)(42) to identify the gene sets overrepresented at the top or bottom of the ranked list using pathway definition file "Mouse_GOBP_AllPathways_no_GO_iea_July_03_2023_symbol.gmt" downloaded from http://baderlab.org/GeneSets. Only ontologies with more than 15 genes and less than 200 genes were considered. According to the GSEA User Guide, the nominal p value estimates the significance of the observed enrichment score for a single gene set. The FDR is the estimated probability that a gene set with a given enrichment score (normalized for gene set size) represents a false positive finding. A reported p value of zero (0.0) indicates an actual p value of less than 1/number-of-permutations, which was set as 2,000 in our analysis as recommended.(42) The top ranked pathways were manually curated to remove the repetitive pathways and visualized using dot plots. The leading-edge genes, defined as the subset of genes within a gene set that contribute most to the enrichment score, identified in the enriched pathways were visualized using pheatmap v1.0.12.

**Single-cell RNA sequencing (scRNA-seq) and data analysis**

**scRNA-seq with aortic cells isolated from *Ctrl* and *M-Lipa*$^{KI}$ mice**: To assess the impacts of *M-Lipa*$^{KI}$ on plaque cell composition and function, aortic single cells isolated from *Ctrl* and *M-Lipa*$^{KI}$ mice fed a WD for 16 weeks were subjected to scRNAseq as described.(43) Briefly, mouse hearts were perfused in situ with cold PBS twice. The ascending aorta, aortic arch, and descending thoracic aorta were excised, cleaned of perivascular adipose tissue, and cut into small pieces to facilitate enzymatic digestion. The aortic tissues were digested into single cells in 1 mL RPMI medium containing 4 U/mL of Liberase TM (Sigma, 5401127001), 60 U/mL of hyaluronidase (Sigma, H3506), and 60 U/mL of DNase I (Worthington Biochemical Corporation, LS006333) at 37°C for 40 min. The enzymes were then neutralized by adding 4 mL of RPMI medium supplemented with 10% FBS. The obtained cell suspensions were filtered through 100 μm cell strainers and stained in 500 μL FACS buffer including DRAQ5 (BioLegend, 424101,1:1000) and DAPI (BioLegend, 422801, 1:3000) for 10 min on ice. Viable singlets (DRAQ5$^{+}$DAPI$^{-}$) were collected into DMEM/F12 medium (Cytiva SH30261.01) containing 10% FBS, and immediately submitted to the

Single Cell Analysis Core at the JP Sulzberger Columbia Genome Center. Cells from three male mice per genotype were pooled together.

Samples were prepared using the 10x Genomics Chromium Single Cell 3' Reagent Kits v2 (mouse samples) according to the manufacturer's instructions. 5,000 cells and 200 M reads per sample were targeted. FASTQ files were aligned to mouse reference genome build mm10 and filtered with Cell Ranger v.3.0.1 pipeline. A unique molecular identifier (UMI) count matrix was created for each sample and imported to Seurat 4.3.0.1 in R.(44) Cells were retained if all of the following criteria were satisfied: number of genes per cell (> 1500 and < 6000), number of UMIs (< 20000), and percentage of mitochondrial reads (< 10%). The *Ctrl* and *M-Lipa$^{KI}$* datasets were integrated using the pipeline in Seurat built on canonical correlation analysis (CCA). Integration anchors were identified based on the union of highly variable genes across datasets (2000 genes) and the first 30 dimensions from CCA. Two-dimensional Uniform Manifold Approximation and Projection (UMAP) was constructed with 25 nearest neighbors and the first 40 principal components as input. Clusters were identified with a resolution of 0.6. Cells identified as smooth muscle cells (SMCs) and macrophages were separately extracted from the dataset. Subclustering was subsequently performed on each cell type to identify distinct subtypes. In macrophages, 36 cells (0.33% of total macrophages) were excluded from visualization and analysis as they were SMC/SMC-like cells appearing in macrophage clusters or macrophages appearing in SMC/SMC-like clusters, due to apparent UMAP visualization artifacts. The count of different cell types and subclusters of macrophage and SMC cells were compared between *M-Lipa$^{KI}$* versus *Ctrl* using Chi-square test, followed by Bonferroni-adjusted pairwise comparison using the chisq.posthoc.test 0.1.2.(45) Differential expression analysis between clusters were performed using the MAST 1.26.0 test with a minimum proportion threshold (`min.pct`) of 0.1.(46) Genes with an absolute fold change > 1.2 and Bonferroni-corrected P < 0.05 were defined as differentially expressed (DE). Ingenuity Pathway Analysis (IPA, Qiagen) was performed using DE genes. Significantly altered canonical pathways were ranked based on the Z-Score.

**scRNA-seq with CD45$^{+}$ aortic cells isolated from *M-Lipa$^{KI}$* mice**: To investigate the functional and transcriptional differences between *Lipa-overexpressing* (GFP$^{+}$) and *Ctrl* (GFP$^{-}$) macrophages in response to the same atherogenic environment, CD45$^{+}$GFP$^{+}$ and CD45$^{+}$GFP$^{-}$ aortic cells that obtained from *M-Lipa$^{KI}$* mice were routed to scRNA-seq. Briefly, the aortic single-cell suspensions isolated from *M-Lipa$^{KI}$* mice fed a WD for 16 weeks were stained in 400 µL of FACS buffer containing CD45.2-APC (BioLegend 109814, clone 104, 2.5 ng/µL) and Calcein Violet (Thermo Scientific, C34858, 100 ng/mL) for 15 min on ice. Propidium Iodide (PI) solution (BioLegend, 421301, 10 ng/µL) was added immediately before sorting. Viable (Calcein Violet$^{+}$PI$^{-}$) CD45$^{+}$GFP$^{+}$ and CD45$^{+}$GFP$^{-}$ cells were collected into DMEM/F12 medium containing 10% FBS, and immediately submitted to the Single Cell Analysis Core. Cells from five male mice were pooled together.

The Fastq files from each sample were processed as described above. Cells were retained if all of the following criteria were satisfied: number of genes per cell (> 600 and < 6000), number of UMIs (< 20000), and percentage of mitochondrial reads (< 5%). The GFP$^{+}$ and GFP$^{-}$ datasets were integrated using Seurat's canonical correlation analysis pipeline, identifying integration anchors based on 3000 highly variable genes and the first 30 dimensions from CCA. Two-dimensional Uniform Manifold Approximation and Projection (UMAP) was constructed with 20 nearest neighbors and the first 40 principal components as input. Clusters were identified with a resolution of 0.4. Cell counts of each macrophage subcluster (clusters 0, 1, 2, 4, and 5) from $GFP^-$ and $GFP^+$ macrophages were compared using Chi-square test, followed by Bonferroni-adjusted pairwise comparison using the chisq.posthoc.test 0.1.2.(45) Differntial expressing genes between clusters were performed using the MAST 1.26.0 test with min.pct threshold as 0.1.(46) Genes with an absolute fold change > 1.5 and Bonferroni-corrected P < 0.05 were defined as DE genes. Ingenuity Pathway Analysis (IPA, Qiagen) were performed using DE genes. Significantly altered canonical pathways were ranked based on IPA Z-Score.

**Statistical analyses**

All statistical analyses were performed using GraphPad Prism 9 as described below, unless otherwise specified in the figure legends. When sample size (n) ⩾ 6, data were tested for normality using Shapiro-Wilk test (when n < 8) or D'Agostino-Pearson test (when n ≥ 8). F-test of equality of variances was performed to compare the two sample variances. Data that passed normality tests are presented as mean ± standard error of mean (SEM) and analyzed using two-tailed Student's t-test for comparison of two groups and equal variances (or with Welch's correction if F-test was not satisfied), or one-way ANOVA with Tukey's post-hoc tests for one independent variable with more than two groups. Data that did not follow a normal distribution were analyzed using nonparametric tests, i.e., Mann-Whitney U test or Kruskal-Wallis test, and are presented as median ± 95% confidence interval (CI). Nonparametric tests were also used when n < 6. P < 0.05 was considered statistically significant. The number of independent experiments and biological replicates is specified in the figure legends.

### SUPPLEMENTAL FIGURES AND FIGURE LEGENDS

### Figure S1

**A**

| Tissue | Samples | NES | p-value | m-value |
|---|---|---|---|---|
| Whole Blood | 670 | 0.509 | 7.2e-66 | 1.00 |
| Spleen | 227 | 0.363 | 8.4e-21 | 1.00 |
| Adipose - Visceral (Omentum) | 469 | 0.358 | 3.5e-18 | 1.00 |
| Adrenal Gland | 233 | 0.353 | 5.8e-18 | 1.00 |
| Lung | 515 | 0.288 | 1.3e-23 | 1.00 |
| Pancreas | 305 | 0.250 | 1.5e-14 | 1.00 |
| Thyroid | 574 | 0.238 | 9.6e-17 | 1.00 |
| Stomach | 324 | 0.235 | 1.0e-10 | 1.00 |
| Heart - Left Ventricle | 386 | 0.223 | 3.2e-12 | 1.00 |
| Colon - Transverse | 368 | 0.218 | 9.0e-11 | 1.00 |
| Muscle - Skeletal | 706 | 0.206 | 1.5e-11 | 1.00 |
| Colon - Sigmoid | 318 | 0.202 | 4.1e-6 | 1.00 |
| Pituitary | 237 | 0.177 | 3.2e-4 | 1.00 |
| Adipose - Subcutaneous | 581 | 0.177 | 2.0e-10 | 1.00 |
| Liver | 208 | 0.144 | 7.6e-4 | 0.971 |
| Heart - Atrial Appendage | 372 | 0.142 | 2.2e-5 | 0.975 |
| Prostate | 221 | 0.140 | 5.5e-4 | 1.00 |
| Artery - Aorta | 387 | 0.136 | 1.2e-5 | 0.972 |
| Skin - Not Sun Exposed (Suprapubic) | 517 | 0.126 | 2.5e-5 | 0.756 |
| Small Intestine - Terminal Ileum | 174 | 0.124 | 8.8e-4 | 0.646 |
| Vagina | 141 | 0.122 | 0.1 | 0.599 |
| Skin - Sun Exposed (Lower leg) | 605 | 0.121 | 6.8e-6 | 0.588 |
| Ovary | 167 | 0.110 | 0.03 | 0.334 |
| Breast - Mammary Tissue | 396 | 0.106 | 5.2e-4 | 0.194 |
| Esophagus - Gastroesophageal Junction | 330 | 0.102 | 3.2e-3 | 0.101 |
| Testis | 322 | 0.101 | 1.1e-3 | 0.0680 |
| Nerve - Tibial | 532 | 0.0955 | 2.5e-5 | 0.00 |
| Minor Salivary Gland | 144 | 0.0911 | 0.2 | 0.130 |
| Esophagus - Mucosa | 497 | 0.0834 | 0.005 | 0.00 |
| Artery - Coronary | 213 | 0.0833 | 0.07 | 0.0720 |
| Brain - Substantia nigra | 114 | 0.0771 | 0.2 | 0.0840 |
| Esophagus - Muscularis | 465 | 0.0613 | 0.04 | 0.00 |
| Uterus | 129 | 0.0612 | 0.4 | 0.0400 |
| Kidney - Cortex | 73 | 0.0397 | 0.7 | 0.205 |
| Artery - Tibial | 584 | 0.0197 | 0.4 | 0.00 |
| Brain - Nucleus accumbens (basal ganglia) | 202 | 0.00976 | 0.8 | 0.00 |
| Cells - Cultured fibroblasts | 483 | -0.00602 | 0.8 | 0.00 |
| Brain - Amygdala | 129 | -0.0141 | 0.8 | 0.00 |
| Brain - Hypothalamus | 170 | -0.0153 | 0.7 | 0.00 |
| Brain - Caudate (basal ganglia) | 194 | -0.0348 | 0.3 | 0.00 |
| Brain - Putamen (basal ganglia) | 170 | -0.0408 | 0.3 | 0.00 |
| Brain - Anterior cingulate cortex (BA24) | 147 | -0.0650 | 0.3 | 0.00 |
| Brain - Hippocampus | 165 | -0.0687 | 0.04 | 0.00 |
| Brain - Frontal Cortex (BA9) | 175 | -0.118 | 0.04 | 0.00 |
| Brain - Cortex | 205 | -0.163 | 3.0e-4 | 0.00 |
| Brain - Cerebellar Hemisphere | 175 | -0.168 | 0.01 | 0.00 |
| Cells - EBV-transformed lymphocytes | 147 | -0.183 | 0.05 | 0.00 |
| Brain - Spinal cord (cervical c-1) | 126 | -0.199 | 2.2e-3 | 0.00 |
| Brain - Cerebellum | 209 | -0.335 | 2.4e-7 | 0.00 |

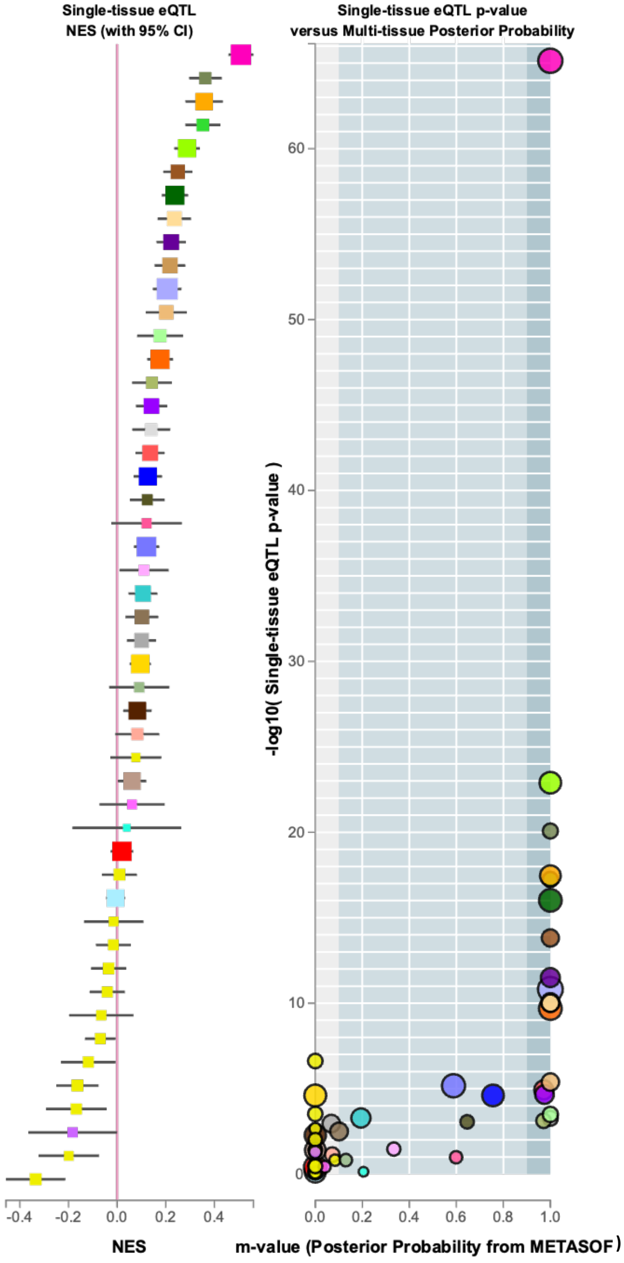

B

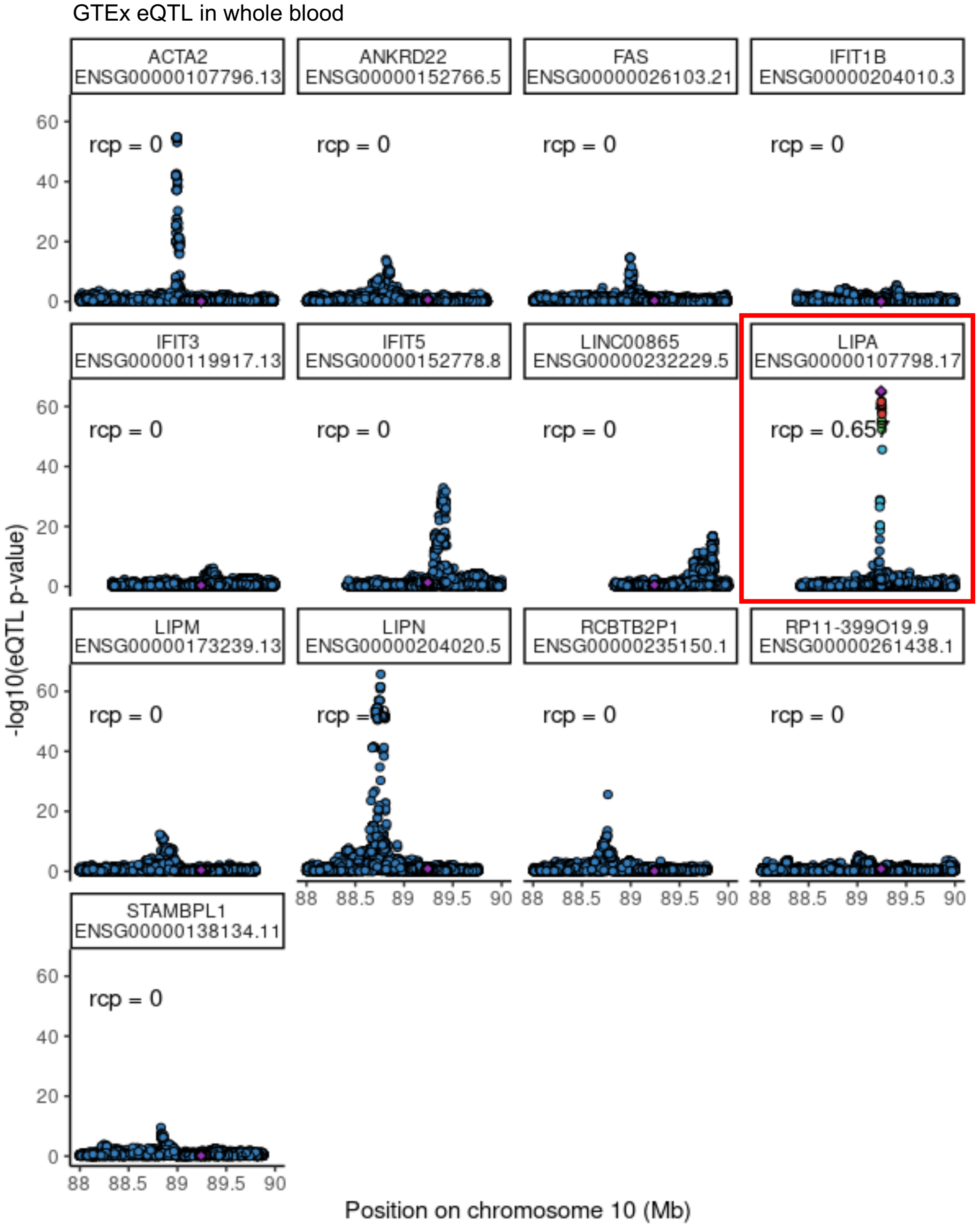

**Figure S1. *LIPA* eQTL and colocalization analysis in GTEx tissues. A,** Expression quantitative trait locus (eQTL) analysis of *LIPA* in Genotype-Tissue Expression project (GTEx) tissues. **B,** Colocalization analysis by integrating CARDIoGRAMplusC4D GWAS dataset with the GTEx eQTL data in blood suggests that *LIPA* is the only nearby gene (± 1 Mb flanking the *LIPA* gene) showing strong regional colocalization probability (rcp = 0.657), indicating *LIPA* as the candidate causal gene at the locus

### Figure S2

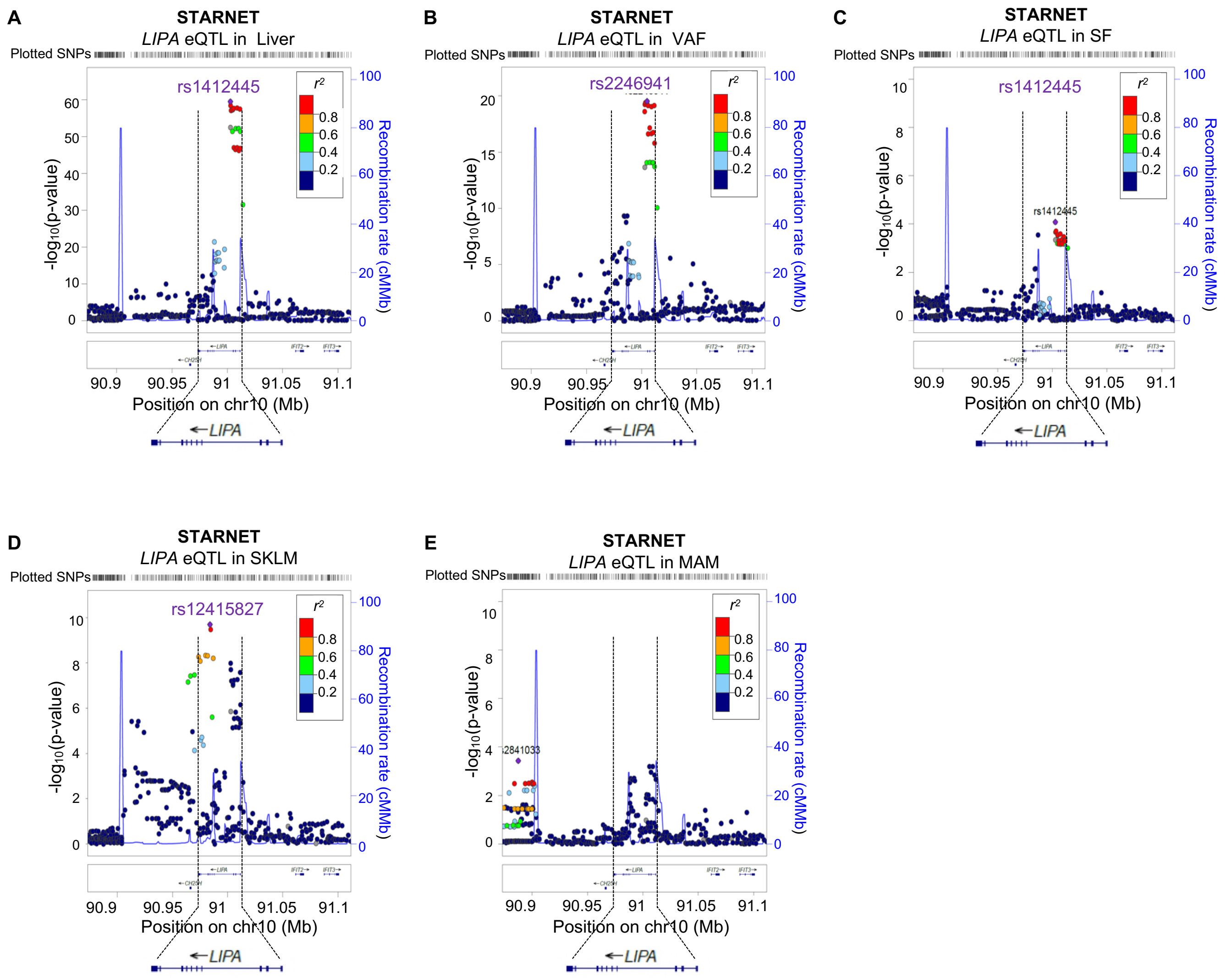

**Figure S2. *LIPA* eQTLs in STARNET tissues.** The Stockholm-Tartu Atherosclerosis Reverse Network Engineering Task (STARNET) dataset allows eQTL analysis in nine cardiometabolic tissues/cell types from individuals with coronary artery disease (CAD). The LocusZoom plots visualize *LIPA* eQTLs in liver (**A**), visceral abdominal fat (VAF, **B**), subcutaneous fat (SF, **C**), skeletal muscle (SKLM, **D**), and lesion-free internal mammary artery (MAM, **E**). The most significant eQTL SNP is marked by a purple diamond. Other SNPs in the region are color-coded by their linkage disequilibrium (LD) with the top SNP according to pairwise $r^2$.

**Figure S3**

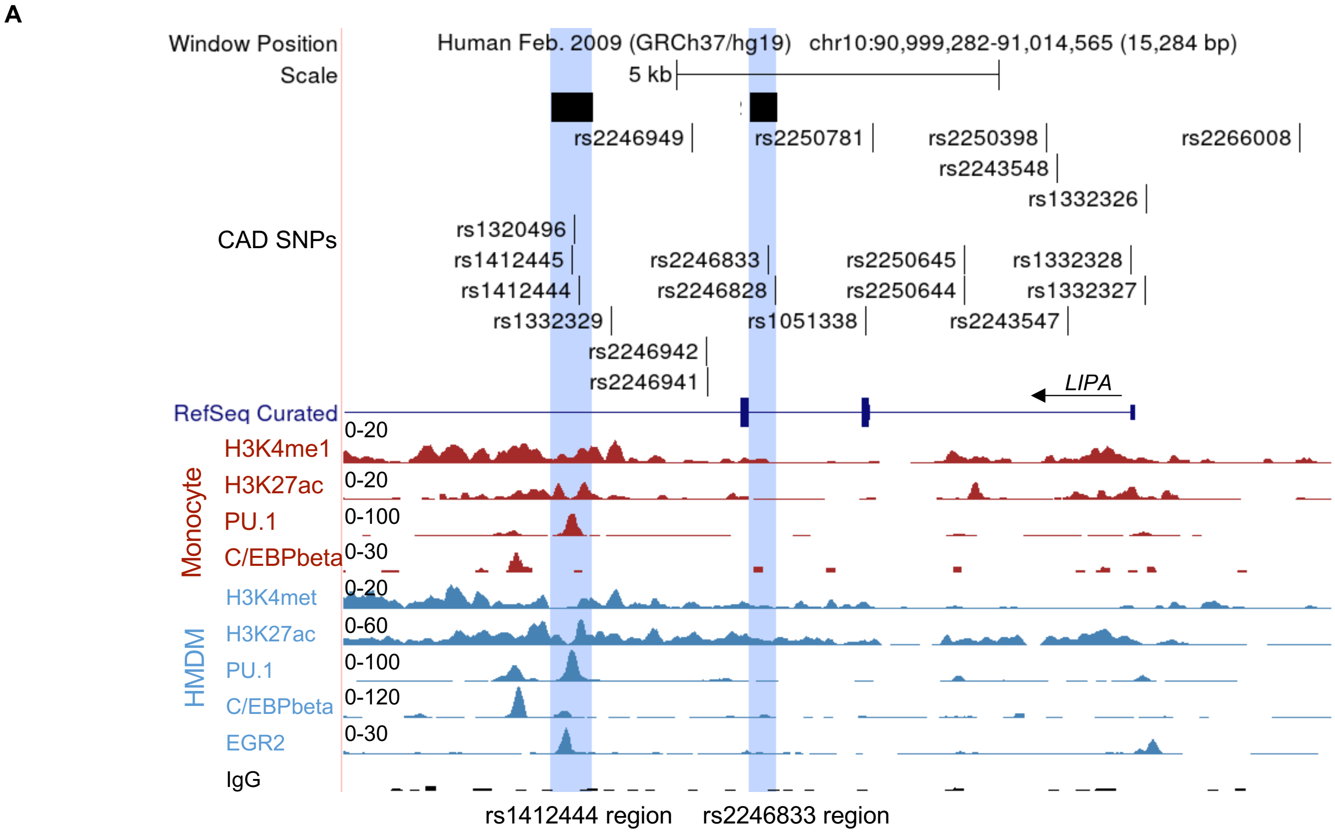

**Figure S3. Genome browser view of the regulatory landscape of the *LIPA* locus in human monocytes and macrophages.** UCSC Genome Browser view of the *LIPA* locus showing all the significant GWAS SNPs for CAD (as listed in **Table S2**), including the lead SNP (rs1412444) and SNPs in high LD, *LIPA* transcripts, and the regulatory landscape in human CD14$^{+}$ monocyte and monocyte-derived macrophages (HMDM). The potential enhancer regions prioritized for functional validation, rs1412444 region and rs2246883 region, are highlighted.

**Figure S4**

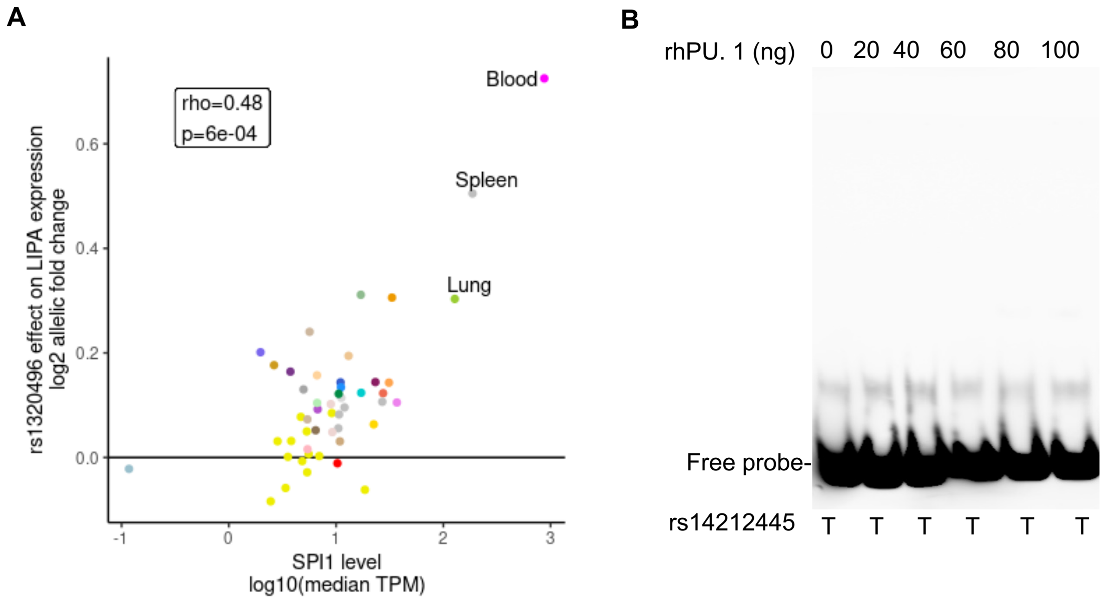

**Figure S4. The SNP rs1412445 is unlikely the functional SNP mediating *LIPA* expression via interaction with transcription factor PU.1. A**, Relationship between rs1320496 *LIPA* eQTL effect size (log2 allelic fold change) and the level of *SPI1* (log10(median transcripts per million)) across GTEx tissues. *LIPA* eQTL effect size increases as the expression of *SPI1* increases (Spearman rho = 0.48, P = 6E-04), suggesting PU.1 (encoded by *SPI1*) binding may regulate the *LIPA* eQTL effect. **B**, Electrophoretic Mobility Gel Shift Assay for the sequences harboring rs1412445 with risk allele (T) using series amount of recombinant human PU.1 protein. Results suggest that the rs1412445 region with the T alleles shows no evidence of PU.1 binding.

**Figure S5**

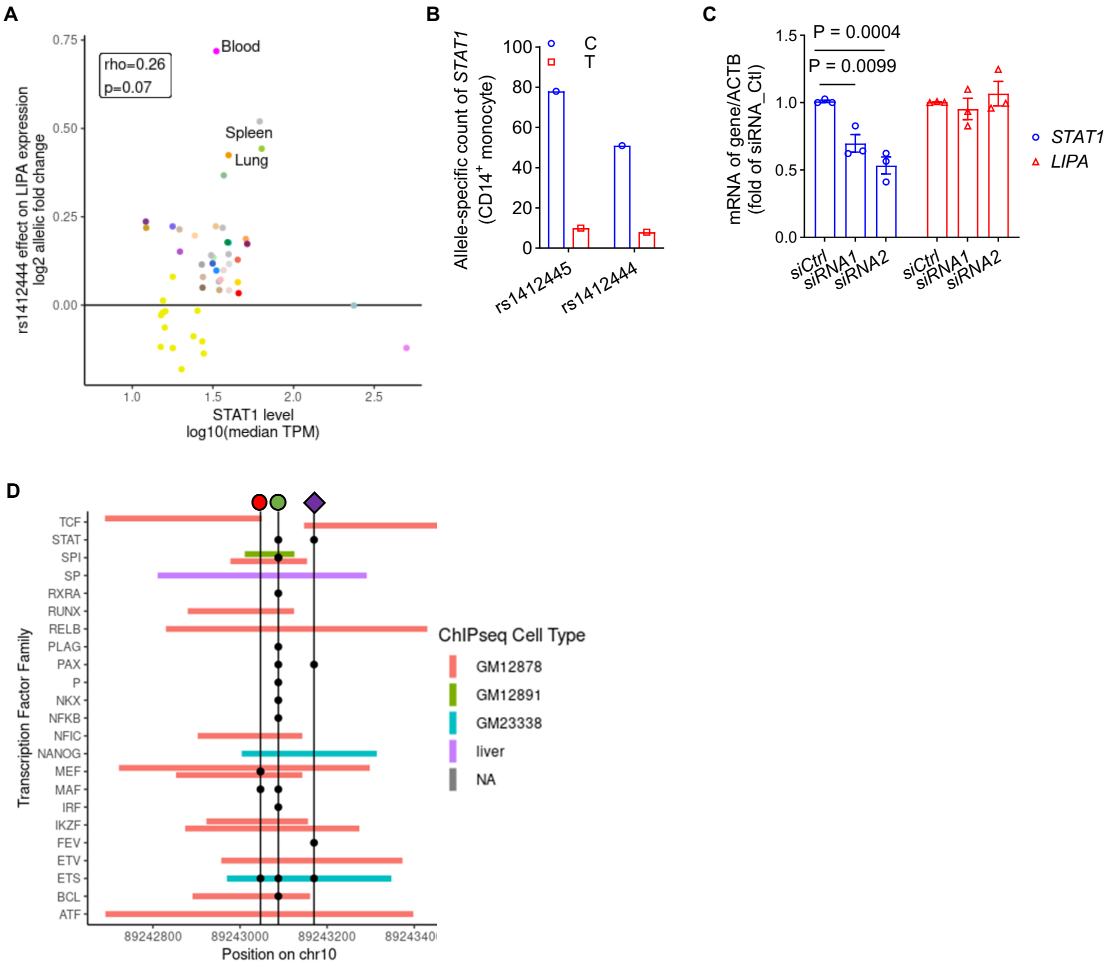

**Figure S5. Transcription factor STAT1 unlikely regulates *LIPA* expression, despite evidence of binding and motif alteration in the CAD risk variant-containing enhancer regions. A**, Correlation between rs1412444 *LIPA* eQTL effect size (log2 allelic fold change) and the level of *STAT1* (log10(median transcripts per million)) across GTEx tissues. *LIPA* eQTL effect size increases as the expression of *STAT1* increases (Spearman rho = 0.26, P = 0.07), suggesting *STAT1* binding may regulate the *LIPA* eQTL effect. **B**, Allele-specific binding (ASB) analysis for *STAT1*. STAT1 ChIP-seq data in CD14$^{+}$ monocytes (n = 1 experiment) in ENCODE dataset indicates ASB for SNPs rs1412445 and rs1412444. Risk alleles (T, red) exhibit reduced STAT1 binding compared to non-risk alleles (C, blue). **C**, siRNA-mediated knockdown of *STAT1* did not alter the expression of *LIPA* in THP-1 monocytes. (n = 3 independent experiments with 2 technical replicates, data are presented as median ± 95%CI). **D**, Transcription factor binding of rs1412444 region. ENCODE transcription factor ChIP-seq peaks (bars) are shown for the ~400 bp flanking region of the SNP rs1320496, while ENCODE transcription factor motif disruptions (black dots) are shown for the three SNPs (rs1412445, red dot; rs1320496, green dot; and rs1412444, purple diamond). Transcription factors are organized by family instead of specific transcription factors due to motif similarity within a family. Many transcription factors have some overlap in this region, SPI and and other three transcription factors families, MEF, ETS, and BCL, have both an overlapping ENCODE ChIP-seq peak and motif.

### Figure S6

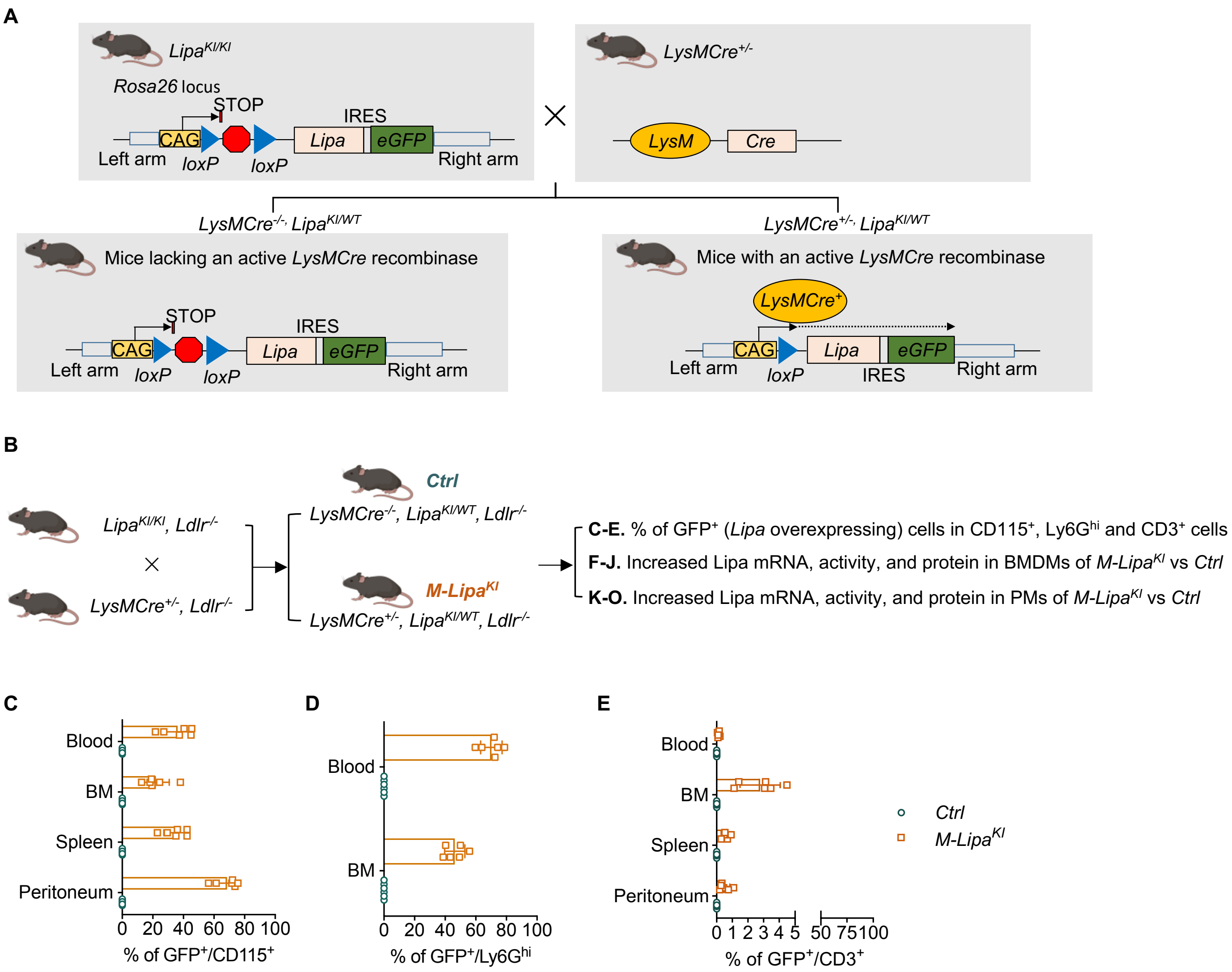

A
LipaKI/KI
Rosa26 locus
STOP
IRES
CAG
Lipa
eGFP
Left arm
loxP
loxP
Right arm
LysMCre+/-
LysM
Cre
LysMCre-/- LipaKI/WT
Mice lacking an active LysMCre recombinase
LysMCre+/- LipaKI/WT
Mice with an active LysMCre recombinase
LysMCre+
B
LipaKI/KI, Ldlr-/-
LysMCre+/-, Ldlr-/-
Ctrl
LysMCre-/-, LipaKI/WT, Ldlr-/-
M-LipaKI
LysMCre+/-, LipaKI/WT, Ldlr-/-
C-E. % of GFP+ (Lipa overexpressing) cells in CD115+, Ly6Ghi and CD3+ cells
F-J. Increased Lipa mRNA, activity, and protein in BMDMs of M-LipaKI vs Ctrl
K-O. Increased Lipa mRNA, activity, and protein in PMs of M-LipaKI vs Ctrl
C
Blood
BM
Spleen
Peritoneum
0 20 40 60 80 100
% of GFP+/CD115+
D
Blood
BM
0 20 40 60 80 100
% of GFP+/Ly6Ghi
E
Blood
BM
Spleen
Peritoneum
0 1 2 3 4 5 50 75 100
% of GFP+/CD3+
Ctrl
M-LipaKI

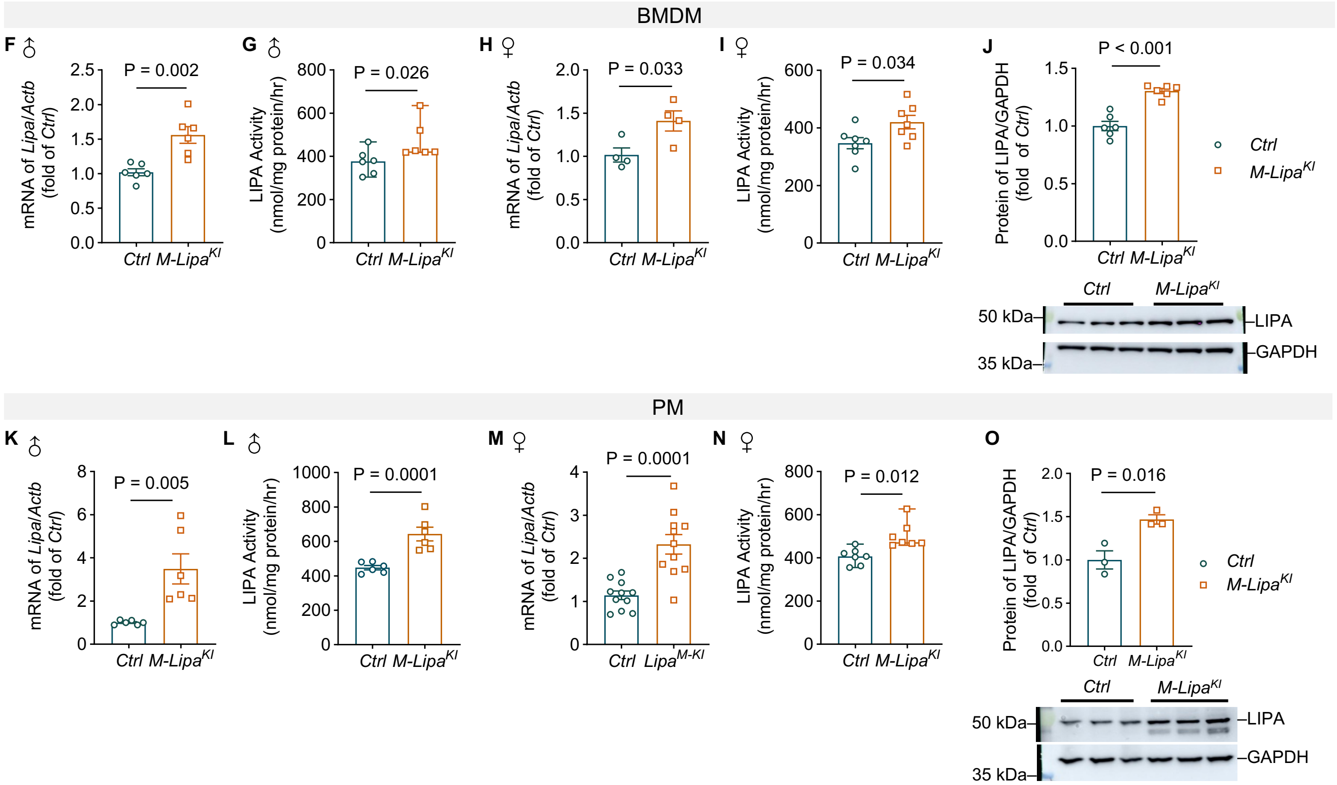

**Figure S6. Generation and validation of *Rosa26* locus *Lipa* knock-in mice for cre-loxp mediated myeloid-specific *Lipa* overexpression. A,** The plasmid construction and breeding strategy for LysMCre-mediated myeloid-specific *Lipa* overexpression. The *Lipa* cDNA is co-expressed with eGFP using an Internal Ribosome Entry Site (IRES). The IRES sequence allows for the independent translation of the cDNA and GFP from a single mRNA transcript. As a result, the expression of GFP serves as a surrogate marker, indicating the successful overexpression of the cDNA. **B**, A summary of strategies validating the myeloid-specific overexpression of *Lipa* at mRNA, protein, and activity levels in both male and female mice. Mice are on an $Ldlr^{-/-}$ background but fed a normal laboratory diet (ND). **C-E**, The percentage of $GFP^+$ (*Lipa*-overexpressing) cells in $CD45^+CD115^+$ monocytes and macrophages (**C**), $CD115^-Ly6G^{hi}$ neutrophils (**D**), and $CD3^+$ lymphocytes (**E**) in the blood, bone borrow (BM), spleen, and peritoneum. (n = 3 female mice and 3 male mice). **F-J,** Increased LIPA at the levels of mRNA (**F,** n = 6 male; **H,** n = 4 female), enzyme activity (**G**, n = 6 male; **I**, n = 7 female), and protein (**J,** n = 6 mice, mixed sexes) in *M-Lipa*$^{KI}$ bone marrow derived macrophages (BMDMs). **K-O**, Increased LIPA at levels of mRNA (**K**, n = 6 male; **M**, n = 11 female), enzyme activity (**L**, n = 6 male; **N**, n = 7 female), and protein (**O,** n = 3 mice, mixed sexes) in *M-Lipa*$^{KI}$ peritoneal macrophages (PMs). Data are presented as mean ± SEM, except for **H, L,** and **O** are presented as median ± 95% CI.

### Figure S7

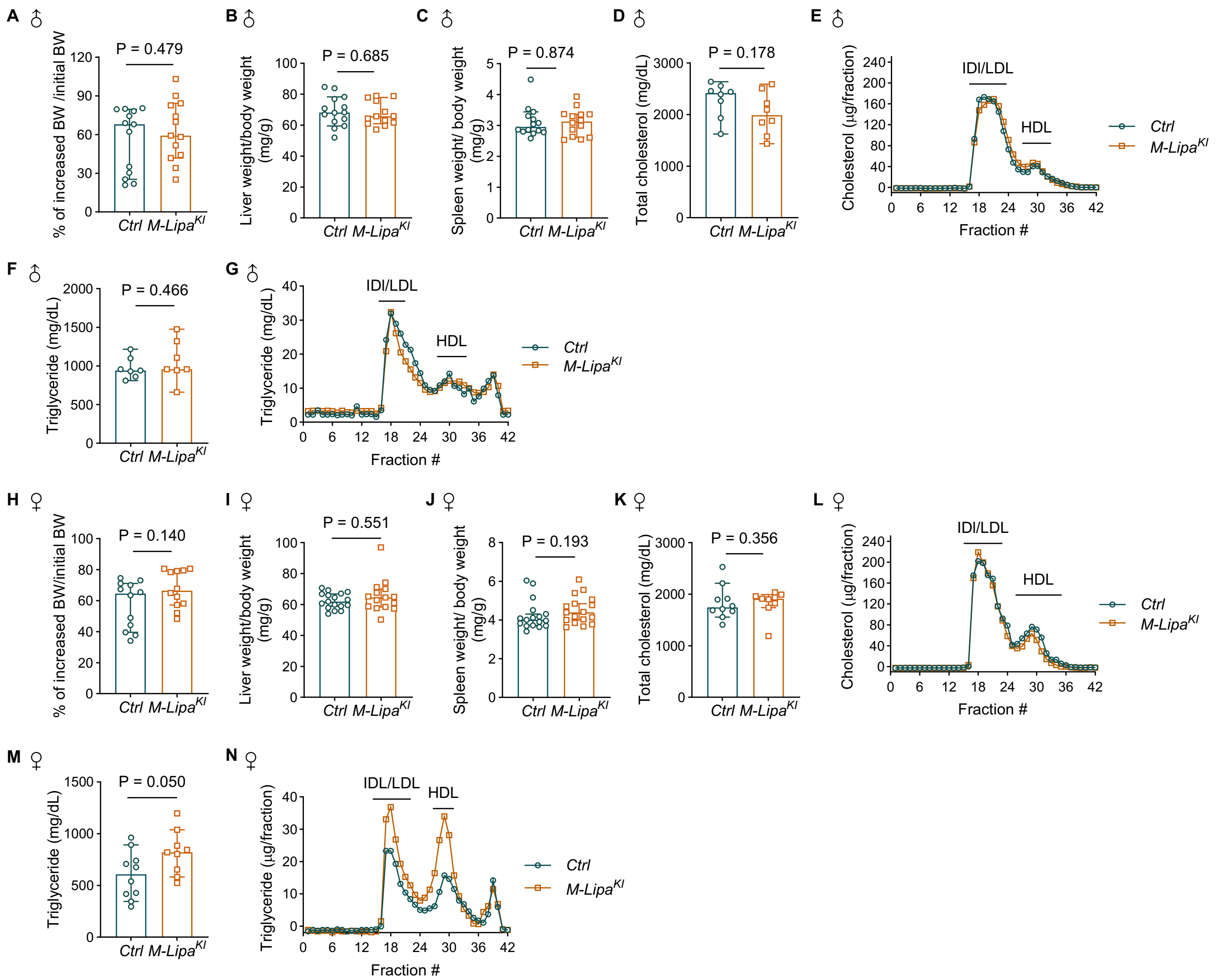

**Figure S7. Myeloid overexpression of *Lipa* did not affect body weight, organ weight, and lipid profile in mice on the atherosclerosis-prone *Ldlr*$^{-/-}$ background and fed a Western diet (WD) for 16 weeks. A-C,** The gained body weight (**A**, n = 13 mice, median ± 95% CI), liver weight/body weight ratio (**B,** n = 14 mice), and spleen weight/body weight (**C,** n = 17 mice) in male mice fed a WD for 16 weeks. **D-E**, The levels of cholesterol in plasma (**D,** n = 8 male mice) and in the fractions of low-density lipoprotein (LDL) and high-density lipoprotein (HDL) (**E,** n = pooled 5 male mice) after fasting for 4 h. **F-G**, The levels of triglyceride in plasma (**F,** n = 7 male mice) and in the fractions of LDL and HDL (**G,** n = pooled 5 male mice) after fasting for 4 h. **H-J,** The body weight (**H**, n = 12 mice, median ± 95% CI), liver weight/body weight ratio (**I**, n = 16 mice), and spleen weight/body weight ratio (**J,** n = 17 mice) in female mice fed a WD for 16 weeks. **K-L,** The levels of total cholesterol in plasma (**K**, *Ctrl*, n = 10 female mice; *M-Lipa*$^{KI}$, n = 9 female mice; median ± 95% CI ) and in fractions of LDL and HDL (**L**, n = pooled 6 female mice) after fasting for 4 h. **M-N,** The levels of triglyceride in plasma (**M**, n = 10 female mice) and in the fractions of LDL and HDL (**N**, n = pooled 6 female mice) after fasting for 4 h. Data are presented as mean ± SEM, except otherwise specified above.

### Figure S8

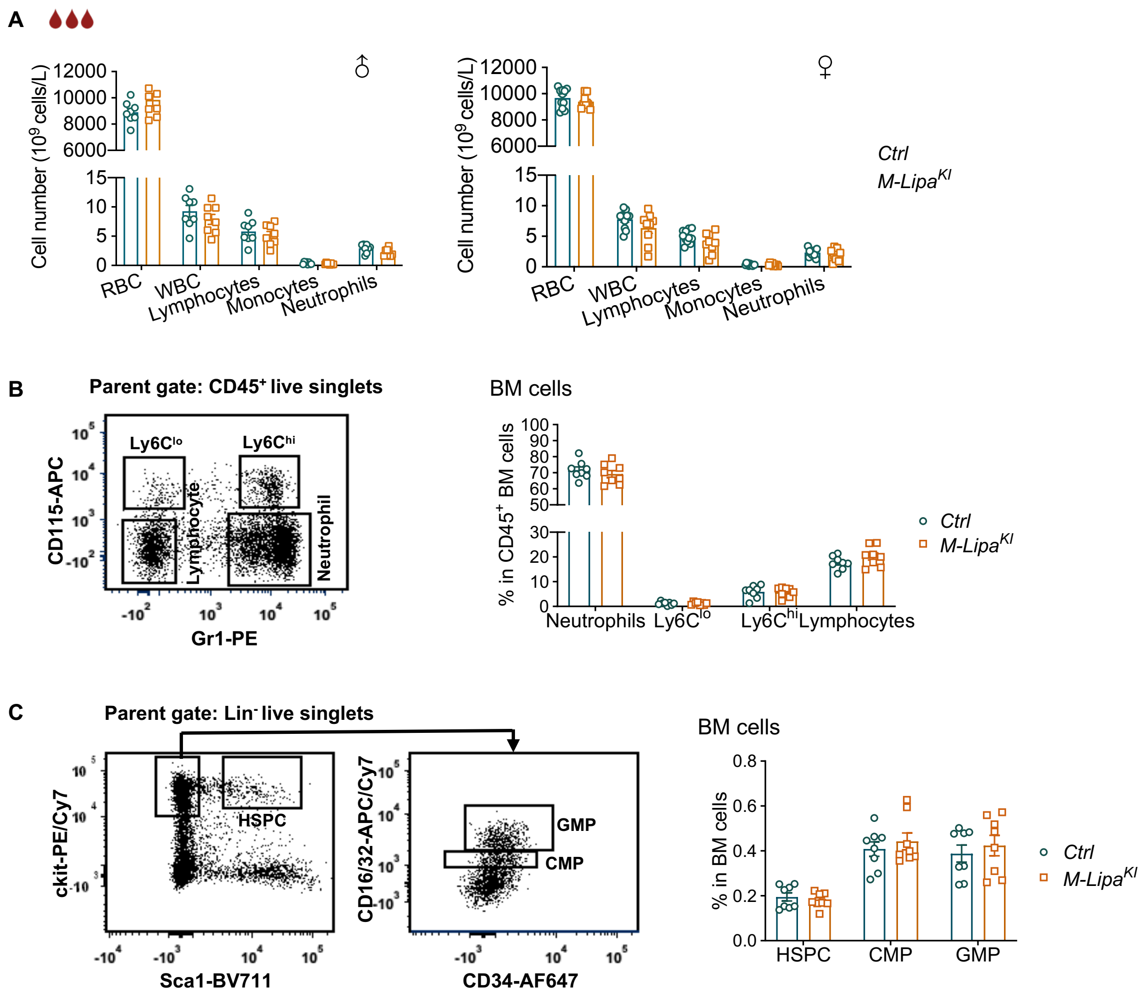

**Figure S8. Myeloid *Lipa* overexpression did not alter blood cell count or hematopoiesis in mice on the atherosclerosis-prone *Ldlr*$^{-/-}$ background and fed a WD for 16 weeks. A,** Cell counts for red blood cells (RBC), white blood cells (WBC), lymphocytes, monocytes, and neutrophils by Complete Blood Count with Differential (left: n = 8 male mice, right: n = 11 female mice for *Ctrl* and n = 9 female mice for *M-Lipa*$^{KI}$). **B,** Gating strategy to identify CD115$^+$Gr1$^{lo}$ and CD115$^+$Gr1$^{hi}$ monocytes, CD115$^-$Gr1$^{hi}$ neutrophils, and CD115$^-$Gr1$^{lo}$ lymphocytes in the BM by flow cytometry. The percentage of Ly6C$^{lo}$ and Ly6C$^{hi}$ monocytes, neutrophils, and lymphocytes in the BM is comparable between *Ctrl* and *M-Lipa*$^{KI}$ mice (n = 2 female mice and 6 male mice). **C**, Gating strategy to identify hematopoietic stem and progenitor cells (HSPC, Lin$^-$cKit$^+$Sca1$^+$), common myeloid progenitor (CMP, Lin$^-$cKit$^+$CD16/32$^{med}$CD34$^{med}$), and granulocyte-monocyte progenitor (GMP, Lin$^-$cKit$^+$CD16/32$^{hi}$CD34$^{hi}$) in the BM by flow cytometry. The percentage of HSPC, CMP, and GMP in the BM is comparable between *Ctrl* and *M-Lipa*$^{KI}$ mice (n = 2 female mice and 6 male mice). Data were represented as mean ± SEM.

### Figure S9

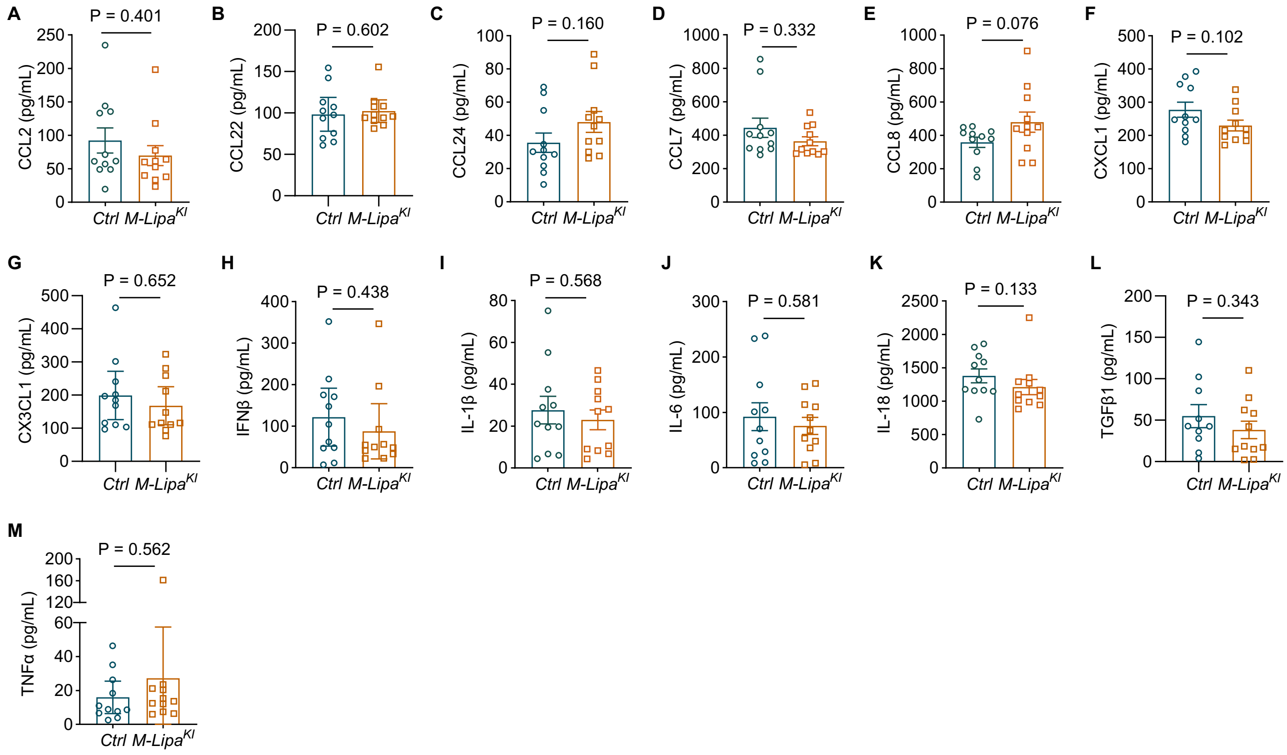

**Figure S9. Myeloid overexpression of *Lipa* did not affect circulating levels of chemokines and cytokines**. Mice were fed a WD for 16 weeks and plasma samples were collected. **A-M,** Plasma CCL2 (**A**), CCL22 (**B**), CCL24 (**C**), CCL7 (**D**), CCL8 (**E**), CXCL1 (**F**), CX3CL1 (**G**), IFNβ (**H**), IL-1β (**I**), IL-6 (**J**), IL-18 (**K**), TGFβ1 (**L**), and TNFα (**M**) were measured by customized LEGENDplex Kit. n = 4 male mice and 7 female mice, using the average of two technical replicates. Data are presented as Mean ± SEM, expect for **B**, **G**, **H**, and **M**, which are presented as median ± 95% CI.

### Figure S10

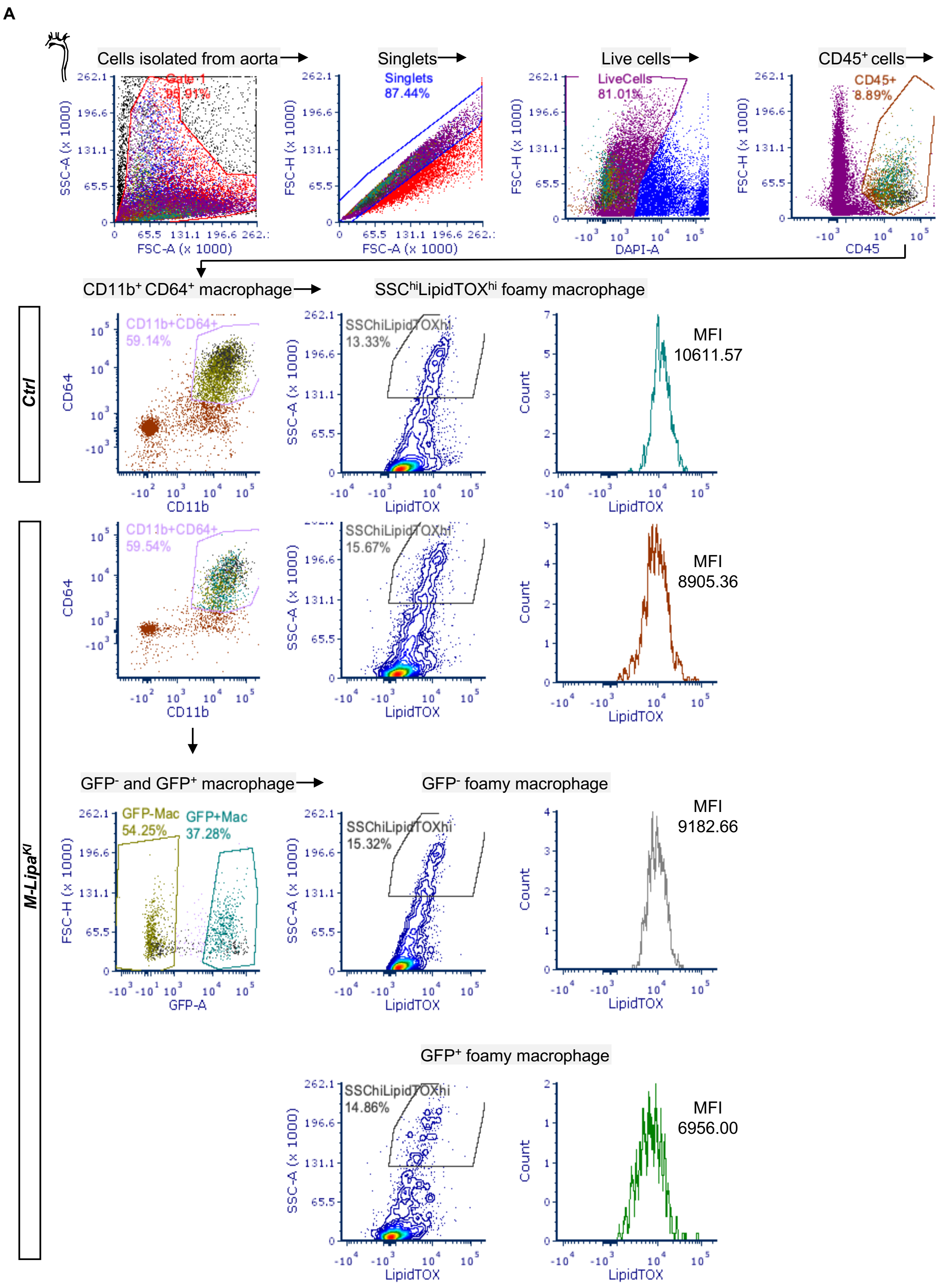
A
Cells isolated from aorta
Singlets
Live cells
CD45⁺ cells
Gate 1
95.91%
Singlets
87.44%
LiveCells
81.01%
CD45+
8.89%
SSC-A (x 1000)
FSC-A (x 1000)
FSC-H (x 1000)
DAPI-A
CD45
CD11b⁺ CD64⁺ macrophage
SSChiLipidTOXhi foamy macrophage
Ctrl
CD11b+CD64+
59.14%
SSChiLipidTOXhi
13.33%
MFI
10611.57
CD64
CD11b
LipidTOX
Count
M-LipaKI
CD11b+CD64+
59.54%
SSChiLipidTOXhi
15.67%
MFI
8905.36
GFP⁻ and GFP⁺ macrophage
GFP⁻ foamy macrophage
GFP-Mac
54.25%
GFP+Mac
37.28%
SSChiLipidTOXhi
15.32%
MFI
9182.66
GFP-A
GFP⁺ foamy macrophage
SSChiLipidTOXhi
14.86%
MFI
6956.00

B

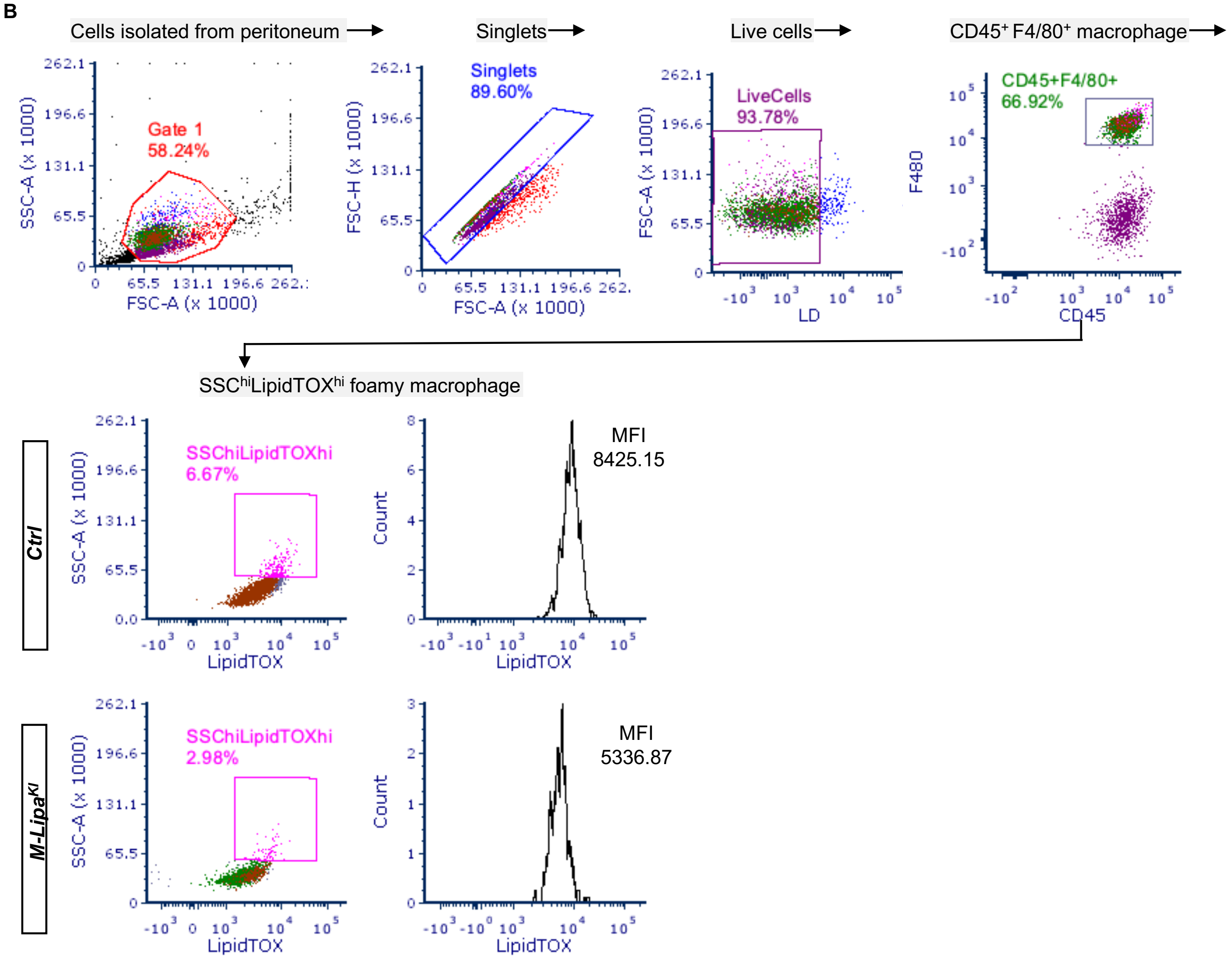

**Figure S10. Gating strategy for aortic macrophages and PMs. A,** Gating strategy for the characterization of foamy cells (SSC$^{hi}$LipidTOX$^{hi}$) in CD45$^{+}$CD11b$^{+}$CD64$^{+}$ aortic macrophages by flow cytometry. **B,** Gating strategy for the characterization of foamy cells (SSC$^{hi}$LipidTOX$^{hi}$) in F4/80$^{+}$ PMs.

### Figure S11

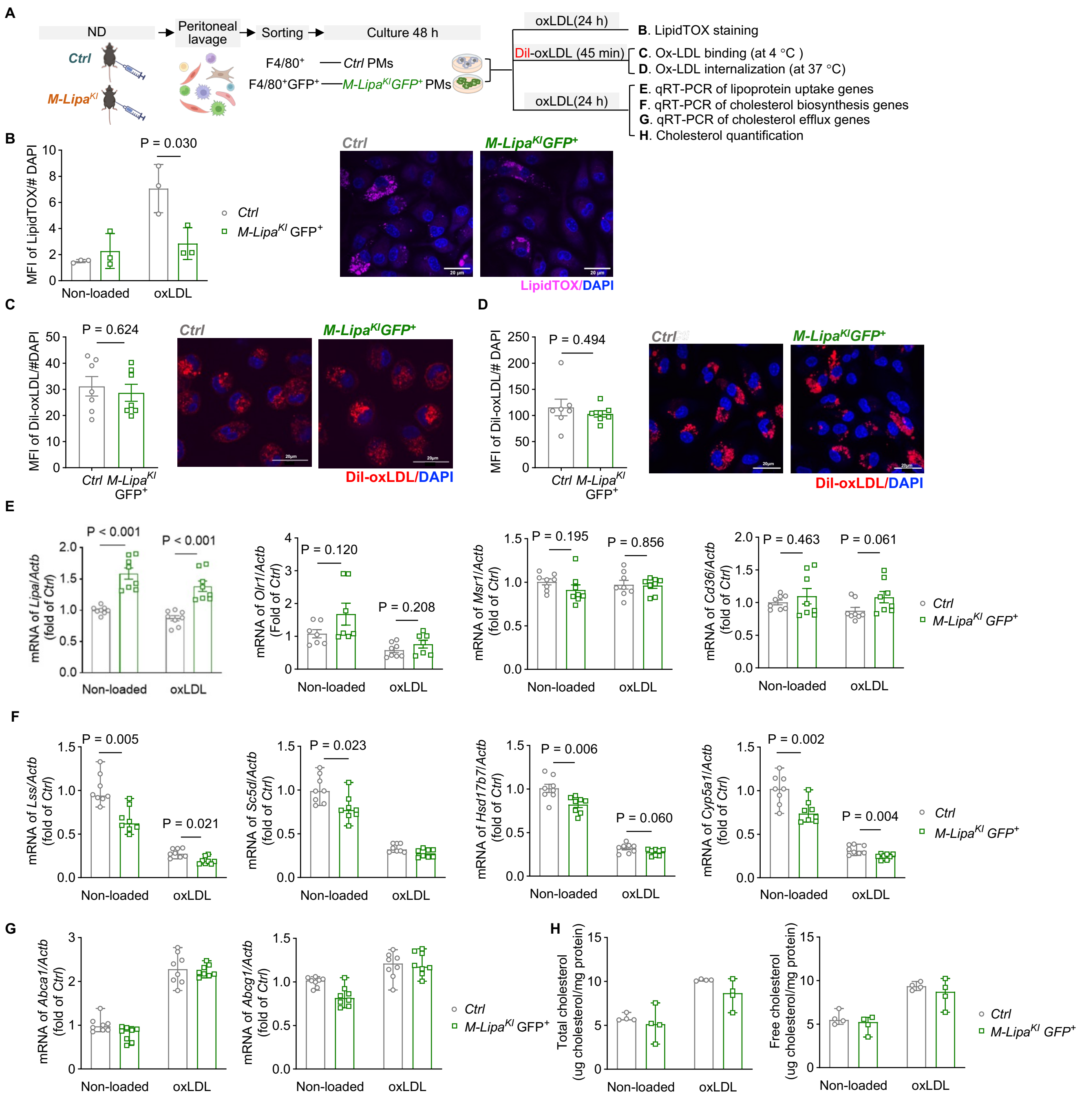

**Figure S11. Characterization of oxLDL loaded PMs in mice fed a normal laboratory diet (ND). A,** Schematic figure for the study design. F4/80$^+$ PMs sorted from the *Ctrl* mice and F4/80$^+$GFP$^+$ PMs sorted from the *M-Lipa$^{KI}$* mice (8-12 weeks old), both fed a normal laboratory diet (ND), were subjected to indicated assays. **B,** MFI of LipidTOX was quantified in sorted PMs incubated with or without 50 µg/mL oxLDL for 24 h. n = 3 male mice. Data are presented as median ± 95% CI. Scale bars = 20 µm. **C-D,** Sorted PMs were loaded with 10 µg/mL Dil-oxLDL for 45 min at 4°C to monitor lipoprotein binding (**C**) and at 37°C to characterize lipoprotein internalization (**D**). n = 3 male mice and 4 female mice. Data are presented as mean ± SEM. Scale bars = 20 µm. **E,** mRNA of *Lipa, Olr1, Msr1,* and *Cd36* in PMs treated with or without 50 μg/mL oxLDL for 24 h. **F**, mRNA of *Lss*, *Sc5d*, *Hsd17b7*, and *Cyp5a1* in PMs treated with or without 50 μg/mL of oxLDL for 24 h. **G**, mRNA of *Abca1* and *Abcg1* in PMs treated with or without 50 μg/mL of oxLDL for 24 h. n = 4 male mice and 4 female mice. Data are presented as mean ± SEM for **E-G**. **H,** Quantification of total cholesterol (left) and free cholesterol (right) by Amplex Red Cholesterol Assay in PMs treated with and without 50 μg/mL of oxLDL for 24 h. n = 4 male mice. Data are presented as median ± 95% CI.

**Figure S12**

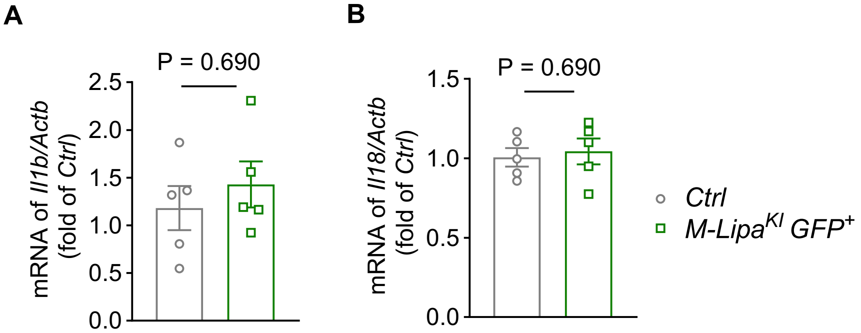

**Figure S12. Myeloid overexpression of *Lipa* did not affect the mRNA expression of *Il1b* and *Il18*.** *Ctrl* and *M-Lipa*$^{KI}$ mice were fed a WD for 16 weeks. The F4/80$^{+}$ PMs from *Ctrl* mice and F4/80$^{+}$GFP$^{+}$ PMs from *M-Lipa*$^{KI}$ mice were collected using a BD Influx Cell Sorter (BD Bioscience). **A-B,** mRNA of *Il1b* (**A**) and *Il18* (**B**) in PMs freshly sorted from *Ctrl* and *M-Lipa*$^{KI}$ fed with 16 weeks of WD. n = 5 female mice, using the average of two technical replicates. Data are presented as median ± 95% CI.

### Figure S13

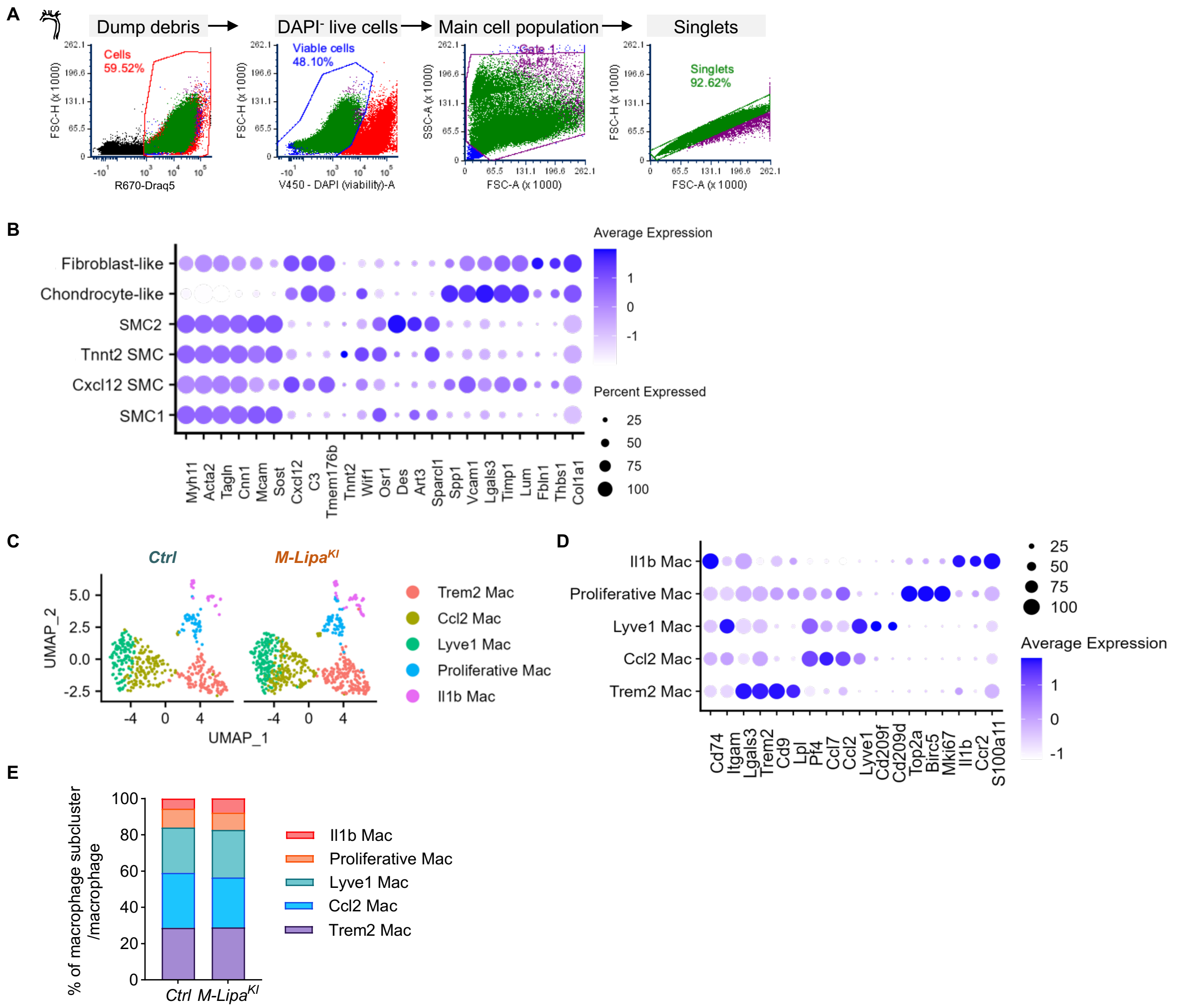

**Figure S13. Subsetting smooth muscle cells and macrophages from scRNA-seq data of all aortic cells for targeting analysis to identify subclusters. A,** Gating strategy for sorting viable single cells from the aortas of *Ctrl* and *M-Lipa*$^{KI}$ mice fed a WD for 16 weeks. **B**, Dot plot visualization of top marker genes of each smooth muscle cell subcluster. **C**, Uniform Manifold Approximation and Projection (UMAP) visualization of macrophage subclusters reveals 5 macrophage subclusters. **D**, Dot plot visualization of top marker genes of each macrophage subcluster. **E**, Stacked bar plot showing the proportion of each macrophage subcluster in *Ctrl* (left) and *M-Lipa*$^{KI}$ (right). Data were analyzed by Chi-square test with Bonferroni correction. **Mac**, macrophages.

### Figure S14

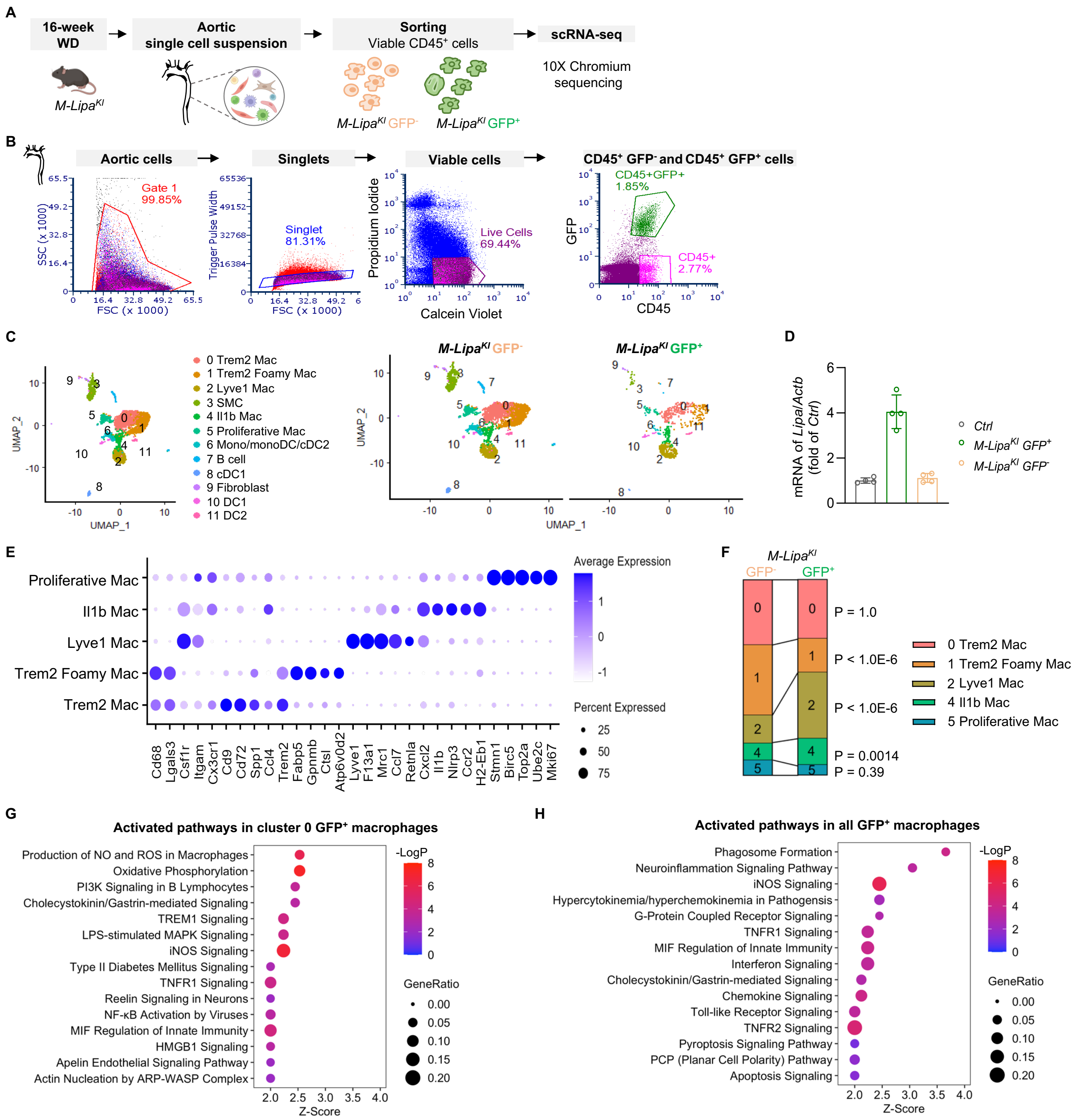

**Figure S14. ScRNA-seq of CD45$^+$ aortic cells isolated from *M-Lipa$^{KI}$* mice fed a WD for 16 weeks. A,** Schematic figure for the study design. Viable CD45$^+$GFP$^-$ and CD45$^+$GFP$^+$ cells isolated from aortas of *M-Lipa$^{KI}$* mice fed a WD for 16 weeks were subjected to scRNA-seq. Cells were obtained from a pooled sample of n = 5 male mice. **B,** Gating strategy for sorting viable CD45$^+$GFP$^-$ and CD45$^+$GFP$^+$ cells from the aortas of *M-Lipa$^{KI}$* mice. **C,** UMAP visualization of clustering reveals 12 cell clusters. **D**, Increased

*Lipa* mRNA in $GFP^+$ aortic macrophages isolated from the *M-Lipa*$^{KI}$ mice compared to aortic macrophages from the *Ctrl* mice and $GFP^-$ aortic macrophages isolated from the *M-Lipa*$^{KI}$ mice ($P < 0.001$, $n = 4$ independent experiments, with each experiment pooled 2-3 mice and has 2 technical replicates). **E**, Dot plot visualization of top marker genes of each macrophage subcluster (clusters 0, 1, 2, 4, and 5). **F**, Stacked bar plot showing the proportion of each macrophage subcluster in $GFP^-$ (left) and $GFP^+$ (right) $CD45^+$ aortic cells isolated from *M-Lipa*$^{KI}$ mice. Data were analyzed by Chi-square test with Bonferroni correction. Mac, macrophages. **G-H,** Top activated canonical pathways by Ingenuity Pathway Analysis (IPA). **G,** The top activated canonical pathways in cluster 0 $GFP^+$ *Lipa*-overexpressing aortic macrophages. **H,** The top activated canonical pathways in all $GFP^+$ *Lipa*-overexpressing macrophages. **Mac**, macrophages.
